# Machine learning-guided spatial omics for tissue-scale discovery of cell-type-specific architectures

**DOI:** 10.64898/2026.02.12.705598

**Authors:** Yumin Lian, Diane-Yayra Adjavon, Takashi Kawase, Jun Kim, Greg Fleishman, Stephan Preibisch, Jan Funke, Zhe J. Liu

## Abstract

Multiplexed protein imaging enables spatially resolved analysis of molecular organization in tissues, but existing spatial proteomics platforms remain constrained in scalability, throughput, and integration with RNA measurements and interpretable computational analysis. Here, we present an integrated spatial omics framework that combines highly multiplexed protein and RNA imaging with explainable machine learning to map cell-type-specific molecular and structural architectures at tissue scale. Using this platform, we simultaneously profiled up to 46 proteins and 79 RNA species across ∼370,000 cells in intact mouse brain tissue at diffraction-limited subcellular resolution (∼260 nm). We developed a scalable, open-source computational pipeline for large-scale image processing and analysis, and show that nuclear protein and chromatin features alone are sufficient to accurately classify brain cell types and their spatial organization. Incorporation of explainable deep learning further enabled identification of human-interpretable, cell-type-specific subnuclear structural features directly from imaging data, with independent quantitative validation. Together, this integrated experimental and computational framework enables tissue-scale spatial proteomics-based cell-type classification and structural feature discovery, providing a broadly applicable platform for mechanistic studies, high-content screening, and translational applications.

## Introduction

Biological systems are composed of interconnected molecular components that are spatially and temporally organized to perform complex functions. A central challenge in modern biology is to quantify and understand how these molecular components (*e.g.*, RNAs, proteins, and their complexes) are arranged within cells and tissues to support diverse biological states. This requires technologies that can simultaneously measure multiple classes of biomolecules and map their spatial relationships in intact tissues. High-dimensional, spatially resolved omics approaches are well suited to this task, offering the ability to link gene expression programs, subcellular organization, and cell-cell interactions across tissue architectures.

Recent years have witnessed rapid progress in spatial transcriptomics and proteomics. On one end of the spectrum, highly multiplexed RNA and DNA imaging methods such as MERFISH^1^, seqFISH^2^, and STARmap^3^ enable the mapping of thousands of RNA species within intact tissues using barcode- or sequencing-based cross-round demixing. In parallel, imaging-based spatial proteomics platforms such as CODEX^4^ and sequential immunostaining platforms^5–7^ provide subcellular localization of tens of protein targets through antibody based iterative labeling and imaging. However, CODEX lacks signal amplification and is therefore less suited for detecting low-abundance proteins, while sequential immunostaining platforms, despite using secondary antibody amplification, lacks a unified barcoding framework for simultaneous, high-throughput measurement of both RNA and proteins in the same sample.

Mass spectrometry-based spatial proteomics provides a complementary approach, combining laser microdissection with protein identification to quantify thousands of proteins per cell, but it lacks subcellular spatial resolution^8^. At the opposite end, super-resolution imaging techniques such as FLASH-PAINT^9^ and SUM-PAINT^10^ achieve nanometer-scale localization of ∼30 protein targets, enabling the visualization of molecular interactions at near-molecular precision. Yet, these techniques rely on high-numerical-aperture, short–working-distance objectives and single-molecule localization accumulation, which restrict field of view, imaging depth, and throughput. Consequently, they are not easily applicable to comprehensive, tissue-scale studies spanning tens of thousands of cells.

These limitations underscore the pressing need for a platform that integrates high multiplexing, deep imaging, and scalable throughput while jointly profiling RNA and proteins in intact tissues with high subcellular resolution. To meet this need, we developed cycleHCR^11^, a multi-omics imaging platform that combines RNA and protein barcoding within a unified framework. The original system encodes up to 2,700 RNA targets and 192 antibodies using a single-shot-per-target strategy coupled with hybridization chain reaction (HCR)^12, 13^-based signal amplification. This built-in amplification provides strong signal gain, enabling reliable detection of low-abundance targets deep within tissues while boosting photon yield for rapid, high-throughput volumetric imaging (Fig. 1). As a result, the fully automated cycleHCR system enables simultaneous RNA and protein imaging across tens of thousands of cells in intact tissues.

**Fig. 1.**
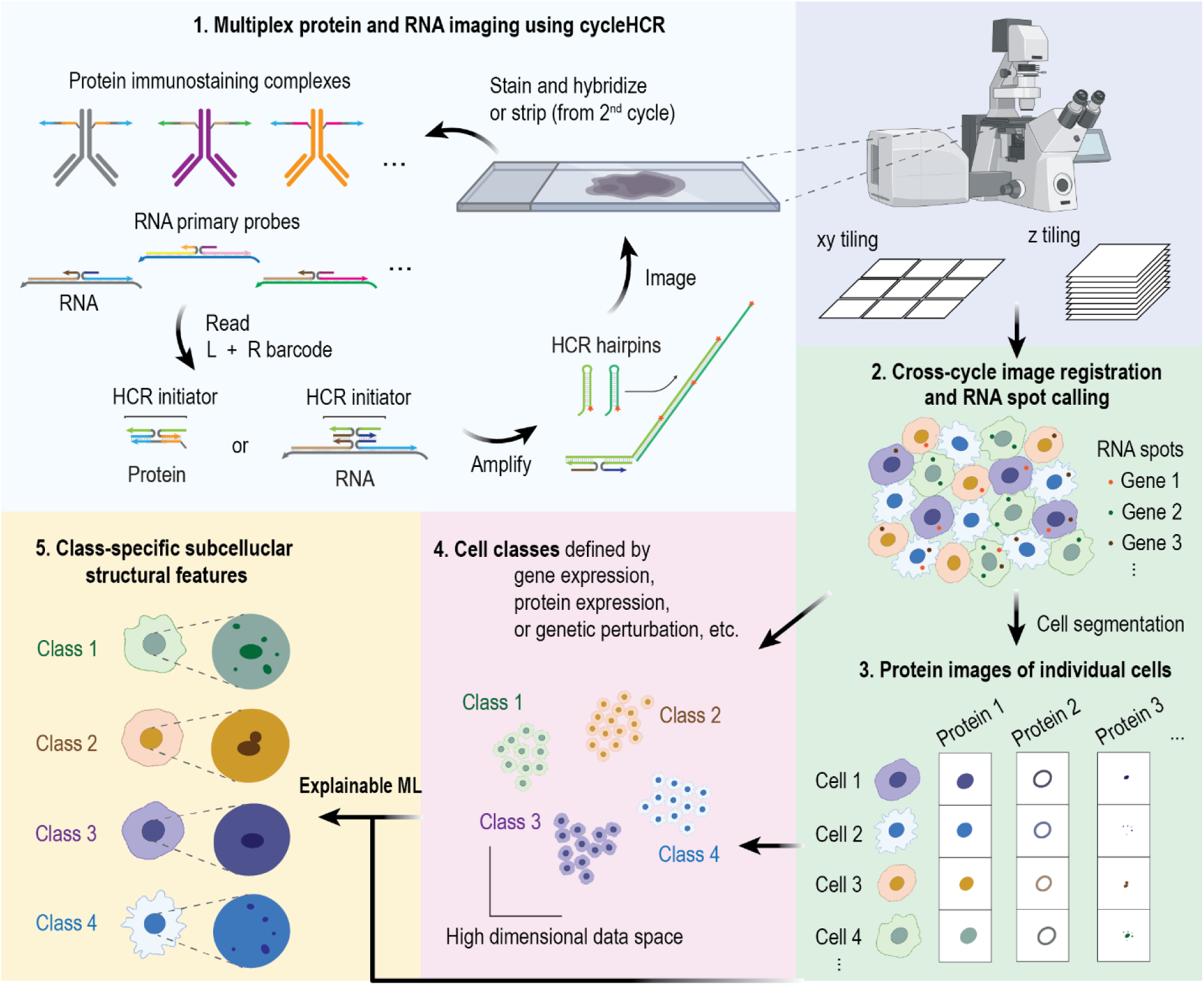
Machine-learning-guided cycleHCR spatial omics reveals cell-type-specific subcellular structures. Step 1: Tissue samples are labeled with barcoded antibody and RNA probe pools, each carrying unique left (L) and right (R) barcodes. Protein and RNA targets are imaged sequentially through cycles of L+R readout hybridization, HCR amplification, and chemical stripping using an automated fluidics-microscope system. Step 2: Volumetric image stacks are stitched and registered across imaging cycles, followed by RNA spot detection. Step 3: 3D cell images are segmented for RNA spot-to-cell assignment and extraction of single-cell protein images. Step 4: Cells are classified using multi-omics measurements or potentially known genetic perturbations. Step 5: Explainable machine-learning methods identify cell-type-specific subcellular structural features directly from protein images.

In parallel, large-scale monoclonal antibody libraries developed through consortium and commercial efforts have generated more than ten thousand validated antibodies^14^. To fully leverage these resources, it is essential to expand the cycleHCR antibody barcoding system to a comparable scale, capable of encoding thousands of antibodies while maintaining high molecular specificity. Such an expanded framework would eliminate the need for secondary antibodies, avoiding cross-reactivity among antibodies from the same species. This system would also allow multiplex imaging of arbitrary combinations of encoded antibodies, greatly enhancing scalability and flexibility for spatial proteomics imaging.

In addition, previous computational pipelines were limited in their ability to extract the high-resolution subcellular information provided by cycleHCR. Here, we substantially advance this analysis framework by implementing robust and scalable single-cell segmentation, quantitative nuclear protein profiling, image-based cell-type classification, and explainable machine-learning that identify human-interpretable and quantifiable cell-type-specific structural features (Fig. 1).

## Results

### Expanded and highly specific cycleHCR barcodes for antibodies

The cycleHCR system enables highly multiplexed protein imaging by tagging antibodies with unique DNA barcodes (Fig. 2a). In the first step, a DNA docking strand is covalently conjugated to the antibody through a light-activated photocrosslinking reaction^15^. A second DNA strand, termed the gel-anchoring probe, then hybridizes to this docking strand. The probe carries paired left (L) and right (R) barcode sequences and a reactive group that covalently incorporates the barcode into the surrounding hydrogel matrix. Once anchored, the barcode remains permanently embedded even after antibody removal by proteinase K treatment, allowing the tissue to be cleared and repeatedly re-imaged without signal loss.

**Fig. 2.**
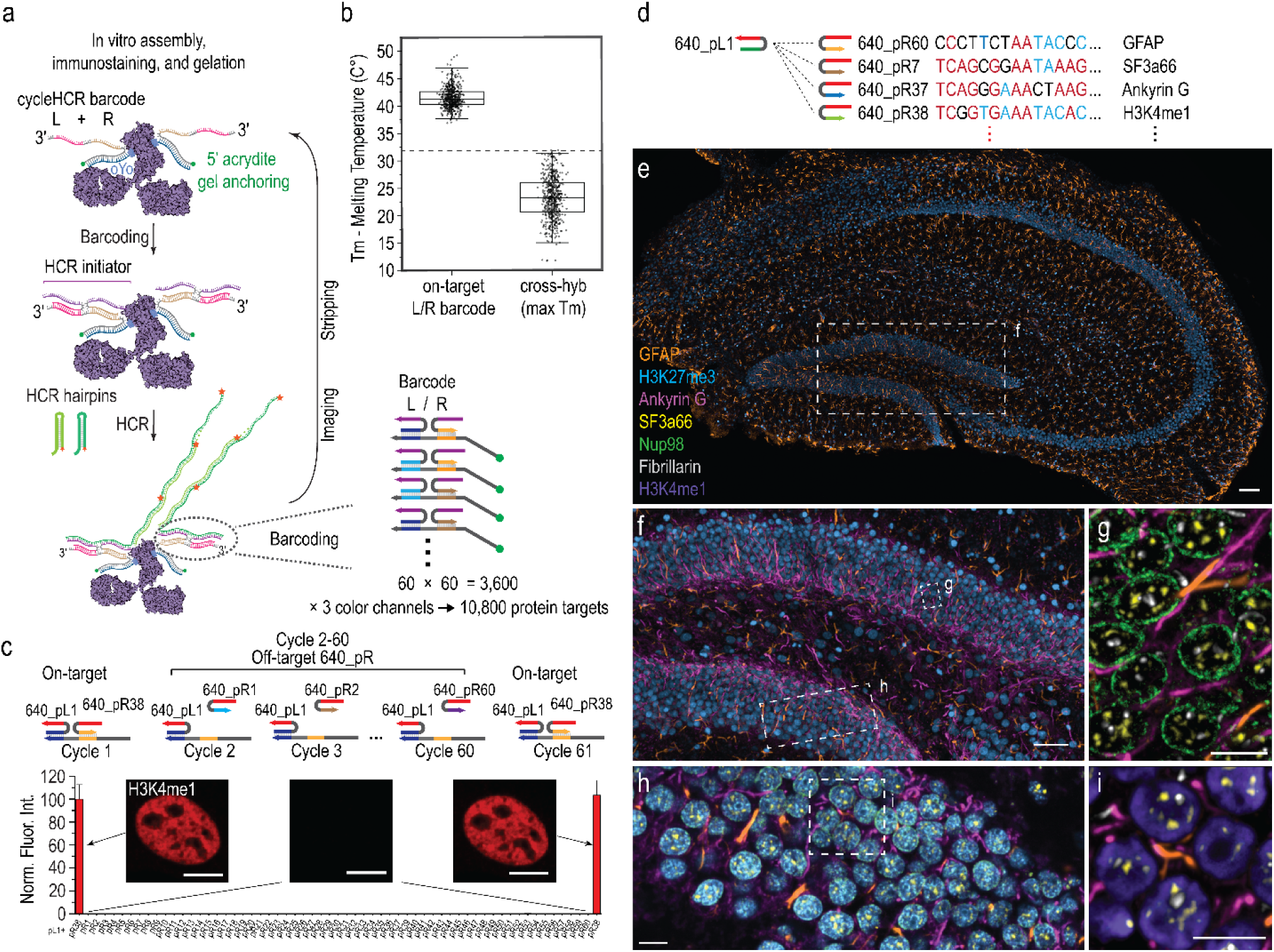
Scalable cycleHCR barcodes for multiplexed antibody imaging. **a,** Schematic of the cycleHCR barcoding strategy. Orthogonal left (L) and right (R) barcode pairs enable up to 10,800 uniquely addressable protein targets. Antibodies are conjugated to oYo-linked DNA oligos and hybridized to 5′ acrydite-modified gel anchoring probes containing L + R barcode sequences. Samples are embedded in polyacrylamide, covalently incorporating the anchoring probes. In each imaging cycle, matching L + R probes and an HCR initiator trigger hybridization chain reaction (HCR) with fluorophore-labeled hairpins. After imaging, barcodes and HCR probes are stripped before the next cycle. Arrowhead indicates the 3′ DNA end. **b,** Melting temperatures (Tm) of 540 barcode probes for proteins [(60 L + 60 R) × 3 color channels = 360 probes] and RNAs [(30 L + 30 R) × 3 channels = 180 probes]. The maximum cross-hybridization Tm is calculated among probes sharing the same split-initiator sequence. The dashed line indicates the 32 °C hybridization temperature. **c,** Barcode cross-reactivity test using H3K4me1 labeling (640 nm channel) in NIH/3T3 cells. Cycle 1 used on-target barcodes; cycles 2–60 reused the same L barcode while iterating through the other 59 R barcodes. The final cycle repeated the on-target pair. Mean nuclear fluorescence (n = 575 cells, 8 fields) is shown ± 95% CI. Scale bar, 10 μm. **d,** Multiplexed protein imaging in a 40-μm mouse hippocampus section using 7 antibody targets encoded with a shared left (L) barcode and distinct right (R) barcodes with the highest sequence similarity (Extended Data Fig. 2). Red and blue nucleotides indicate sequences shared across the listed barcodes. **e,** Astrocyte marker GFAP and heterochromatin marker H3K27me3 in the full hippocampus. Scale bar, 100 μm. **f,** Enlarged region from (e) showing GFAP, H3K27me3, and Ankyrin G (axon initial segment); 3D views in (e,f) rendered in Imaris. Scale bar, 50 μm. **g,** Single-z plane of boxed region in (f) showing GFAP, Ankyrin G, SF3a66 (spliceosome), Nup98 (nuclear pore), and fibrillarin (nucleolar fibrillar center). Scale bar, 10 μm. **h,** Zoomed view from (f) showing GFAP, Ankyrin G, SF3a66, Nup98, and H3K27me3. Scale bar, 10 μm. **i,** Further zoom-in from (h) showing GFAP, Ankyrin G, H3K4me1, SF3a66, and fibrillarin. Scale bar, 10 μm.

To expand the barcoding capacity of cycleHCR for antibody imaging, we built upon our previous RNA barcoding framework (30 × 30 barcodes)^11^ while maintaining strict orthogonality between the RNA and protein modules. Using stringent DNA hybridization design parameters (see Methods), we generated 60 unique left and 60 unique right barcode sequences across three fluorescence channels. Each probe was designed with an average on-target melting temperature of ∼42 °C, while the maximum cross-hybridization melting temperature was constrained below 32 °C - the temperature used in all fluidic and imaging steps - to ensure high specificity and minimal cross-reactivity (Fig. 2b). This design expands the theoretical barcoding capacity up to 10,800 unique antibodies (60 × 60 × 3 channels).

To validate the specificity of the expanded antibody barcoding system, we first tested it in a controlled setting by fixing the left barcode and cycling through all possible right barcode combinations for three highly abundant nuclear proteins across 3 color channels (Fig. 2c and Extended Data Fig. 1). As predicted, we observed negligible cross-reactivity, and fluorescence signals could be reliably and robustly re-detected even after 60 rounds of imaging. To further validate the specificity of our system in tissue, we fixed the left barcode in the 640 nm channel and deliberately selected seven right barcodes with the highest sequence similarity scores (Fig. 2d and Extended Data Fig. 2). These barcodes were conjugated to antibodies targeting GFAP, H3K27me3, Ankyrin G, SFa66, Nup98, Fibrillarin, and H3K4me1, which display well-characterized, non-overlapping spatial patterns in the hippocampal brain slice. As expected, the observed spatial distributions of these seven antibody targets, imaged across seven sequential rounds, closely matched their known biological localizations, with minimal spatial crosstalk between imaging rounds (Fig. 2e–h and Extended Data Fig. 3–4). Together, these results demonstrate that the expanded protein barcoding system performs as designed with high molecular specificity.

### Hierarchical labeling enables cost-efficient antibody multiplexing

A key challenge in scaling cycleHCR protein imaging lies in the cost of synthesizing unique photocrosslinking docking strands (oYo linkers)^15^ for every antibody. To overcome this, we designed a hierarchical two-part labeling strategy that decouples the docking strand (photocrosslinking linker) from the cycleHCR barcode probe carrying the left–right barcode pair and gel-anchoring group (Fig. 2a, 3a).

**Fig. 3.**
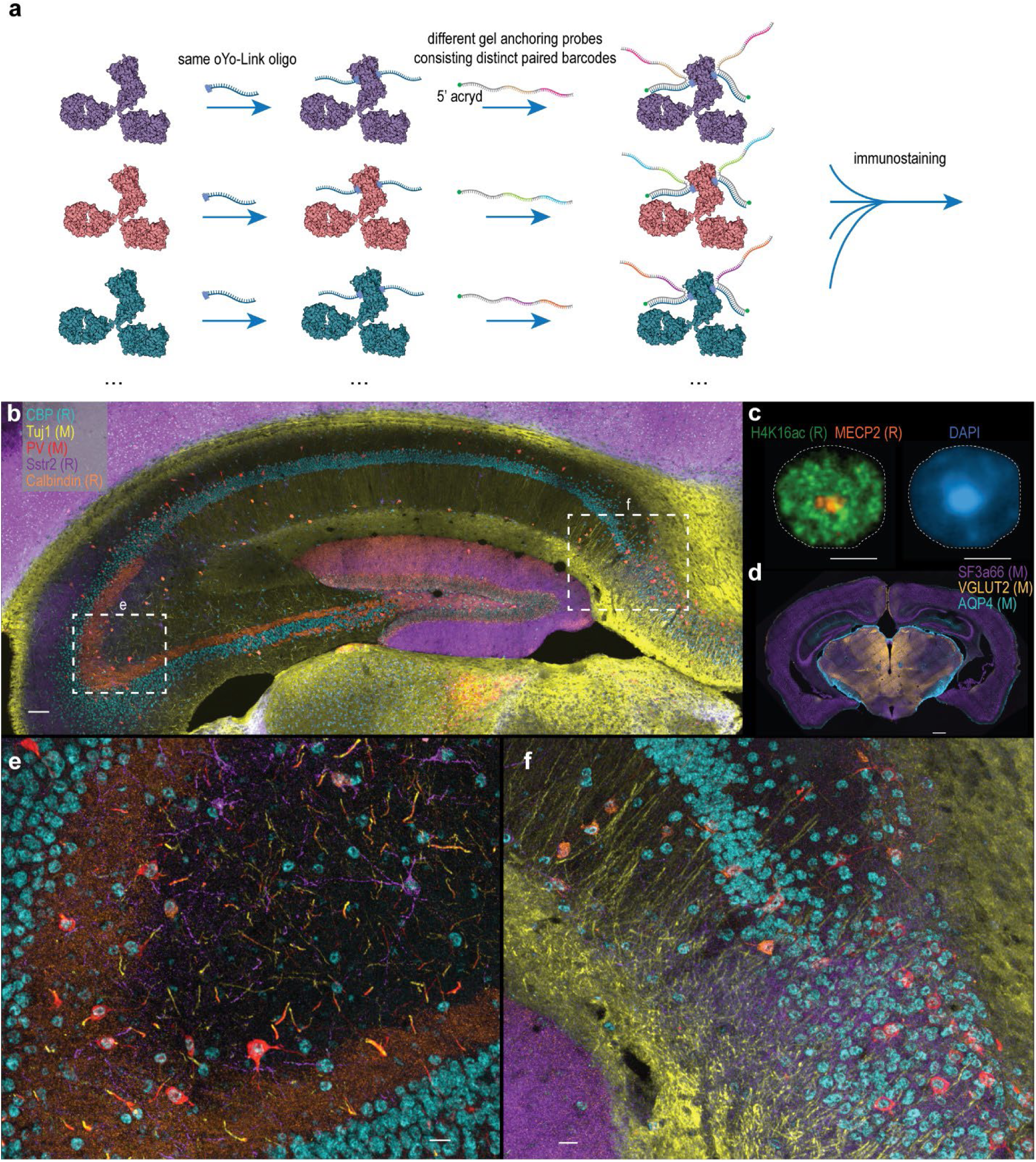
Modular barcoding strategy for cost-efficient multiplex antibody labeling. **a,** Schematic of the two-part labeling workflow. Multiple antibodies are photo-crosslinked to the same oYo-Link oligo, then individually hybridized with distinct barcode probes carrying unique L + R barcode pairs. The barcoded antibody complexes are pooled for multiplex immunostaining. **b–f,** Distinct protein patterns visualized by cycleHCR in a mouse brain section using antibody complexes conjugated with the same oYo-Link oligo. Antibodies from the same host species were used simultaneously without cross-reactivity (host species indicated for each antibody target in the figure: R, rabbit; M, mouse). Orthogonal 3D rendering performed in Imaris. **b,** Five-color composite showing CREB-binding protein (CBP), neuron-specific βIII-tubulin (Tuj1), parvalbumin (PV), cannabinoid receptor 1 (CB1), somatostatin receptor 2 (Sstr2), and calbindin in the hippocampal region of a mouse brain section. These targets belong to a subset of 11 antibodies crosslinked to the same oYo-Link sequence (No. 2; Supplementary Table 1) within the complete 46-protein panel. Scale bar, 100 μm. **c,** (Left) 3D composite of two nuclear proteins, H4K16ac and MECP2, from the same oYo-Link (No. 2) set. (Right) Corresponding DAPI channel of the same nucleus. Scale bars, 5 μm. **d,** Three-color composite of SF3a66, AQP4, and VGLUT2 from another oYo-Link (No. 1) set (four antibodies) within the 46-target panel. Scale bar, 500 μm. **e,f,** Higher-magnification views of boxed regions in (b), showing fine subcellular localization and structural detail. Scale bars (e, f), 20 μm.

Because each antibody-linker complex is assembled independently *in vitro*, we examined whether multiple antibodies could share a common docking strand without compromising specificity (Fig. 3a). To minimize potential cross-talk among antibodies labeled with the same docking sequence, we used an excess of barcoding probes relative to docking strands, ensuring complete and saturated pairing of docking–anchoring complexes prior to antibody mixing and staining. Under these conditions, we observed minimal signal cross-talk, even for highly abundant targets such as the cytoskeletal protein Tuj1 and abundant nuclear proteins including MECP2 and H4K16ac in brain tissue (Fig. 3b-c). Using this system, antibodies derived from the same host species could also be encoded using identical docking strands without detectable interference (Fig. 3b–f). This two-part architecture greatly reduces reagent cost and preparation complexity. Only the cycleHCR barcode probe (costing approximately $100 per antibody) requires customization, whereas the photocrosslinking docking chemistry can be shared across multiple antibodies. Together, this design provides a cost-effective and scalable barcoding framework that maintains high specificity and potentially enables large-scale integration of antibody libraries for multiplexed spatial omics imaging.

### Integrated RNA-protein multi-omics imaging in tissues

To demonstrate the capabilities of the expanded cycleHCR imaging platform, we performed multiplexed protein and RNA imaging on 40-60 µm coronal sections of whole mouse brain tissue slices. Following stringent antibody selection and validation (See Methods for details), the brain section was stained with 46 cycleHCR-barcoded antibodies (Fig. 4a–d, Extended Data Fig. 5–8, Supplementary Table 1) and hybridized with cycleHCR RNA probe libraries targeting 79 cell type-specific genes (Supplementary Table 2). Automated imaging of all targets was conducted using our automated fluidic and imaging system^11^, which implements the unified left + right barcode decoding scheme for both protein and RNA targets.

**Fig. 4.**
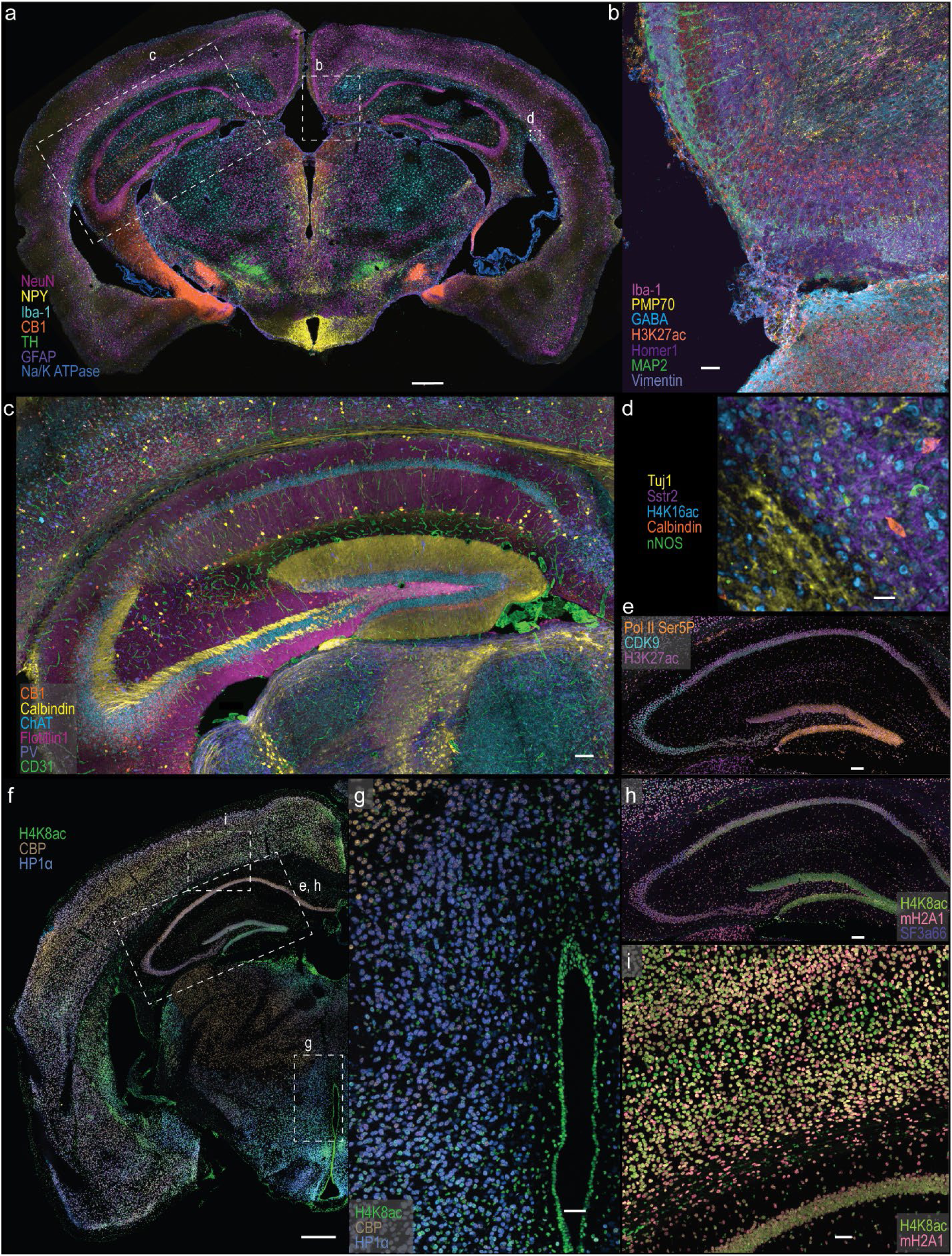
Unified multiplex RNA and protein imaging in mouse brain. **a–d,** Multiplexed protein imaging in a 40-μm mouse brain section (Brain Section 1) using the unified RNA–protein cycleHCR barcoding scheme. In total, 46 proteins and 79 RNA species were imaged within the same sample. **a,** Seven-color composite showing NeuN, NPY, Iba-1, CB1, TH, GFAP, and Na/K ATPase across the whole brain section. Scale bar, 500 μm. **b–d,** Enlarged cortical and hippocampal regions showing distinct subsets of proteins, such as glial, synaptic, cytoskeletal, and chromatin modification markers (e.g., Iba-1, Homer1, MAP2, Vimentin, H3K27ac), highlighting the platform’s ability to resolve diverse molecular architectures within intact tissue. Scale bars: (b) 50 μm, (c) 100 μm, (d) 20 μm. **e–i,** Nuclear protein distributions in a separate 40-μm mouse brain section (Brain Section 2) imaged for 16 proteins and 79 RNA targets. **e,** Three-color composite of Pol II Ser5P, CDK9, and H3K27ac in the hippocampus showing spatially distinct expression patterns. Scale bar, 100 μm. **f,** Composite of CBP, H4K16ac, and HP1α in the left hemisphere. Scale bar, 500 μm. **g,** Higher-magnification view from the thalamus region in (f) highlighting nuclear intensity variation. Scale bar, 50 μm. **h,** Composite of H4K8ac, mH2A1, and SF3a66 within the hippocampus. Scale bar, 100 μm. **i,** Zoomed cortical region showing heterogeneous distribution of H4K8ac and mH2A1. Scale bar, 50 μm. All images were acquired using orthogonal 3D rendering in Imaris.

The imaging system is equipped with a 40× long-working-distance (300 µm) silicone oil immersion objective with a high numerical aperture (N.A.) of 1.25 and a spinning disk-based optical sectioning module coupled to a beam uniformizer. This configuration ensures homogeneous illumination across the entire field of view (Extended Data Fig. 9), a critical factor for unbiased intensity quantification across large imaging areas.

For downstream analysis, we developed a distributed computational pipeline that performs image stitching^16^, cross-round image registration^17^ (Extended Data Fig. 10), RNA spot detection^18^ (Extended Data Fig. 11), and multiscale rendering for efficient visualization. Single-cell segmentation was further enhanced using a customized and improved implementation of Cellpose^19^, a training-free foundation model that accurately delineates nuclear boundaries (Extended Data Fig. 12) and operating directly on large datasets in a distributed, block-wise manner (Extended Data Fig. 13). Following segmentation, RNA spots were assigned to individual cells to generate tissue-wide spatial gene expression profiles, which exhibited strong concordance with the Allen Brain Atlas ISH reference dataset^20^ (Extended Data Fig. 14).

To evaluate reproducibility, we conducted a second round of imaging focusing on 16 protein targets (Fig. 4e–i and Extended Data Fig. 15) alongside the same set of 79 RNA targets. The resulting RNA and protein maps closely matched those from the initial dataset (Extended Data Fig. 5–8, 13, 14), and quantitative analysis across biological replicates revealed high correlations (r = ∼0.9 and P < 10⁻^6^) at both RNA and protein levels (Extended Data Fig. 16), confirming the robustness and reproducibility of the cross-modular cycleHCR framework.

### High-fidelity, tissue-wide imaging reveals spatial epigenetic heterogeneity

The HCR-based signal amplification yielded exceptionally high signal-to-noise ratios, enabling robust detection of diverse proteins spanning a wide dynamic range from abundant cytoskeletal components (e.g., GFAP, Vimentin, MAP2, βIII-tubulin (Tuj1)) to low-abundance transcription regulators and post-translationally modified species (e.g., CBP, H3K27ac, H4K8ac, Pol II Ser5phos; Fig. 4). The resulting tissue-scale datasets preserved fine subcellular detail while maintaining uniform sensitivity across brain regions and imaging depths. This capability enables seamless navigation from brain-wide overviews to subcellular features, revealing distinct regionally enriched cell types (e.g., NeuN, Iba-1, CB1, Calbindin, PV, GABA, NPY, TH, nNOS), vascular networks (CD31), and nuclear structural markers (e.g., MECP2, heterochromatin; H4K16ac, euchromatin; Fig. 3c and Fig. 5a).

**Fig. 5.**
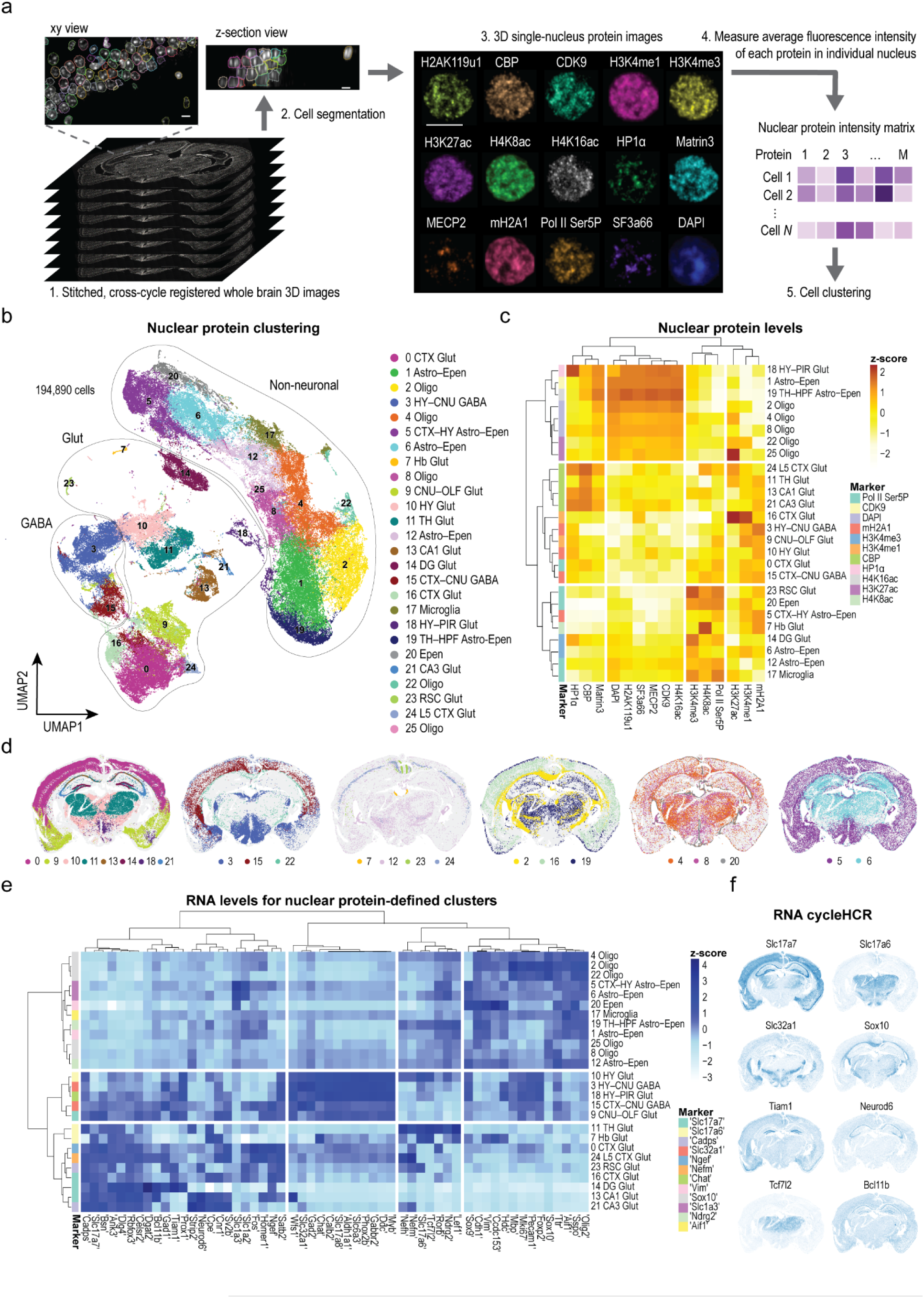
Nuclear protein signatures define spatial cell identities in the brain. **a,** Workflow for constructing a single-nucleus protein fluorescence intensity matrix from a 40-μm mouse whole-brain section imaged by cycleHCR. Nuclear protein images were stitched and registered across imaging cycles, and nuclei were segmented in 3D from DAPI images using Cellpose (194,890 nuclei from Brain Section 2). Average fluorescence intensities of 14 nuclear markers and DAPI were quantified for each nucleus to generate a 15-channel intensity matrix used for UMAP and cell clustering. Scale bars, 10 μm. **b,** UMAP visualization of single-nucleus clusters based on fluorescence intensities of 15 nuclear markers. Clusters correspond to major cell types - non-neuronal (astrocytes, ependymal cells, oligodendrocytes, microglia), glutamatergic neurons, and GABAergic neurons - annotated by matching to RNA expression profiles in (e). **c,** Heatmap of normalized mean fluorescence intensities for nuclear markers across clusters. Marker identity for each cluster corresponds to the nuclear or epigenomic protein with the highest log fold-change relative to other clusters. **d,** Spatial distribution of protein-defined clusters across the mouse brain reveals distinct anatomical organization. **e,** Heatmap of normalized RNA expression levels for clusters in (b), showing strong concordance between protein- and RNA-defined classifications. Cell types were assigned using cell type marker genes exhibiting a high log fold-change relative to other clusters. **f,** Representative RNA cycleHCR spatial maps illustrating gene-expression domains corresponding to the protein-based clusters. Matrix processing and cell clustering were performed using the Squidpy pipeline.

During data exploration, we observed pronounced variability in protein levels of transcription-associated and epigenetic regulators previously considered to be ubiquitously expressed (e.g., CDK9, SF3A66, CBP, H4K8ac, HP1α, Pol II Ser5P, H4K8ac, and mH2A1; Fig. 4e–i). This heterogeneity was evident across both brain regions and cell types. For example, the transcriptional coactivator CBP displayed higher concentrations in the thalamus compared to the hypothalamus (Fig. 4f), and cells in the cerebral cortex exhibited distinct epigenetic states demarcated by the sharp morphological boundary relative to neighboring cerebral nuclei (Extended Data Fig. 17). Similarly, H4K8ac levels were substantially elevated in ependymal cells lining the ventricles compared to surrounding cell populations (Fig. 4g). Notably, even components of the general transcriptional machinery, including Pol II Ser5P (transcription initiation), CDK9 (elongation), and SF3A66 (splicing), showed clear gradient patterns within the hippocampus (Fig. 4e,h; Extended Data Fig. 15). Together, these observations suggest that epigenetic regulation in the brain is highly context-dependent and spatially organized. The resulting diversity in protein composition and chromatin states provides a rich molecular landscape that may serve as the basis for cell-type classification directly from imaging-based spatial proteomic data.

### Nuclear protein profiles accurately classify brain cell types

Tissue-scale, imaging-based cell type classification has traditionally relied on multiplex RNA profiling^20, 21^. To test whether cell types can be distinguished directly from protein-level information, we developed a computational pipeline to extract individual nuclear images from 3D-segmented volumes and quantify nuclear protein enrichment across all detected targets. The resulting single-cell protein intensity matrices were subjected to dimensionality reduction and clustering using Uniform Manifold Approximation and Projection (UMAP)^22^ analysis (Fig. 5a).

As an initial test, we applied this approach to cultured primary hippocampal neurons and astrocytes, achieving 99.7% separation accuracy between the two cell types in unsupervised UMAP clustering (Extended Data Fig. 18). Similarly, extending the same framework to mouse brain tissue revealed distinct clusters (Fig. 5b), each characterized by unique variations in nuclear protein and epigenetic modification levels (Fig. 5c). These clusters were not only separated in UMAP space but also exhibited distinct spatial distributions within the tissue (Fig. 5d), indicating that nuclear protein heterogeneity also encodes spatial information in tissue organization. Consistent with the strong correlations observed between protein measurements in two biological replicates (Extended Data Fig. 16), UMAP projections derived from independent brain datasets showed highly consistent cluster structure and homogeneous mixing (Extended Data Fig. 19), confirming the robustness and reproducibility of the protein-based classification. Similarly, UMAP analysis based on RNA expression profiles from the same two samples yielded highly concordant, spatially distinct clusters across replicates (Extended Data Fig. 20).

To assign cell-type identities to these clusters, we annotated the protein-based classifications by integrating RNA expression profiles from 79 cell type-specific marker genes and anatomical locations (Fig. 5e,f). At the broadest level, the resulting annotations corresponded to non-neuronal, glutamatergic, and GABAergic neuronal populations. Within these groups, finer subclusters reflected regionally distinct glial and neuronal cell types (Fig. 5b–e with annotated cluster labels; Supplementary Table 3). Together, these results demonstrate that nuclear protein and epigenetic modification profiles alone can accurately predict cell identity and spatial position within complex brain tissue, uncovering previously unrecognized heterogeneity in nuclear and epigenetic states even among closely related cell types.

### Explainable ML-guided discovery of cell-type-specific structural features

While UMAP-based clustering relied exclusively on nuclear protein abundance, the barcode-expanded cycleHCR platform additionally captures rich subcellular organization at diffraction-limited resolution (∼260 nm). Such high-dimensional imaging data provide an opportunity to move beyond molecular abundance and directly interrogate structural features associated with cell identity. Deep neural networks are well suited for learning and classifying such high-dimensional image data and have recently been extended to enable cross-domain image translation^23^.

As a conservative proof of concept, we focused our initial analysis on four well-defined, spatially distinct excitatory neuronal populations (TH Glut, CA1, CA3, and layer 5 cortical glutamatergic neurons), selected to span distinct anatomical regions and epigenetic states while enabling controlled evaluation of the framework. Individual nuclei were extracted using Cellpose-based segmentation masks, and convolutional neural network (CNN) classifiers were trained on single-channel nuclear protein images labeled by protein-based cell-type clusters. Across individual protein channels, the classifiers achieved F1 scores ranging from 0.55 to 0.95, compared with 0.25 expected by random assignment (Extended Data Fig. 21a), demonstrating that images of individual nuclear markers contain substantial discriminative information for cell-type classification. As expected, classification performance improved with multi-channel integration: combining all 15 nuclear protein channels achieved an average F1 score of 0.95 and classification accuracy of 0.987. SHAP (SHapley Additive exPlanations)^24^ analysis using single-cell nuclear protein intensities further revealed that individual nuclear proteins contribute differentially to cell-type prediction (Extended Data Fig. 22), highlighting cell-type-specific weighting of distinct protein channels by the classification model.

To uncover subnuclear structural features underlying these distinctions, we applied Quantitative Attributions with Counterfactuals (QuAC)^25^, an explainable machine learning framework that integrates cross-domain image translation with quantitative feature attribution. Using a generative adversarial network (GAN)-based counterfactual transformation^26^, nuclei were computationally transformed from one predicted cell type to another while preserving overall morphology and spatial context. QuAC then quantified the minimal, spatially localized changes required to switch classification outcomes, generating attribution masks that highlight the most influential subcellular regions (Fig. 6a; Extended Data Fig. 23).

**Fig. 6.**
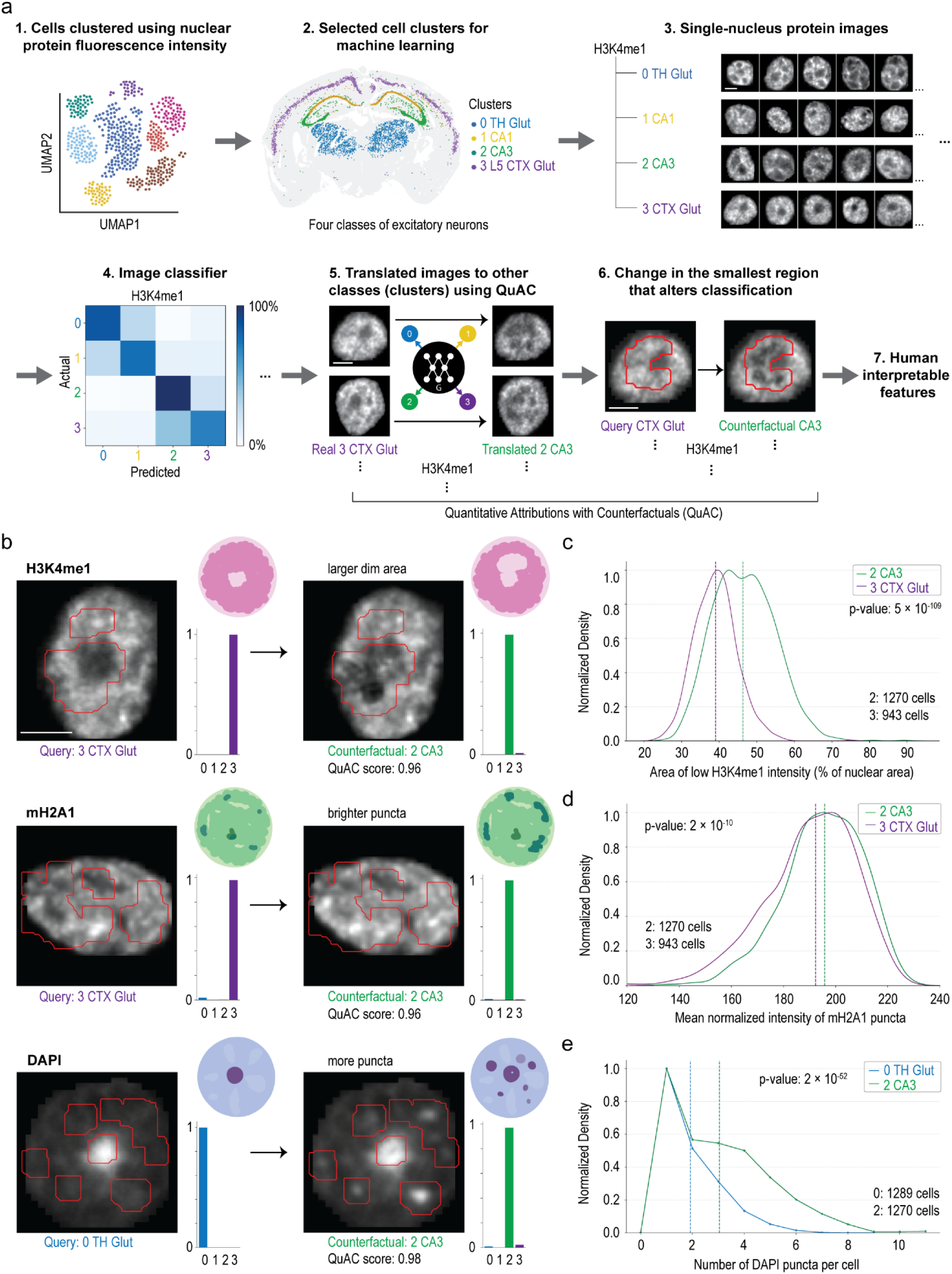
Cell-type-specific nuclear architectures revealed by explainable ML. **a,** Workflow for identifying cell type-specific structural features in protein cycleHCR images using QuAC. Four clusters of excitatory neurons from distinct anatomical regions were selected from the UMAP classification in Fig. 5b; ∼2,000 cells per cluster). Single-nucleus images for each nuclear marker were extracted using Cellpose segmentation masks and divided into training (70%) and validation (30%) datasets. One-channel classifiers were trained for each marker using a modified MONAI workflow with a pretrained DenseNet169 model. QuAC employs a StarGAN-based translation network to generate counterfactual images - synthetic versions of a nucleus transformed into another cell class - and identifies minimal attribution masks whose replacement is sufficient to alter classification. Human inspection of query and counterfactual pairs reveals recurrent structural changes associated with specific cell classes. **b,** Representative query and counterfactual images for three nuclear markers. Red contours denote attribution masks; adjacent schematics illustrate interpreted structural differences. Classification probabilities for each class are shown at right. The QuAC score quantifies attribution efficiency as a function of mask area and change in classification probability between query and counterfactual images. **c,** Normalized density plot of the proportion of dark (heterochromatic) nuclear area across all H3K4me1 training images, identified using Otsu thresholding^31^. **d,** Normalized density plot of the mean normalized intensity of bright puncta (top 10% brightest pixels) in all mH2A1 training images. **e,** Normalized density plot of the number of bright puncta across all DAPI training images, determined using Otsu thresholding. **c–e,** Median values are indicated by dashed vertical lines. P values were calculated using pairwise Mann-Whitney U tests. Scale bars, 5 μm.

These attribution masks revealed biologically interpretable, cell-type-specific structural differences on single-channel nuclear protein images. For example, compared with cortical glutamatergic neurons, CA3 neurons exhibited enlarged central nuclear regions depleted of H3K4me1, a euchromatin-associated mark, alongside expanded peripheral domains enriched in mH2A1, a repressive histone variant. CA3 neurons also displayed more dispersed DAPI puncta relative to thalamic glutamatergic neurons (Fig. 6b). Importantly, these ML-identified features guided the design of deterministic measurements and were independently validated using Otsu thresholding with high statistical confidence (P < 10⁻⁹; Fig. 6c–e), confirming the robustness of the analysis.

Applying the same framework to additional cell populations revealed distinct epigenetic architectures across neuronal and glial classes. For instance, whereas active chromatin marked by H3K4me1 or Pol II was depleted from the nuclear center in cortical GABAergic neurons, astrocyte/ependymal nuclei exhibited centrally enriched active chromatin. This QuAC-identified feature was independently validated at the population level using radial intensity distribution analysis (P < 10⁻^68^; Extended Data Fig. 23). Together, these results demonstrate that the explainable ML framework can extract compact, interpretable, and biologically meaningful subnuclear features directly from imaging data. Although illustrated here using a limited number of representative cell types, the approach is inherently general: extension to additional neuronal and non-neuronal populations requires no modification to the experimental or computational pipeline. By linking subnuclear molecular architecture to cellular identity, this ML-guided spatial omics framework provides both a discovery tool and a principled basis for designing quantitative validation assays.

## Discussion

In this study, we sought to address several practical challenges in spatial omics that have limited the ability to jointly analyze molecular state, subcellular organization, and cell identity at tissue scale. Spatial transcriptomic approaches offer high multiplexing but do not directly capture subcellular protein composition or chromatin state, while existing spatial proteomics methods are often constrained by multiplexing depth, throughput, signal amplification, or integration with RNA measurements. In parallel, although high-resolution imaging can capture rich subcellular information, systematically extracting and interpreting such features across large tissue volumes has remained technically challenging. Here, we developed and demonstrated a cross-modular spatial omics platform that integrates multiplexed protein and RNA imaging with scalable computation modules and explainable machine learning to overcome these limitations (Fig. 1). By expanding high-specificity antibody barcoding, implementing a cost-efficient hierarchical labeling strategy, and incorporating deep-learning-based feature attribution, we mapped transcriptional, epigenetic, and structural heterogeneity across hundreds of thousands of cells in intact mouse brain tissue. Our results show that nuclear protein and chromatin composition alone can accurately predict brain cell identity and spatial position, while explainable ML further reveals interpretable subnuclear signatures that distinguish cell types and anatomical regions. Together, these advances establish a scalable and quantitative spatial framework for acquiring and translating high-dimensional imaging data into integrated molecular profiles and interpretable subcellular architecture within native tissue environments.

Looking ahead, this imaging and computational pipeline can be extended to diseased, aged, or genetically modified tissues to identify epigenetic and subcellular signatures associated with disease states and aging. The resulting image-derived molecular and structural features may provide a quantitative resource for hypothesis generation and for developing biomarkers of disease classification and prognosis. In addition, coupling this platform with genetic or small-molecule perturbation screens would enable systematic, in situ analysis of how specific perturbations influence transcriptional output and subcellular protein organization. Together, these applications offer a general framework for studying gene regulation, cellular architecture, and cell identity within native tissue contexts.

Despite its broad applicability, the current cycleHCR spatial omics platform has several practical limitations. The achievable multiplexing depth is constrained not by barcode design but by the availability and reliability of antibodies that maintain their performance under standardized staining, clearing, and imaging conditions. Extending highly multiplexed protein and RNA cycleHCR imaging to deeper tissue requires investigating protocol compatibility with deep-tissue clearing and immunostaining strategies^27–29^ and optimizing barcode-hybridization uniformity and signal-amplification chemistry for robust performance at depth^30^.

The present explainable ML analysis focuses primarily on single-channel nuclear protein features. This design choice reflects both technical robustness and biological relevance: the nucleus provides a well-defined and reproducible reference compartment that can be segmented reliably at tissue scale, enabling consistent quantitative analysis across large datasets. In contrast, accurate segmentation of cytoplasmic compartments in dense tissues remains challenging and would introduce additional uncertainty into feature extraction. Importantly, this focus is not intrinsic to the cycleHCR platform or the underlying ML framework. As cytoplasmic and whole-cell segmentation foundation models continue to mature, the same analysis pipeline can be readily extended to incorporate cytoplasmic and organelle-level features, enabling more comprehensive characterization of cellular architecture. More broadly, fully leveraging the high-dimensional nature of these datasets will require new computational approaches capable of identifying cross-channel correlations and resolving relative subcellular changes within multidimensional feature spaces. Finally, while the current explainable ML framework outputs attribution masks for expert interpretation, future integration with image-to-language models, enabled by expanded expert annotations, could further automate the translation of structural features into human-readable biological descriptions.

In summary, the integration of cycleHCR-based multiplexed protein and RNA imaging with scalable computational modules and explainable ML establishes a robust foundation for multiscale spatial investigations of biological systems in intact tissues. Continued progress in antibody validation, deep-tissue-compatible sample preparation, and computational analysis will expand the potential of this platform for mechanistic discovery, high-content screening, and translational studies.

## Acknowledgments

We thank Monique Copeland and Benjamin Foster from the Histology Shared Resources at Janelia Research Campus for preparing hippocampal and brain slices. We also thank Deepika Walpita and Phuong Nguyen from the Primary and iPS Cell Culture team at Janelia for preparing hippocampal primary cell cultures. In addition, we acknowledge Melanie Radcliff for her administrative support throughout this project. We appreciate the support from the HHMI Janelia Open Science Software Initiative (OSSI) for maintaining BigStitcher(-Spark), Cellpose, and RS-FISH. This article is subject to HHMI’s Open Access to Publications policy. HHMI lab heads have previously granted a nonexclusive CC BY 4.0 license to the public and a sublicensable license to HHMI in their research articles. Pursuant to those licenses, the author-accepted manuscript of this article can be made freely available under a CC BY 4.0 license immediately upon publication.

## Author contributions

Z.J.L. and Y.L. conceived and designed the study. Z.J.L. conducted fluidics, and microscope assembly and programming, and performed probe design. Y.L. conducted cycleHCR RNA and protein imaging experiments in cultured cells, hippocampal sections, and whole-brain tissue, and performed image processing and single-cell analyses, including image stitching, registration, RNA spot detection, 3D cell segmentation, spot-to-cell assignment, mask-based intensity quantification, UMAP analysis, and gene-to-cluster annotation. D.A. and J.F. developed the QuAC analysis framework. Y.L. and D.A. performed QuAC analyses. T.K., J.K., G.F., and S.P. contributed to development of the image-processing workflow, including distributed image stitching, image registration, RNA spot calling, cell segmentation, and spot-to-cell assignment. Z.J.L. and Y.L. wrote the manuscript with input from all authors.

## Methods

### Cell line

NIH/3T3 cells (ATCC, CRL-1658) were maintained in DMEM (ATCC, 30-2002) supplemented with 10% fetal bovine serum (Gibco, 26140079), 1× GlutaMAX™ Supplement (GIBCO, 35050-061), and 1× Antibiotic-Antimycotic (GIBCO, 15240062) at 37°C in a humidified incubator with 5% CO_2_. For cycleHCR experiments, cells were plated onto cleaned, silanized, and gelatin-coated 40-mm coverslips (Bioptechs, 40-1313-0319; see method below).

### Hippocampal primary cell culture

Dissociated hippocampal primary cells were prepared from postnatal day 3 C57Bl/6 mouse pups (Charles River). The hippocampi were dissected out and digested with papain (Worthington Biochemical). After digestion, the tissues were gently triturated and filtered with the cell strainer. The cell density was counted and ∼8 × 10^5^ cells were plated onto each 40-mm coverslip (Bioptechs, 40-1313-0319) cleaned, silanized, and coated with poly-D-lysine (Sigma Aldrich, P7280; see methods below). The primary cells were maintained in NbActiv4 medium (BrainBits) at 37°C for 7 days before processing for cycleHCR.

### Mouse brain sections

Adult C57BL/6J mice (3–8 months old; Jackson Laboratory, Strain #000664) of both sexes were used in this study. All animal procedures were performed in accordance with protocols approved by the Janelia Research Campus Institutional Animal Care and Use Committee (IACUC). Mice were maintained in a 12-hour light/dark cycle.

#### PFA-fixed whole brain sections

Mice were anesthetized with isoflurane until unresponsive to toe-pinch and transcardially perfused with 15 ml RNase-free PBS (pH 7.4) followed by 50 ml 4% paraformaldehyde (PFA; Electron Microscopy Sciences, 15710) in 0.1 M phosphate buffer (pH 7.4). Brains were removed and post-fixed in a 4% PFA solution in PBS overnight at 4°C, rinsed in PBS, then cryoprotected in 30% (w/v) RNase-free sucrose (Sigma-Aldrich, S7903) in PBS with gentle agitation overnight until equilibrated (tissue sank). Brains were embedded in OCT compound (Fisher Scientific, 23-730-571). 40-μm coronal sections were cut from Bregma −1.5 mm to - 2.5 mm. Cryosections were mounted onto cleaned, silanized, poly-D-lysine-coated coverslips.

#### Fresh-frozen hippocampal section

Mice were anesthetized with isoflurane, confirmed unresponsive to toe-pinch, then decapitated. Hippocampi were rapidly dissected, oriented in OCT within Peel-A-Way embedding molds (Millipore Sigma, E6032), and snap-frozen by immersion in a dry ice/ethanol mixture (∼ 5:1 ratio) for approximately 5 minutes until solid. OCT blocks were equilibrated to −14°C in a cryostat (Leica, CM3050S) for 1 hour before sectioning. Coronal sections (40 μm) were cut around Bregma −1.6 mm and directly mounted on cleaned, silanized, poly-D-lysine-coated coverslips. Brain and hippocampal sections mounted on coverslips were maintained at −80°C before cycleHCR sample preparation.

### Coverslip preparation

Coverslips (40 mm; Bioptechs, 40-1313-0319) were cleaned, silanized, and coated with poly-D-lysine following a published protocol with minor modifications^55^. Cleaned coverslips were silanized by immersion in 0.2% (v/v) triethylamine (Sigma-Aldrich, 471283) and 0.3% (v/v) allyltrichlorosilane (Sigma-Aldrich, 107778) in chloroform (Fisher, C298-500). Silanized coverslips were coated with 100 μg/ml poly-D-lysine (Sigma-Aldrich, P7280) in PBS overnight at room temperature. Poly-D-lysine coating promotes tissue and cell adhesion on the glass surface, while silanization enables covalent attachment of polyacrylamide hydrogels. For NIH/3T3 cell adhesion, silanized coverslips were coated with 0.1% gelatin solution (Millipore, ES-006) for 30 minutes at room temperature.

### cycleHCR probe design

#### L and R barcoding probe sequence expansion for multiplex protein cycleHCR imaging

To expand the L and R barcode space for multiplex protein cycleHCR imaging, we generated candidate 14-bp sequences using the randseq function in MATLAB 2022 and subjected them to a stringent multi-stage filtering pipeline. Sequences containing homopolymer runs of four or more identical nucleotides were excluded to prevent undesirable secondary structures, and candidates predicted to form hairpins or self-dimers were removed. Remaining sequences were restricted to a GC content of 30–60% and a predicted melting temperature (Tm) below 50°C to ensure selective hybridization under assay conditions. Each candidate was then screened against all previously published 30-L and 30-R RNA barcode sequences^11^, their reverse complements, and all newly generated sequences to ensure a maximum cross-hybridization Tm below 32 °C, the operating temperature for all cycleHCR hybridization and imaging steps. This workflow yielded a set of 60 left and 60 right barcodes per color channel. To enable amplification, each left barcode was fused to a color-specific 18-bp split HCR initiator via a 2-bp spacer (AA), and each right barcode was similarly fused to the complementary split initiator. Split-initiator sets B4, B1, and B2 were used for the 488 nm, 561 nm, and 640 nm channels, respectively, following HCR v3.0 design principles^12^. All final L and R barcoding probes were synthesized by Integrated DNA Technologies (IDT).

#### Protein cycleHCR oYo docking and gel anchoring probe design

Fifty 20-bp oYo docking sequences were generated using MATLAB randseq function and screened to ensure a maximum predicted cross-hybridization melting temperature below 40 °C. Oligonucleotides bearing 5′ oYo photocrosslinker modifications were synthesized by AlphaThera, Inc. Before assigning each docking sequence to a specific L+R barcode combination, secondary structure propensity was evaluated with the MATLAB oligoprop function, and the docking sequence with minimal predicted hairpin formation was selected. Gel-anchoring probes were constructed by concatenating the reverse complement of the 20-bp docking strand, a 5-bp randomized linker, the reverse complement of the right barcode, a 2-bp spacer (AA), and the reverse complement of the left barcode. A 5′ acrydite modification (/5Acryd/) was included during IDT synthesis to enable covalent incorporation into the acrylamide hydrogel matrix. All docking sequences, anchoring sequences, and barcode assignments used in this study are listed in Supplementary Table 4.

#### RNA cycleHCR primary probe library assembly

Primary probes for RNA cycleHCR imaging were designed following our previously described framework^11^. For each RNA target, 5–25 primary probe pairs were generated. Each left probe was assembled by concatenating a forward PCR handle containing a T7 promoter (TAATACGACTCACTATAGCGTCATC), the first 45 nt of the 92-nt target region, a TT spacer, a 14-bp left barcode, and a universal reverse oligo (CGACACCGAACGTGCGACAA). Each right probe contained the same T7-promoter PCR handle followed by the 14-bp right barcode, a TT spacer, the final 45 nt of the target region, and the same universal reverse oligo. All primary probes were 92 nt in length and were subsequently amplified and transcribed according to standard cycleHCR protocols. Probe sequences for all 79 RNA targets used for the mouse brain are provided in Supplementary Table 2.

### Antibody validation

To demonstrate multiplex protein cycleHCR, antibodies were first tested using secondary antibody staining on mouse brain sections prepared with the SHIELD protocol^27^. Immunostaining patterns were compared to images provided on antibody vendor websites or in the literature (Supplementary Table 1). Antibodies validated by secondary antibody staining were then tested using cycleHCR (see methods below) on SHIELD-treated mouse brain sections. Antibodies that performed successfully with cycleHCR were included in the final antibody pool for image acquisition. Antibodies intended for cultured cells and hippocampal sections were validated on corresponding PFA-fixed samples.

### Antibody-DNA probe complex construction for protein cycleHCR

Each antibody was independently conjugated to its assigned oYo-Link DNA docking strand and subsequently hybridized in vitro with its corresponding gel-anchoring probe. DNA docking strands are 20-nt oYo-Link DNA oligos (5’ linked; AlphaThera) conjugated to antibodies using a light-activated photocrosslinking reaction following the manufacturer’s instructions. Briefly, 1 μl 33 μM oYo-Link oligo solution was gently mixed with 1 μg antibody before reaction in the photo-crosslinking device. Gel-anchoring probes are 55-nt DNA oligos bearing a 5′ acrydite modification (IDT). For in vitro hybridization of DNA docking strand with gel-anchoring probe, 1 μl oYo-oligo-conjugated antibody was combined with 1 μl of 50 μM gel-anchoring probe in PBS and 0.7 μl of 40% dextran sulfate (Sigma-Aldrich, D4911) in PBS. The mixture was incubated at room temperature for 1 hour with gentle shaking.

Conjugated and in vitro hybridized antibody-DNA probe complexes (1 μl each) were combined in blocking solution with final concentrations of 0.25% (v/v) Triton X-100 (Sigma-Aldrich, X100), 0.5 mg/ml sheared salmon sperm DNA (Invitrogen, AM9680), 10 mg/ml nuclease- and proteinase-free BSA (Sigma-Aldrich, 126609), and 1× PBS. To reduce non-specific binding, rabbit, mouse, and goat IgG (Sigma-Aldrich, I5006, I5381, I5256) were added at a 10-fold excess relative to their corresponding primary antibodies. For joint protein and RNA cycleHCR, SUPERase·In RNase Inhibitor (Invitrogen, AM2694) was included at a final concentration of 1 U/μl. The final volume of the antibody-DNA probe complex solution was 100–120 μl.

### RNA primary probe construction

RNA primary probes were constructed using a previously described enzymatic amplification workflow^1^. Briefly, the RNA primary probe oligo library (Twist Bioscience) was amplified by PCR using KAPA HiFi HotStart (Roche, KK2502) with primers containing T7 promoter sequences (IDT). The PCR product was purified using the DNA Clean and Concentrator Kit (Zymo Research, D4003) and re-amplified before being converted into RNA via in vitro transcription using the HiScribe T7 High Yield RNA Synthesis Kit [New England Biolabs (NEB), E2040S] for 7 hours to overnight. The RNA product was purified using the Monarch RNA Cleanup Kit (NEB, T2040) and converted back to DNA via reverse transcription using Maxima H Minus Reverse Transcriptase (Thermo Scientific, EP0753). The RNA template was removed by overnight digestion with Thermolabile USER II Enzyme (NEB, M5508) followed by alkaline hydrolysis with NaOH (final concentration 1 M) at 65°C for 15 minutes. Hydrolysis was neutralized with acetic acid (final concentration 1 M). The resulting ssDNA primary probes were purified using the ssDNA/RNA Clean & Concentrator (Zymo Research, D7011).

### Cells and fresh frozen tissue immunostaining for cycleHCR

Cells and 40-μm fresh frozen hippocampal sections were fixed in 4% PFA for 10 minutes, followed by 3 washes in PBS for 5 minutes each. Cells were permeabilized in 0.5% Triton X-100 in PBS for 15 minutes at room temperature, while 40-μm hippocampal sections were permeabilized in 0.5% Triton X-100 in PBS for 30 minutes at room temperature. After permeabilization, samples were washed 3 times in PBS for 5 minutes each and incubated in blocking solution for 1 hour at room temperature. After blocking, samples were incubated in the antibody-DNA probe complex solution in an in-house-built hybridization chamber (see method below) overnight at 4°C. Immunostaining was completed by washing samples 3 times in PBS for 5 minutes each.

### Mouse whole brain section immunostaining for cycleHCR

Whole brain sections were prepared using the SHIELD protocol^27^ prior to joint protein and RNA cycleHCR to ensure uniform biomolecule preservation and effective tissue clearing. PFA-fixed mouse whole brain sections were post-fixed in a mixture of SHIELD-ON buffer and SHIELD-Epoxy solution (LifeCanvas) at a ratio of 7:1. Sections were incubated in this epoxy solution at 4°C for 3 hours and subsequently at room temperature for 2 hours with gentle shaking.

Following fixation, sections were cleared in delipidation buffer [300 mM SDS (Fisher, BP166-500), 10 mM sodium borate (prepared by dissolving boric acid; Thermo Scientific, 42348100), 100 mM sodium sulfite (Sigma-Aldrich, S0505), pH 9.0] at 37°C for 4 hours to overnight with gentle shaking. After clearing, sections were rinsed 3 times in PBS, then washed in 1% Triton X-100 in PBS at room temperature with gentle shaking for 1 hour to thoroughly remove residual SDS. Finally, sections were incubated in blocking solution at room temperature for 1–2 hours prior to downstream immunostaining steps.

For imaging 46 proteins in Brain Section 1, immunostaining was performed in two sequential steps. First, the section was incubated in a solution containing 18 antibody-DNA probe complexes targeting nuclear proteins (Supplementary Table 4, #1–18) at 4°C overnight. The section was then washed 3 times in PBS for 5 minutes each, fixed in 4% PFA for 5 minutes at room temperature, and washed again in PBS (3 × 5 min). Next, the section was incubated in a solution containing the remaining 28 antibody-DNA probe complexes (Supplementary Table 4, #19–46) at 4°C overnight. For imaging 16 proteins in Brain Section 2, the section was incubated at 4°C overnight in the corresponding antibody-DNA probe complex solution (Supplementary Table 4, #1–9, 12–16, 18, 23). Immunostaining was terminated by washing the sections 3 times in PBS for 5 minutes each.

### Hybridization chamber construction

Custom hybridization chambers were constructed to facilitate controlled-volume (∼100 μl) incubations for procedures such as immunostaining, RNA probe hybridization, and HCR amplification. To assemble the chambers, a 19-mm-diameter ring gasket was cut from a 500-μm-thick silicone sheet (Bioptechs, 1907-1422-500) using a hollow punch (13-in-1 heavy-duty set). The gasket was then affixed to a pre-cleaned glass slide (Corning, 2947-75X50) using 100% silicone sealant (Gorilla). During incubations, sample-bearing coverslips were inverted onto the chambers with the tissue immersed in the incubation solution within the gasket well.

### Gel embedding

After immunostaining, samples were incubated in 0.1 mg/ml Acryloyl-X SE (AcX; Invitrogen, A20770) solution in PBS for 1 hour at room temperature. AcX solution was prepared and preserved as previously described^56^. Samples were washed twice in PBS for 15 minutes each. To ensure homogeneous penetration of acrylamide solution into tissues before gelation, hippocampal and brain sections were incubated in acrylamide solution [4% acrylamide/bis-acrylamide (Bio-Rad, 1610154), 60 mM Tris-HCl pH 8 (Corning, 46-031-CM), 0.3 M NaCl (Corning, 46032)] on ice for 10 minutes with gentle shaking. Gelation solution was prepared by adding ammonium persulfate (Sigma-Aldrich, A3678) and TEMED (Sigma-Aldrich, T7024) to the acrylamide solution to final concentrations of 0.03% and 0.15%, respectively. For gel embedding, 100 μl of gelation solution was pipetted onto a Gel Slick (Lonza, 50640)-coated glass slide (Corning, 2947-75X50). The sample-bearing coverslip was slowly lowered onto the solution to immerse the tissue or cells without trapping air bubbles. After 1.5 hours at room temperature, the coverslip was carefully lifted from the slide using a syringe needle without damaging the gel. Excess gel on the coverslip was trimmed with a razor blade to fit within a 19-mm-diameter hybridization chamber.

### Proteinase K digestion

Cells were digested using a previously described protocol^55^. Gel-embedded brain sections were digested with slight modifications to a previous protocol^56^. Sections were incubated overnight at 37°C with proteinase K (80 U/ml; 1:100 dilution; NEB, P8107S) in 2× SSC (Invitrogen, 15557044) containing 2% SDS and 0.5% Triton X-100. Digestion was terminated by washing the sections twice in 5× SSC-T [5× SSC, 0.1% Tween 20 (Bio-Rad, 1610781)] for 15 minutes each.

### RNA primary probe hybridization for cycleHCR

Proteinase K-digested brain sections were hybridized with pre-amplified RNA primary probes targeting 79 genes. The primary probe library (∼25 μg ssDNA) was concentrated to approximately 10 μl using a Vacufuge Plus (Eppendorf) in vacuum-aqueous mode at room temperature, then diluted in 90 μl of hybridization buffer [50% (v/v) formamide (deionized; Ambion, AM9342), 10% (w/v) dextran sulfate, 2× SSC] for a final volume of ∼100 μl. Each brain section was incubated with this probe solution in a hybridization chamber at 37°C for around 48 hours using a RapidFISH Slide Hybridization Incubator (Boekel Scientific, 13-245-230). After RNA primary probe hybridization, sections were washed in pre-warmed 80% formamide solution (80% formamide, 4× SSC, 0.1% Triton X-100) at 32°C for 20 minutes. Sections were rinsed twice and washed for 10 minutes in 5× SSC-T. Samples could be stored in 5× SSC-T at 4°C for weeks before acquiring cycleHCR images using the automated fluidics-microscope system.

### Barcoding probe hybridization and HCR amplification

CycleHCR signals for specific protein and RNA targets can be examined manually before proceeding to automated dataset acquisition. Three protein or RNA targets, one in each fluorescence channel (488 nm, 561 nm, 640 nm), can be examined in each cycle. At the beginning of each cycle, samples were incubated for 1 hour at room temperature in 100 μl of barcoding probe solution containing 200 nM each of Left and Right probes in 10% EC buffer [10% (v/v) ethylene carbonate (Sigma-Aldrich, E26258), 10% dextran sulfate, 2× SSC]. Samples were washed in 10% formamide solution (10% formamide, 0.1% Triton X-100, 2× SSC) for 10 minutes at room temperature, then rinsed twice and washed for 5 minutes in 5× SSC-T. For HCR amplification, hairpin amplifiers h1 and h2 (2 μl each; B4 for 488 nm; B3 for 647 nm; B2 for 560 nm; Molecular Instruments, Inc.) were heated at 95°C for 90 seconds in separate tubes, then cooled to room temperature in the dark for 30 minutes. The snap-cooled h1 and h2 hairpins were mixed in 100 μl of amplification buffer (10% dextran sulfate, 0.1% Tween 20, 5× SSC). Samples were incubated in the hairpin amplification solution at 32°C for 1.5 hours, then washed twice in 5× SSC-T for 10 minutes each.

### Imaging and stripping

Following HCR amplification, samples were assembled in an open FCS2 imaging chamber (Bioptechs) and immersed in 5× SSC-T for imaging. The imaging system and conditions are described below. After imaging, barcoding probes and HCR amplicons were stripped by incubating samples in 80% formamide solution at 32°C for 20 minutes. Samples were rinsed twice and washed for 10 minutes in 5× SSC-T. At this stage, samples were ready for the next cycle of barcoding and HCR amplification targeting additional genes or proteins.

### cycleHCR fluidics and imaging system

#### Automated fluidics system

The fluidics system was engineered to deliver precise flow control and efficient reagent mixing, both critical for cycleHCR imaging. It includes an OB1 flow controller (Elveflow) to regulate system flow rates, two MUX distribution valves (Elveflow) to route reagents to specific channels, and a BFS Coriolis flow sensor (Elveflow) for real-time flow monitoring and feedback control. The configuration accommodates 10 probe tubes for L + R readout probes, 4 solution reservoirs for buffers and wash solutions, and 2 tubes containing the h1 and h2 hairpin solutions. Real-time mixing of h1 and h2 hairpins was achieved by alternating flow between the two tubes using a 2-to-1 MUX valve, preventing premature background amplification. The optimized flow rate was 150 μl/min to balance throughput and sample integrity. All buffers were filtered through 0.22-μm vacuum filters (Corning, 431098) prior to use. Following primary probe hybridization, samples mounted on 40-mm coverslips were assembled in a closed-top FCS2 imaging chamber (Bioptechs) and positioned on a custom stage adapter on the microscope.

#### Fluidics workflow

1. Stripping Buffer (80% formamide, 0.1% Triton X-100, 4× SSC) Incubation (20 min): Removes bound probes and prepares the sample for the next cycle.
2. Wash (20 min): Removes residual reagents using 5× SSC with 0.1% Tween 20 and 5 μg/ml DAPI.
3. Barcoding Probe Incubation (30 min): Introduces mixed Left and Right barcoding probes for target detection in the new cycle. Barcoding probe solution for each cycle was supplied in 1.8 ml of hybridization solution (10% ethylene carbonate, 5× SSC, 10% dextran sulfate) with 200 nM of each probe.
4. Wash (20 min) as in Step 2: Removes excess unbound barcoding probes.
5. Hairpin Mix Injection (5 min): Introduces h1 and h2 hairpins for HCR signal amplification. Stock solutions of h1 and h2 (3 μM; Molecular Instruments) were diluted 1:50 in amplification buffer (5× SSC, 0.1% Tween 20, 10% dextran sulfate). Upon injection, h1 and h2 were mixed at a 1:1 ratio to a final working concentration of 1:100 (30 nM).
6. HCR Amplification (90 min): Incubates samples to allow fluorescent signal buildup.
7. Wash (10 min) as in Step 2 and 4: Removes excess hairpins.
8. Imaging Buffer [50 mM Tris-HCl pH 8, 2 mM Trolox (Sigma-Aldrich, 238813), 1 mg/ml glucose oxidase (Sigma-Aldrich, G2133), 1:50 catalase (Sigma-Aldrich, 02071), 0.8% (w/v) D-glucose, 5× SSC] Injection (20 min): Introduces an oxygen scavenging solution to prevent photo-crosslinking and enhance signal.
9. TTL Signal Activation: Triggers imaging on the microscope after a 10-minute incubation.

#### Microscopy

High-resolution imaging was performed for cycleHCR using a Nikon CSU-W1 spinning disk confocal microscope equipped with these advanced components:

1. Objectives: 40× CFI PLAN Apochromat Lambda S silicone oil immersion objective (N.A. 1.25, working distance 0.3 mm); 40× CFI Apo LWD Lambda S water immersion objective (N.A. 1.15, working distance 0.6 mm; used for hippocampal section and primary cell imaging)
2. Illumination uniformizer: Ensures even illumination across the field of view
3. Six laser lines: 405 nm, 488 nm, 514 nm, 561 nm, 594 nm, and 640 nm for versatile multichannel excitation
4. Camera: Hamamatsu BT Fusion sCMOS camera for high-sensitivity imaging with ultra-low readout noise

#### Imaging conditions and parameters

Images were acquired with 200-ms exposure time in ultra-quiet camera readout mode at 10% laser power. The Nikon Perfect Focusing System (PFS) maintained constant z-position between imaging rounds, ensuring long-term stability throughout multi-cycle acquisition. TTL signals coordinated automated transitions between imaging and fluidic cycles. Temperature was maintained at 32°C to support probe hybridization, HCR amplification, and stripping, using a Tokai Hit lens heater controlled by a TPi controller (TPiE-LH) for the objective, and a Bioptechs FCS controller for the imaging chamber.

#### Volumetric image acquisition

Three-dimensional image stacks were acquired with xy tiling and z-stepping to capture subcellular-resolution volumetric data. Each xy field measured 394.0 μm × 394.0 μm (2304 × 2304 pixels) using the 40× silicone oil immersion objective. Specific parameters varied by sample type:

NIH/3T3 cells: 8 xy tiles sampled across the culture area for representative coverage with 1-μm z-steps; 60 imaging cycles for 3 proteins, one protein per fluorescence channel

Hippocampal tissue sections: 78 xy fields with 20% tile overlap and 1-μm z-steps (91 z-planes acquired); 7 imaging cycles for 7 proteins

Primary hippocampal cells: 40 xy fields with 1-μm z-steps; 7 imaging cycles for 15 proteins

Whole brain sections: Large-format imaging without xy tile overlap and 3-μm z-steps to limit dataset size

Brain Section 1: 379 xy fields, 23 z-planes; 47 imaging cycles for 79 RNA targets and 46 proteins; 14.3 TB raw image dataset

Brain Section 2: 450 xy fields, 23 z-planes; 35 imaging cycles for 79 RNA targets and 16 proteins; 16.1 TB raw image dataset

### Image stitching

3D image tiles of the hippocampal slice were stitched using BigStitcher^16^ and the BigStitcher-Spark framework for distributed processing. After converting the data to multi-resolution N5 format, each imaging round was aligned independently using the 20% xy tile overlap as the initial positional estimate. Calculation of the final 3D affine transformation for each tile involved several steps: pairwise shift estimation via phase correlation, previewing and filtering of pairwise shifts, and global optimization of all shifts. Pairwise shifts were computed on data downsampled 8×8×4 (xyz) with intensities averaged across all channels. The overlapping area was assigned pixel values (with interpolation) from the tile that was imaged first among all tiles overlapping that pixel (one-tile-wins fusion strategy). Image tiles of whole-brain sections, acquired without xy overlap, were converted to N5 format using the same pipeline and positioned using the xyz stage coordinates recorded in the image metadata for each field.

### Image registration

Images from all cycles were registered to a selected reference cycle using the bigstream^17^ (version 1.2.9) Python package to correct for sample or field-of-view shifts across imaging cycles. The reference cycle was chosen from the middle cycles and manually inspected to ensure minimal sample tilt. All other cycles were transformed so that the DAPI channels across all cycles aligned in 3D space.

Since cell culture images were multiple single-tile images, the global affine transformation was sufficient. The affine matrix was estimated using the alignment_pipeline function with a downsampling factor of 2 and a subsampling factor of 2 for 800 iterations. This matrix was then applied to the full image using apply_transform, performing uniform translation, scaling, shear, and rotation.

For hippocampal section images, registration involved sequential global affine and local deformable transformations. The affine matrix was calculated through two sequential alignment steps for 500 iterations: the first step with a subsampling factor of 2 and a downsampling factor of 2; the second step without downsampling. The deformable field was generated using the distributed_piecewise_alignment_pipeline function, initialized with the affine matrix. Deformation was performed with a subsampling factor of 2, a downsampling factor of 2, and a spacing of 128 between control points for 10 iterations. Both the affine and deformable transformations were then executed sequentially using distributed_apply_transform, first applying the global affine correction to the entire image, followed by pixel-wise local deformation. Stitched whole-brain section images were registered using affine transformation only, with the same settings as for the hippocampal section alignment. The larger z-step size (3 μm) used for these samples prevented reliable detection and correction of local tissue deformations.

### Cell segmentation

Nuclei segmentation on hippocampal primary cell cultures was performed using Cellpose^19^ with two separately trained models for xy and yz slices. Images were resampled in the z-direction using ImageJ to achieve isotropic voxels. Both models were trained using the human-in-the-loop feature^57^ with manual annotation and iterative optimization. We implemented a custom 3D segmentation workflow^11^ where flow fields were computed in xy (using the xy model) and in yz and xz (using the yz model) with customized diameters matching the nucleus size. The three orthogonal flows were averaged to generate a consensus flow field, which was used to compute final nuclear masks via Cellpose dynamics.

Nuclei in whole brain sections were segmented in 3D using Cellpose for nuclear protein image analysis. Because of the large dataset size, downsampled 4×4×1 (xyz), stitched, and registered DAPI images were used for Cellpose segmentation. Images were also resampled in the z-direction for isotropic voxels before segmentation. A distributed processing module of Cellpose with GPU acceleration was used to process small image blocks separately, ensuring image blocks could fit in system memory (https://cellpose.readthedocs.io/en/latest/distributed.html). Nuclei segmentation was performed using the 3D mode and ‘cyto3’ model^58^ with a customized diameter. Centers of mass of the cells were computed on the nuclear masks using the center_of_mass function in the scipy.ndimage package.

Segmentation accuracy was evaluated by overlaying raw DAPI images with their corresponding segmentation masks in ImageJ or ORS Dragonfly.

### Cell type classification using nuclear protein intensity in primary cell culture

Neurons and astrocytes in the hippocampal primary cell culture were identified using the cell type markers NeuN and GFAP. A cell was classified as a neuron if the average NeuN fluorescence intensity within its 3D nuclear mask exceeded a threshold of 330. For cells not classified as neurons, the nuclear mask at the middle z-plane was radially expanded by 5 pixels, and the expanded nuclear mask was examined for GFAP expression. Cells with more than 3000 pixels exceeding an intensity value of 3000 in the expanded nuclear mask were classified as astrocytes. The threshold values were empirically optimized using the image batch.

For identified neurons and astrocytes, single-nucleus 3D images of 14 nuclear proteins were extracted from the masked regions in the registered images. The average fluorescence intensity of each protein was calculated per nucleus. Before UMAP analysis, nuclear protein intensities were normalized by linearly scaling the 1st percentile to 0.0 and the 99th percentile to 1.0 without capping. PCA was performed before UMAP^22^ construction and Leiden^59^ clustering using the scanpy^60^ package, using customized parameters to obtain two clusters. The distributions of identified neurons and astrocytes across the two clusters were used to generate a confusion matrix.

### RNA spot detection and spot-to-cell assignment

RNA spots of 79 genes imaged by joint protein and RNA cycleHCR in whole brain sections were detected using RS-FISH^18^. RS-FISH parameters were selected using the software’s interactive tools (Extended Data Fig. 11) and optimized for accurate spot detection with minimal false positives. The same spot detection parameters were applied to both whole-brain sections.

RNA spots were assigned to cells using segmented nuclear masks, expanded to include RNA spots outside nuclei. Nuclear masks in downsampled images [8×8×1 (xyz) from raw images] were expanded through three iterations. In each iteration, voxels in the mask image with 0 values were replaced with the maximum value of their 6 neighbors. Single-cell RNA matrices were generated using the RNA spot-to-cell assignment results.

### Single-cell nuclear protein fluorescence intensity measurement in whole brain sections

For each selected cell, 3D nuclear marker images were extracted from the stitched and registered dataset using the corresponding nuclear mask as the boundary. After converting the images back to the original z scale, nuclear masks [downsampled 8×8×1 (xyz)] smaller than 1500 voxels or with z-depth < 4 voxels were excluded from downstream analyses. Local fine-tuning registration was then performed on each extracted nuclear volume to correct residual misalignments at the single-cell level. Average intensities of nuclear markers were calculated inside each 3D nuclear mask. Multithreaded processing was implemented to extract images from large datasets efficiently.

### UMAP-based cell clustering using nuclear protein fluorescence intensity

Single-nucleus average intensities of 15 nuclear markers were used for UMAP-based clustering of cells in the whole brain sections, including H3K4me1, mH2A1, Pol II Ser5P, CBP, MECP2, H4K8ac, H4K16ac, Matrin3, SF3a66, H3K4me3, HP1α, CDK9, H2AK119u1, H3K27ac, and DAPI. Nuclear protein intensity matrices of single cells were used for UMAP construction and cell clustering. Matrix preprocessing was performed using the scanpy package. Average nuclear protein intensities were normalized per cell using the pp.normalize_total function, and the logarithm of the normalized intensities was calculated using the pp.log1p function in the scanpy package. PCA was performed using the default settings. Harmony^34^ was used to integrate data from the two brain sections in the same UMAP space before UMAP construction and Leiden clustering using customized parameters. Spatial localization of cells colored by cluster was plotted using the squidpy^61^ package.

### RNA and protein expression profiles for nuclear protein-defined cell clusters

RNA expression level was calculated using spot counts/cell. Cells with fewer than 10 RNA spots in total were removed from analysis. Nuclear protein expression level was calculated using average fluorescence intensity in each 3D nuclear mask. RNA and nuclear protein expression of each cluster were compared against expressions in other clusters using the sc.tl.rank_genes_groups function in the scanpy package, after normalization and calculating logarithms as described above. The nuclear protein and the cell type marker gene with the highest log fold change for each cluster against other clusters were assigned as the nuclear protein marker and gene marker of that cluster. Z-scores of each gene or nuclear protein intensity across clusters were calculated for plotting heatmaps using the pheatmap function in R. Genes with high log fold change for each cluster were selected for RNA heatmap plotting. Spatial RNA expression plots of each gene in the whole brain were generated using the normalized logarithmic values with the pl.spatial_scatter function in the squidpy package.

### Cell clustering based on RNA cycleHCR data

UMAP-based cell clustering was performed on filtered and preprocessed single-cell RNA matrices following the same pipeline used for nuclear protein intensity clustering with adjusted parameters.

### Single-nucleus protein image extraction for explainable machine learning

Cell clusters generated based on nuclear protein intensities were used to define cell classes for explainable machine learning. From each cell cluster, 2000 cells were randomly selected to construct the image dataset. Fragmented cells located at the edges of image fields were removed by manual filtering. Selected cells of each class were further randomly assigned to training (70%) and validation (30%) datasets. Nuclear marker images from the middle z-plane of each 3D nuclear mask were extracted for downstream analysis. Single-nucleus images were converted to 8-bit by linearly rescaling intensity values. All images were resized to a uniform 90 × 90 pixels and saved as single-channel TIFF files.

### Image classifier

Image classifiers were trained prior to analysis by the Quantitative Attributions with Counterfactuals (QuAC)^25^ framework for explaining model decisions. A separate classifier was trained for each nuclear marker using a modified multi-spectral microscopy classification workflow from Medical Open Network for AI (MONAI; https://github.com/Project-MONAI).

#### Image loading and processing

Single-channel nuclear images were loaded from pre-organized directories containing training/validation splits with subfolders corresponding to class labels. Image intensities were normalized by linearly scaling the 1st percentile to 0.0 and the 99th percentile to 1.0, with values clipped to the range [0, 1]. To enhance training robustness, random augmentations were applied to the training images: random 90°, 180°, or 270° rotations with a probability of 0.75, random horizontal or vertical flips and random zooming between 0.9× and 1.1× with a probability of 0.5.

#### Model training

A DenseNet169^62^ architecture was used, and initialized to weights pretrained on ImageNet^63^. Models were trained for 50 epochs via back-propagation in PyTorch^64^ using a learning rate of 5×10^-5^ and a cross-entropy loss. Each epoch corresponds to one full iteration over all of the training dataset, with optimization happening in batches of 8 samples at a time. At each batch, model weights were modified to minimize the cross-entropy loss.

#### Classification evaluation

Evaluation metrics were computed on validation data, using the scikit.learn^65^ classification report to monitor precision, recall, accuracy, and F1 score.

### Classification performance with varying numbers of nuclear proteins

To compare image classification performance using different numbers of protein channels, 15-channel nuclear images of the same selected cells for the 4-neuron-cluster dataset (Fig. 6a) were generated. These 15-channel images were used for classification model training and evaluation with the same pipeline described above.

Since training image classification models is computationally intensive, a more comprehensive evaluation of cell type classification performance using varying numbers of nuclear proteins was conducted with nuclear protein intensities using XGBoost^35^ classification. Cell classes were defined by clustering all 194,724 cells from Brain Section 2 into 5 major cell types based on nuclear protein intensities, using the same cell clustering workflow with a lower Leiden resolution. The 5 clusters were annotated based on spatial distribution and gene expression profiles as described above. Nuclear protein intensities of these 5 cell classes were used for classification. Classification models were trained and evaluated using all possible combinations of 3 to 15 nuclear proteins. Cells were randomly split into 70% training and 30% validation datasets. Classification performance metrics, including F1 score, accuracy, area under the ROC curve (AUC), were recorded. Macro-average metrics were calculated for each number of nuclear proteins to examine performance trends as protein number varied.

### Importance of individual proteins for cell type classification

The XGBoost classification model trained with intensities of all 15 nuclear proteins was used for SHAP (SHapley Additive exPlanations)^24^ analysis to assess the contribution of individual proteins to predictions. SHAP value distribution plots were generated for each cell class, showing which proteins had the greatest impact on classification decisions. Overall importance of the proteins was also quantified by calculating the mean absolute SHAP values for each cell cluster.

### QuAC

The explainable machine learning framework QuAC^25^ was obtained from the GitHub repository https://github.com/funkelab/quac. QuAC first trains a conversion network that translates images from one class to another using a StarGAN^26^ model. Training was performed for 40,000 batches of size 16 using the same training and validation datasets for image classification and the trained DenseNet classification model. Checkpoints with the highest conversion rates were visually inspected for realistic generated images before being selected for further analysis. Checkpoints chosen for downstream steps were typically from iterations 35000 – 40000 and exhibited > 0.95 conversion rates for all pairwise class conversions.

In the image-generation step, validation images were converted into other classes. The resulting images were analyzed using discriminative attribution methods to identify the smallest necessary regions of change for converting into another class. Attribution masks were then scored using the QuAC metric, and counterfactual images were created that differ from the original validation image only in the masked region. At least 200 query and counterfactual image pairs with the highest QuAC scores for each nuclear protein were exported for manual inspection. Class-specific nuclear structural features were summarized from features consistently observed across attribution masks.

### Population-level quantification of cell-type-specific nuclear features

Nuclear features identified from QuAC generated images were validated through quantitative measurements on the full training image dataset. Prior to all measurements, background pixels with zero intensity were excluded by applying a non-zero-pixel mask.

1. Dim H3K4me1 regions: Otsu thresholding^31^ was applied to each nucleus. The percentage of pixels below the threshold was computed.
2. Bright puncta intensity in mH2A1 images: For each nucleus, the average intensity of the top 10% brightest pixels was calculated.
3. Bright puncta in DAPI images: Bright regions were identified using Otsu thresholding. Small regions were removed using morphology.remove_small_objects in the scikit-image^66^ package with default parameters. The number of bright puncta was counted as connected bright components using the measure.label function.

Normalized distributions of these measurements were plotted. Pairwise statistical significance between classes was assessed using the Mann-Whitney U test^67^.

For analysis of radial intensity distributions of euchromatin markers H3K4me1 and Pol II Ser5P across two cell-type classes, the centroid of each nuclear image was determined. Distances of all pixels to the centroid were computed and normalized to 0–1 (0 = center; 1 = maximal radius). For each nucleus, radial profiles were generated by averaging intensity values across 100 equally spaced radial bins. Population-level radial profiles (mean ± SD) were obtained by averaging curves across all nuclei from each class. P-values were calculated using the Mann-Whitney U test to compare classes at radial distances of 0 (nuclear center) and 0.7 (near nuclear periphery).

## Code availability

Data analysis pipelines are available on GitHub (https://github.com/liulabspatial/Omics-cycleHCR-QuAC).

## Extended Data Figure and Figure Legends 1 - 23

**Extended Data Fig. 1.**
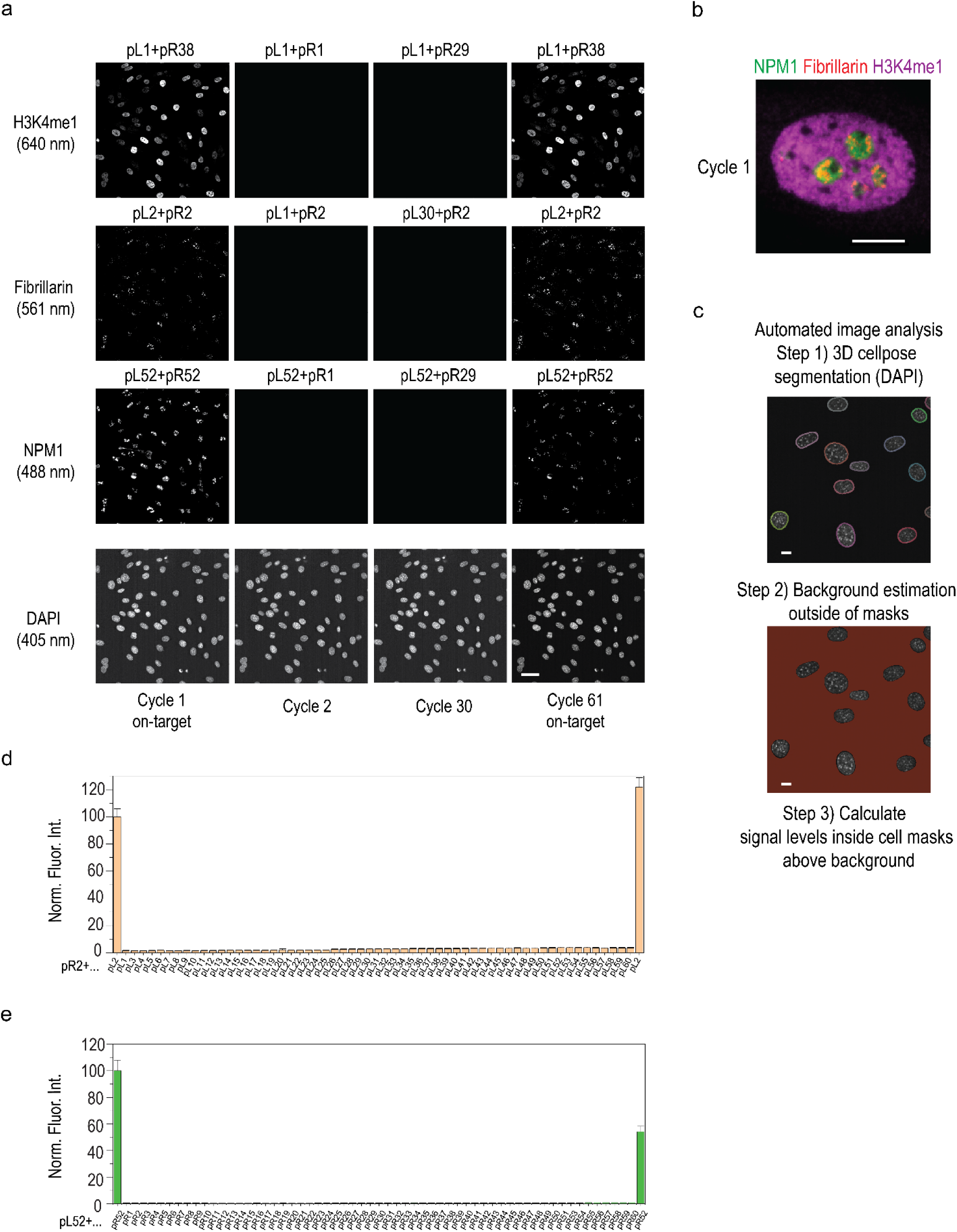
Evaluation of cross-reactivity of L and R barcodes. **a**, Representative images are shown for imaging cycles with on-target barcodes (first and last cycles) and off-target L/R barcodes for the three cycleHCR fluorescence channels. Similar to H3K4me1 imaged for the 640-nm channel as shown in Fig. 1c, fibrillarin was imaged with the same R barcode while cycling through all L barcodes for the 561-nm channel; NPM1 was imaged with the same L barcode while cycling through all R barcodes for the 488-nm channel. The first and the last cycles were imaged with on-target barcodes. Scale bar, 50 μm. **b**, A composite on-target cycleHCR image of the three proteins in a nucleus. Scale bar, 10 μm. **c**, Automated image analysis pipeline for 3D nucleus segmentation and calculation of normalized mean fluorescence intensity above background. Scale bar, 10 μm. **d,e,** Mean fluorescence intensity above background was calculated within 575 nuclear masks from 8 fields of view for the 561-nm channel (**d**) and the 488-nm channel (**e**). The mean signal level of each cycle was normalized, setting the first cycle’s value to 100. Signal levels of the off-target cycles are up to 3.6% of the first cycle for the 561-nm channel and 0.77% of the first cycle for the 488-nm channel, while remaining low (< 0.15%) in the 640 nm channel (Fig. 1c). This suggests that the increasing signal level in the off-target cycles is due to autofluorescence. Error bars are 95% confidence intervals.

**Extended Data Fig. 2.**
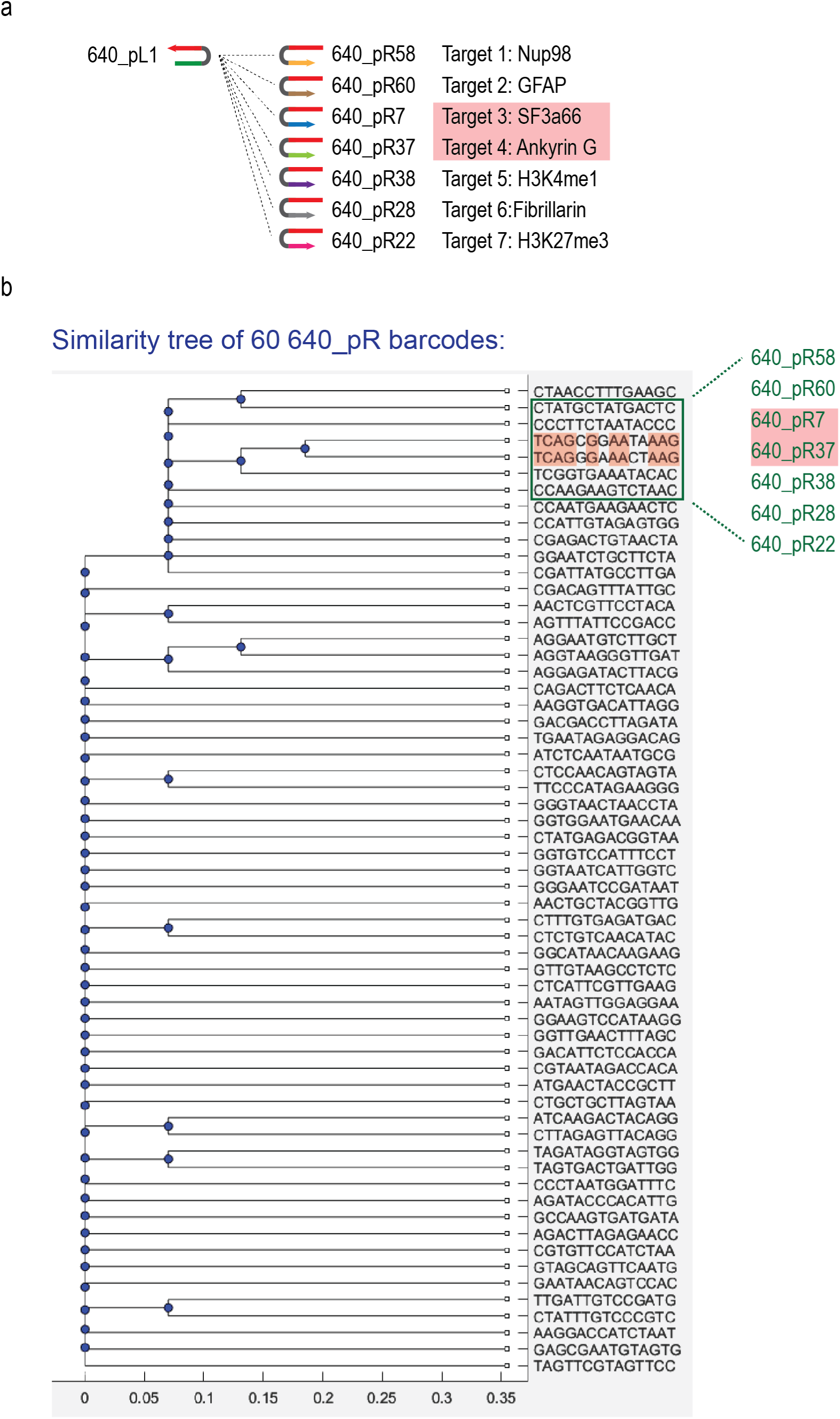
Evaluation of cycleHCR barcode sequence similarity for barcoding specificity test. **a,** Seven protein markers barcoded using the same L barcode and different R barcodes imaged in a mouse hippocampal section (Fig. 2d–h). **b**, The most similar R barcodes were used for barcoding the 7 protein targets in mouse hippocampus. Similarity of the 60 14-nt R barcodes for the 640-nm channel was evaluated using Matlab seqpdist() function with the jukes-cantor method and the phylogenetic tree was constructed by using the phytree() function. Two highly similar R barcodes (640_pR7 and 640_pR37) are highlighted; these sequences differ by only 4 nucleotides.

**Extended Data Fig. 3.**
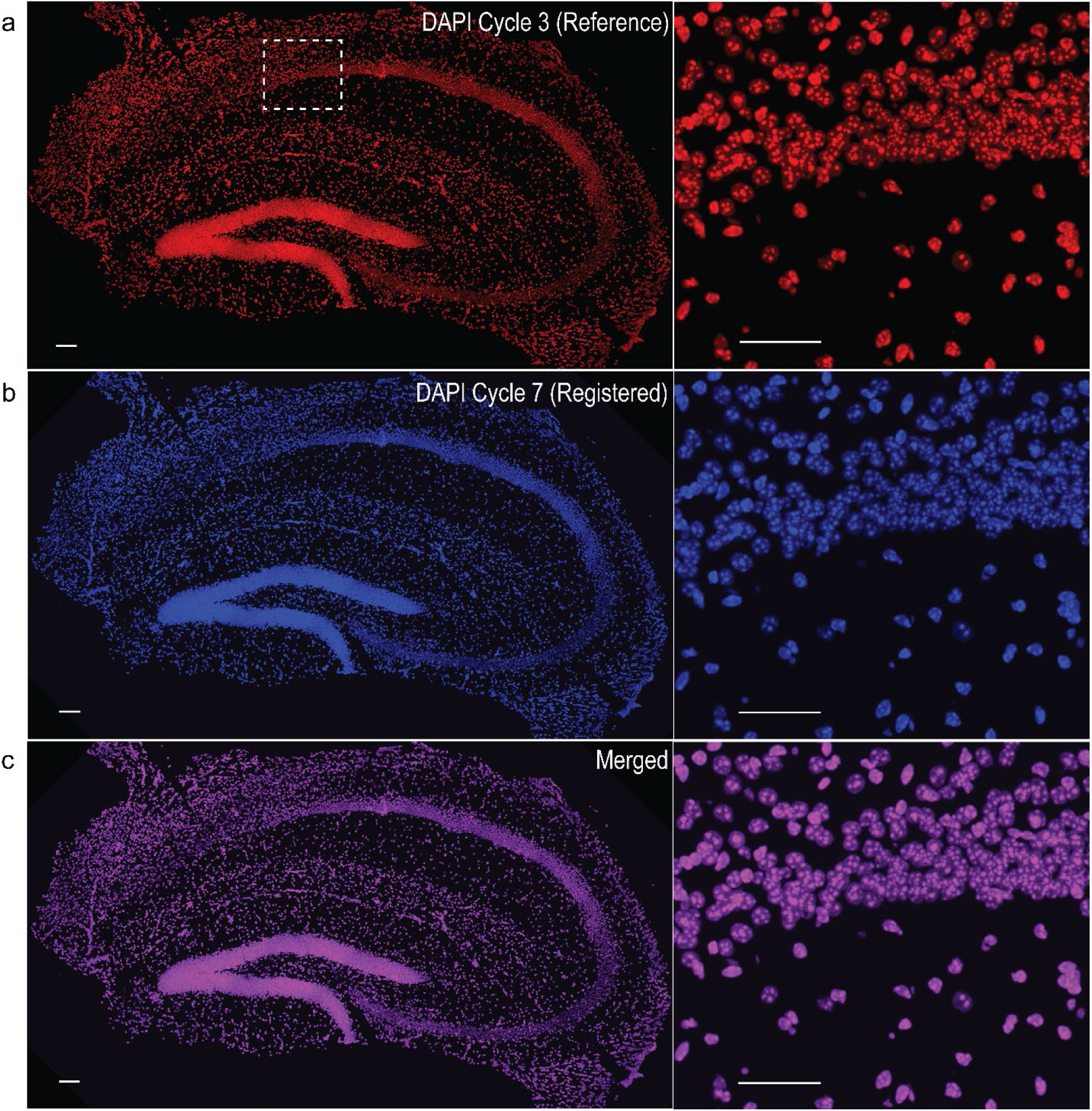
Validation of cross-cycle registration for protein cycleHCR imaging in mouse hippocampal section. Seven proteins were imaged sequentially through 7 cycles in the 640-nm channel. Cross-cycle registration was performed on DAPI images using the Bigstream pipeline. **a**, Stitched DAPI image from Cycle 3 was used as the reference for registration. **b–c**, Registered DAPI image of the last cycle (Cycle 7, **b**) is merged with the reference image (**c**). Images are 3D views of the 40-μm hippocampal section produced by Imaris. Zoomed-in views of the boxed region are in the right panel. Scale bars, 100 μm (left), 50 μm (right).

**Extended Data Fig. 4.**
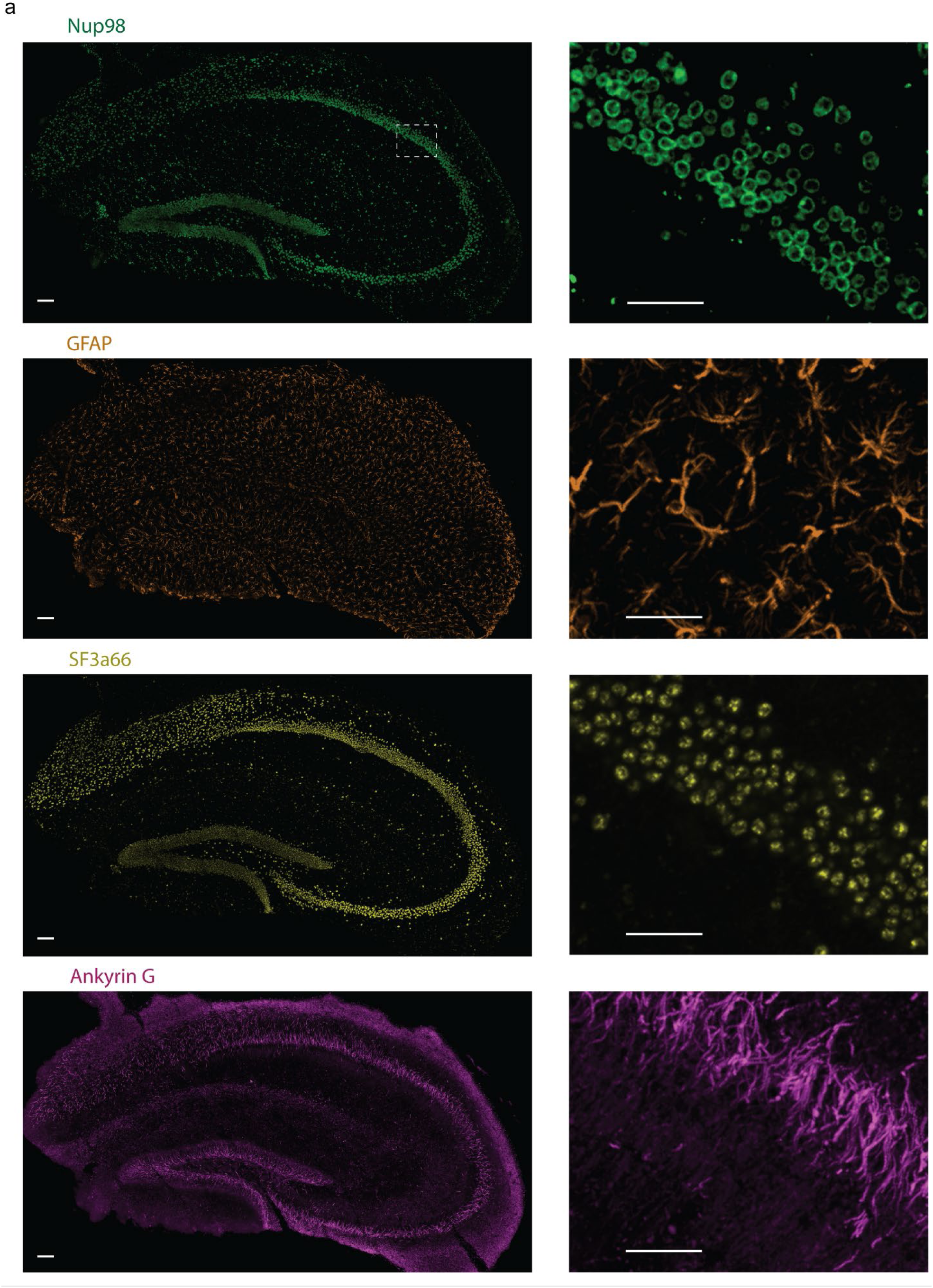
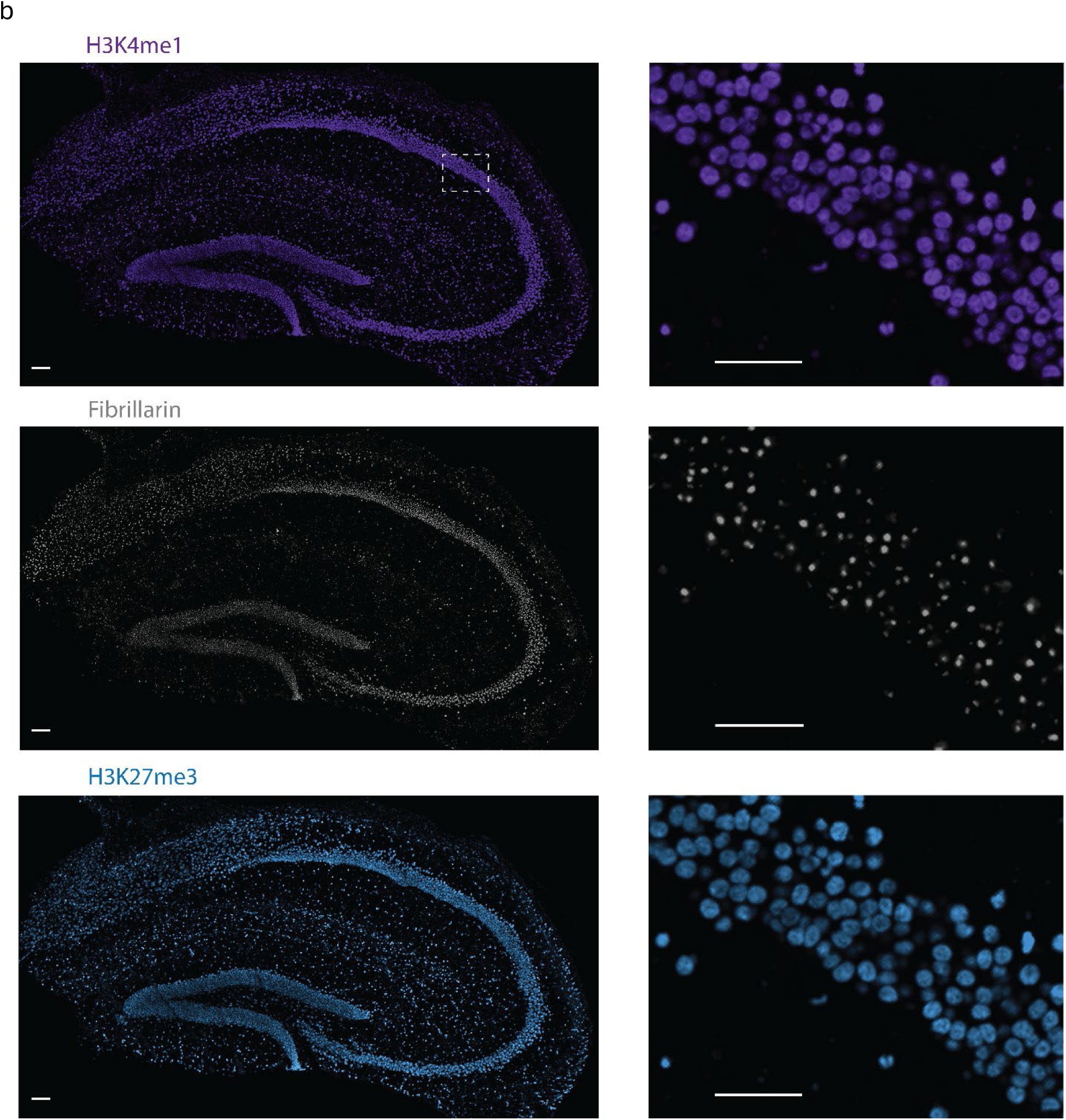
Seven proteins imaged in cycleHCR barcode specificity test in mouse hippocampus. The proteins were imaged using the same L barcode and the most similar R barcode sequences as listed in Extended Data Fig. 2. 3D views of stitched and registered images of the 40-μm mouse hippocampus section were rendered by Imaris. Left panels display the whole hippocampus section. Right panels are zoomed-in views of the boxed region. **a**, Nup98 (nuclear pore), GFAP (Glial Fibrillary Acidic Protein), SF3a66 (spliceosome), Ankyrin G (axon initial segment), **b**, H3K4me1 (enhancer region), fibrillarin (nucleolus fibrillar center), and H3K27me3 (heterochromatin) Scale bars, 100 μm (left), 50 μm (right).

**Extended Data Fig. 5.**
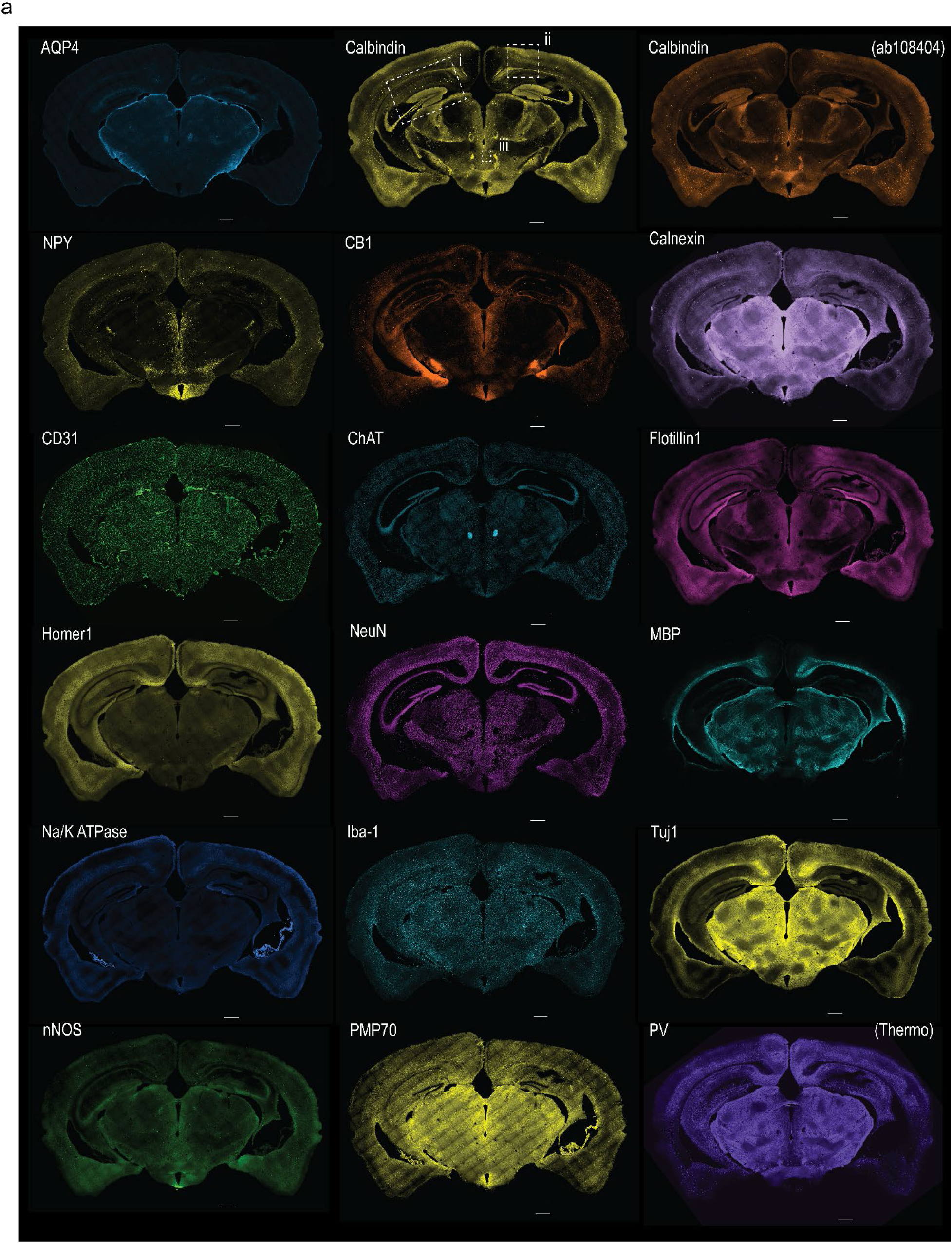
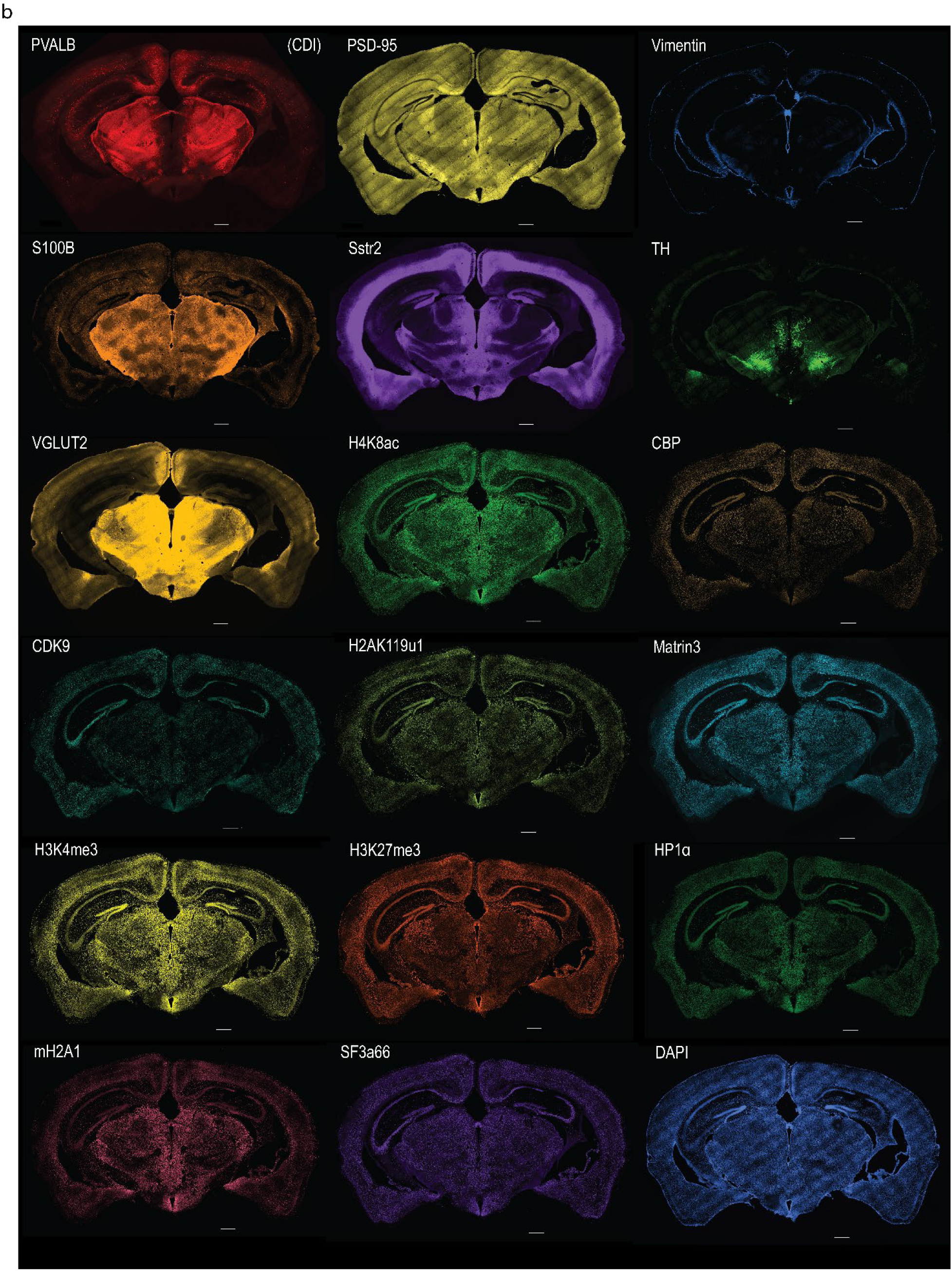
Protein images acquired by multiplex cycleHCR of 46 proteins and 79 RNAs in a mouse whole brain section. **a–b,** Representative images displaying protein expression patterns in Brain Section 1. 3D images of the 40-μm mouse whole brain section were rendered in Imaris. In the middle image on the top row in **a**, the boxes indicate the regions visualized in the following figures (i: Extended Data Fig. 6; ii: Extended Data Fig. 7; iii: Extended Data Fig. 8). Scale bars, 500 μm.

**Extended Data Fig. 6.**
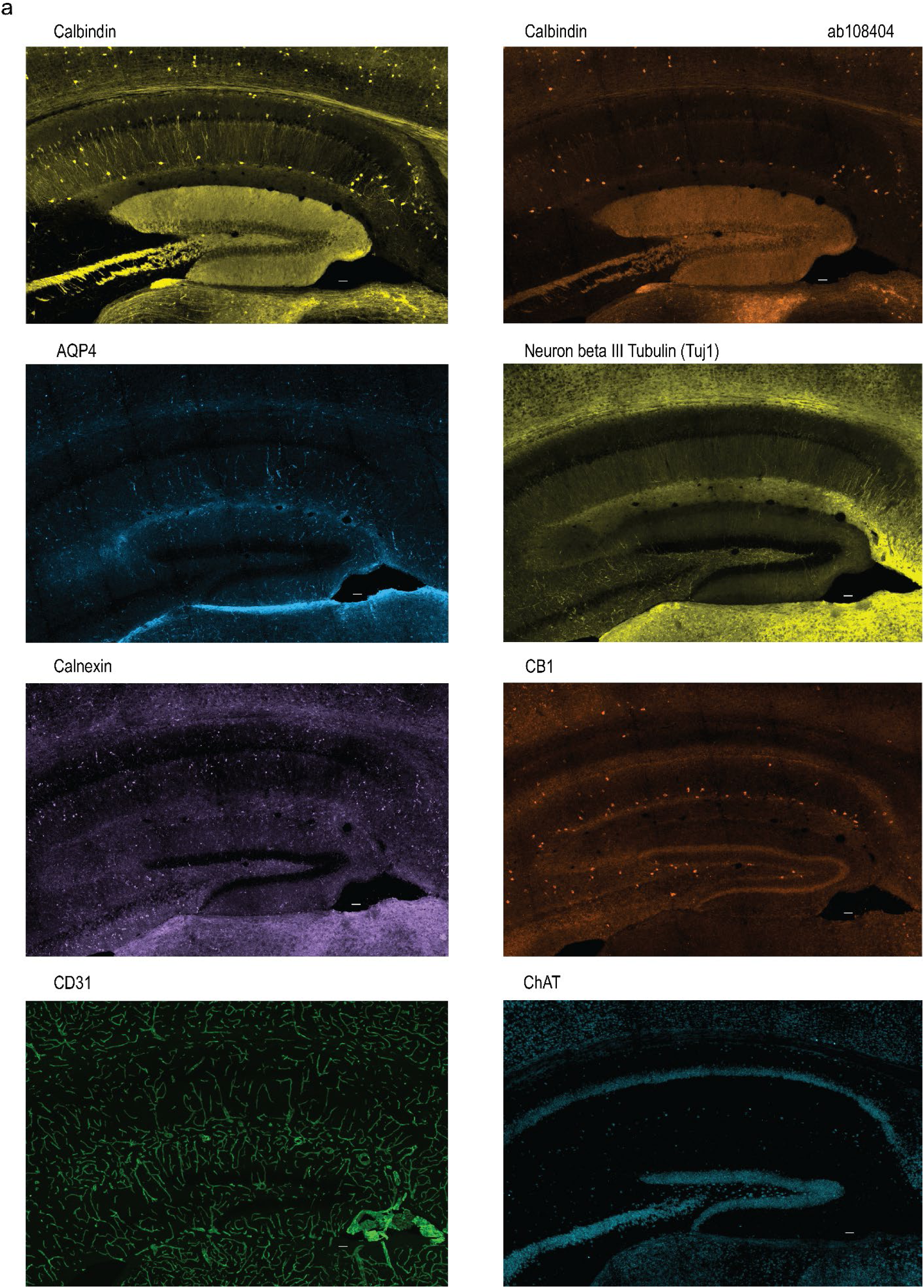
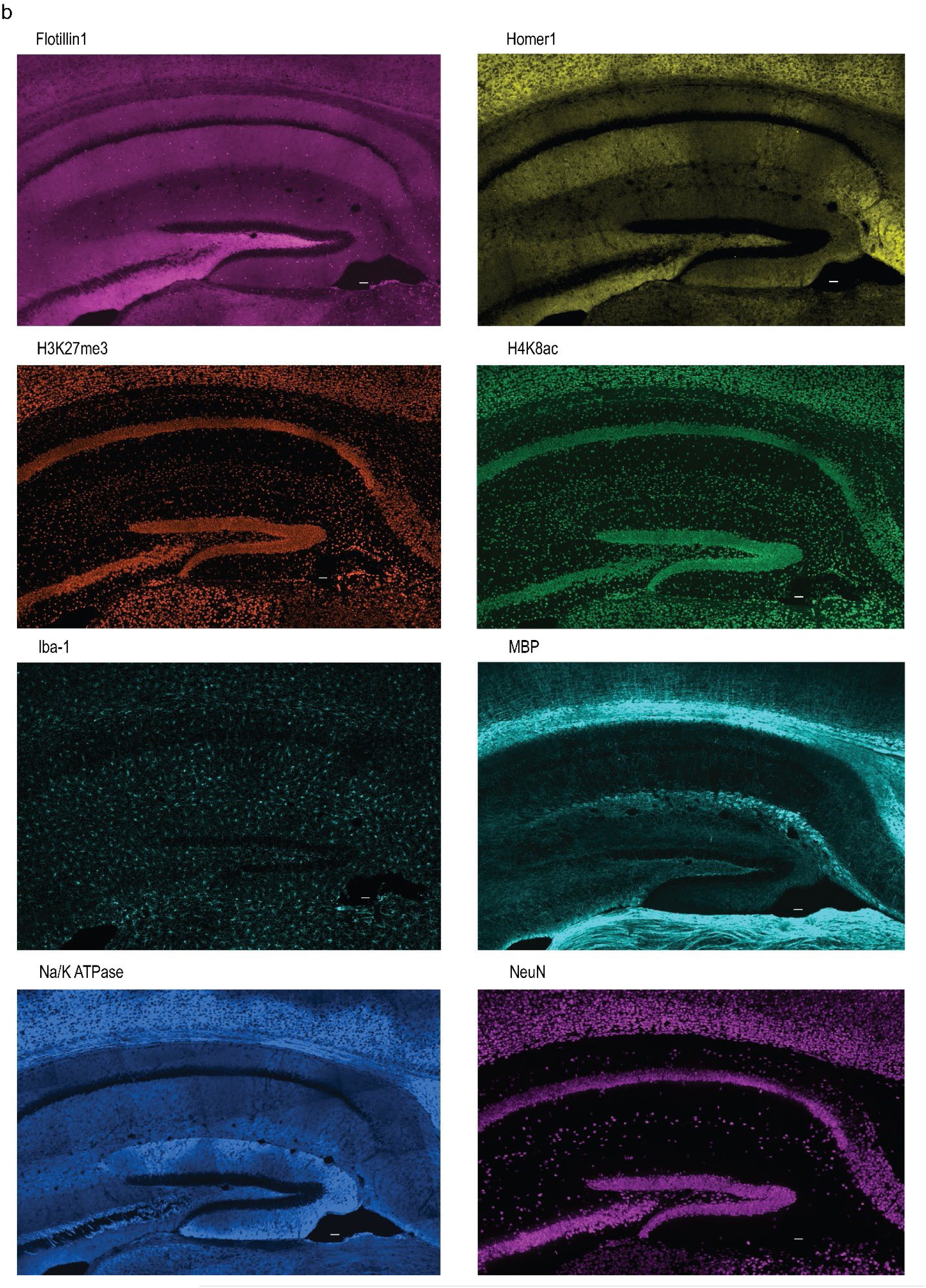
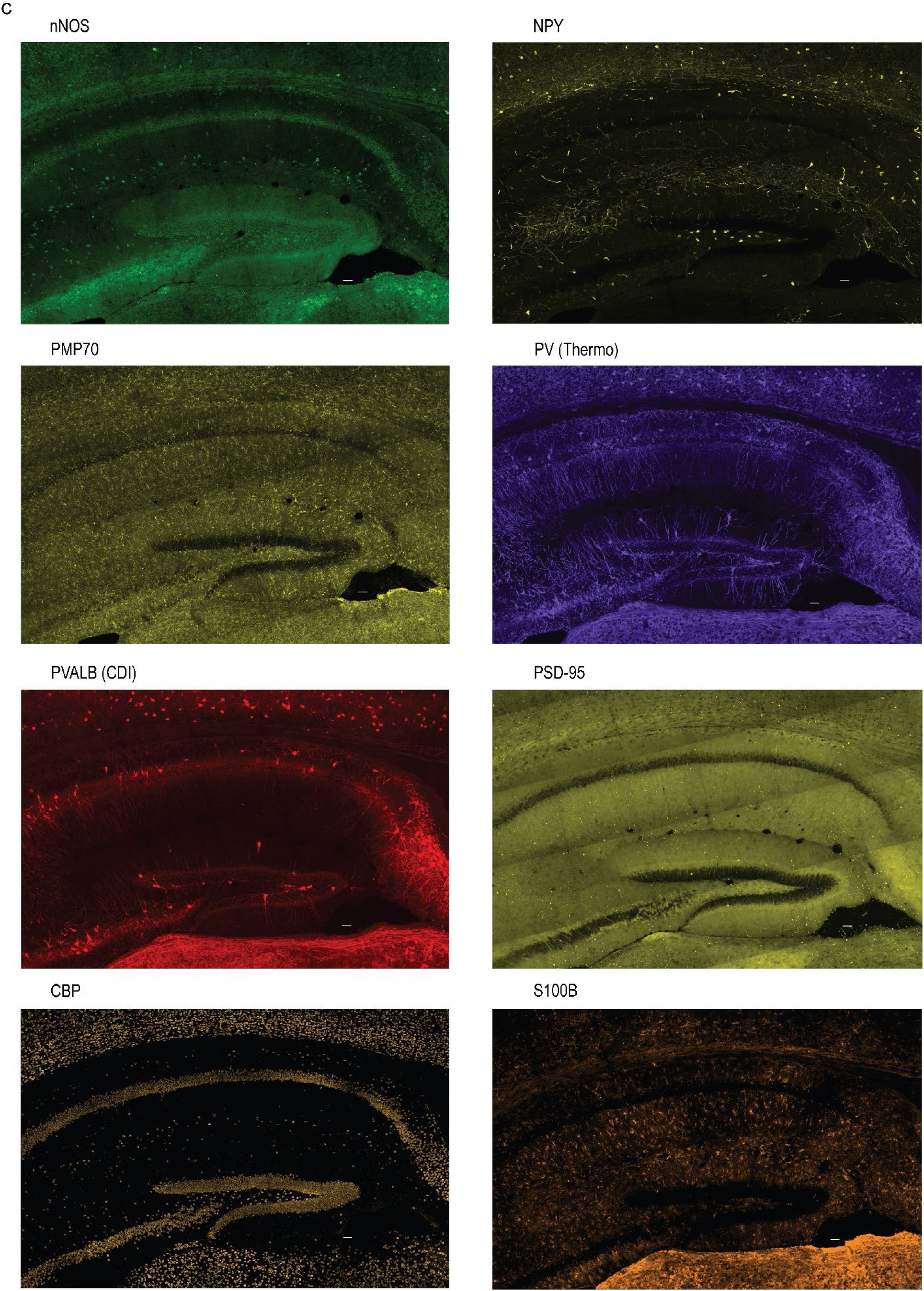
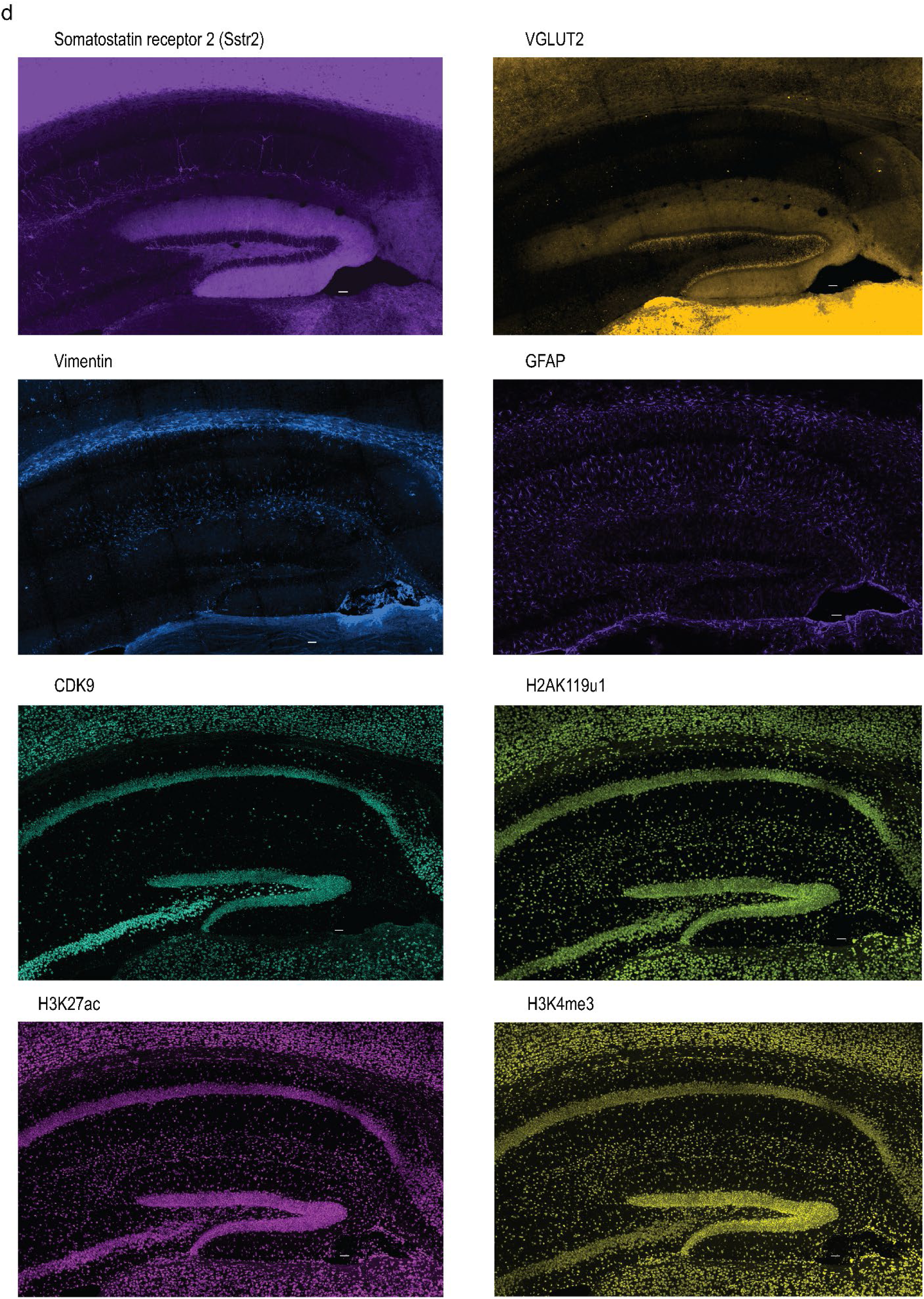
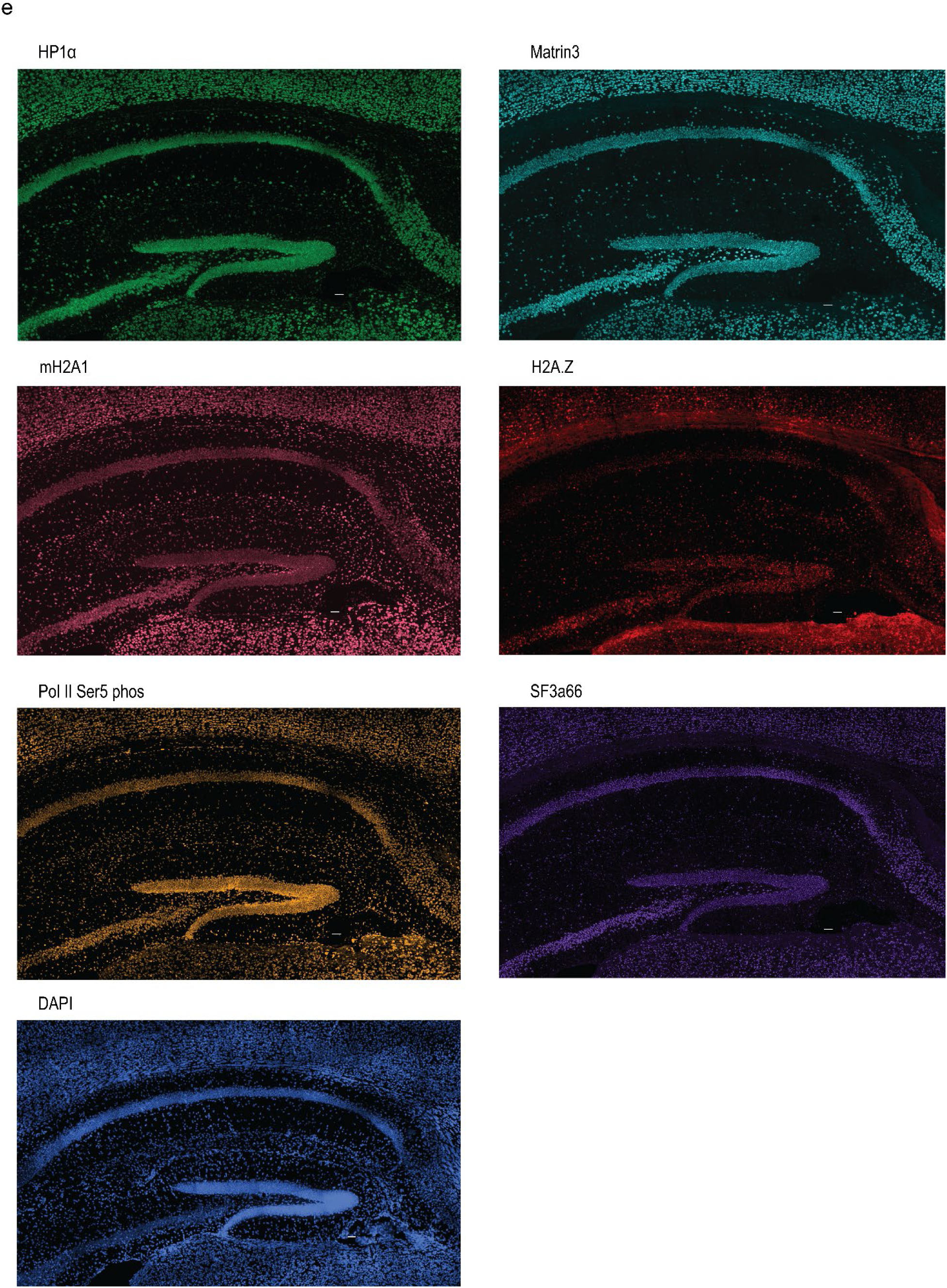
Enlarged hippocampal views of protein images from multiplex protein and RNA cycleHCR. **a–e,** Representative images were chosen to display protein expression patterns in the hippocampus. 3D images were rendered using Imaris. Images shown in this figure are zoomed-in views of the boxed region (i) in Extended Data Fig. 5. Scale bars, 50 μm.

**Extended Data Fig. 7.**
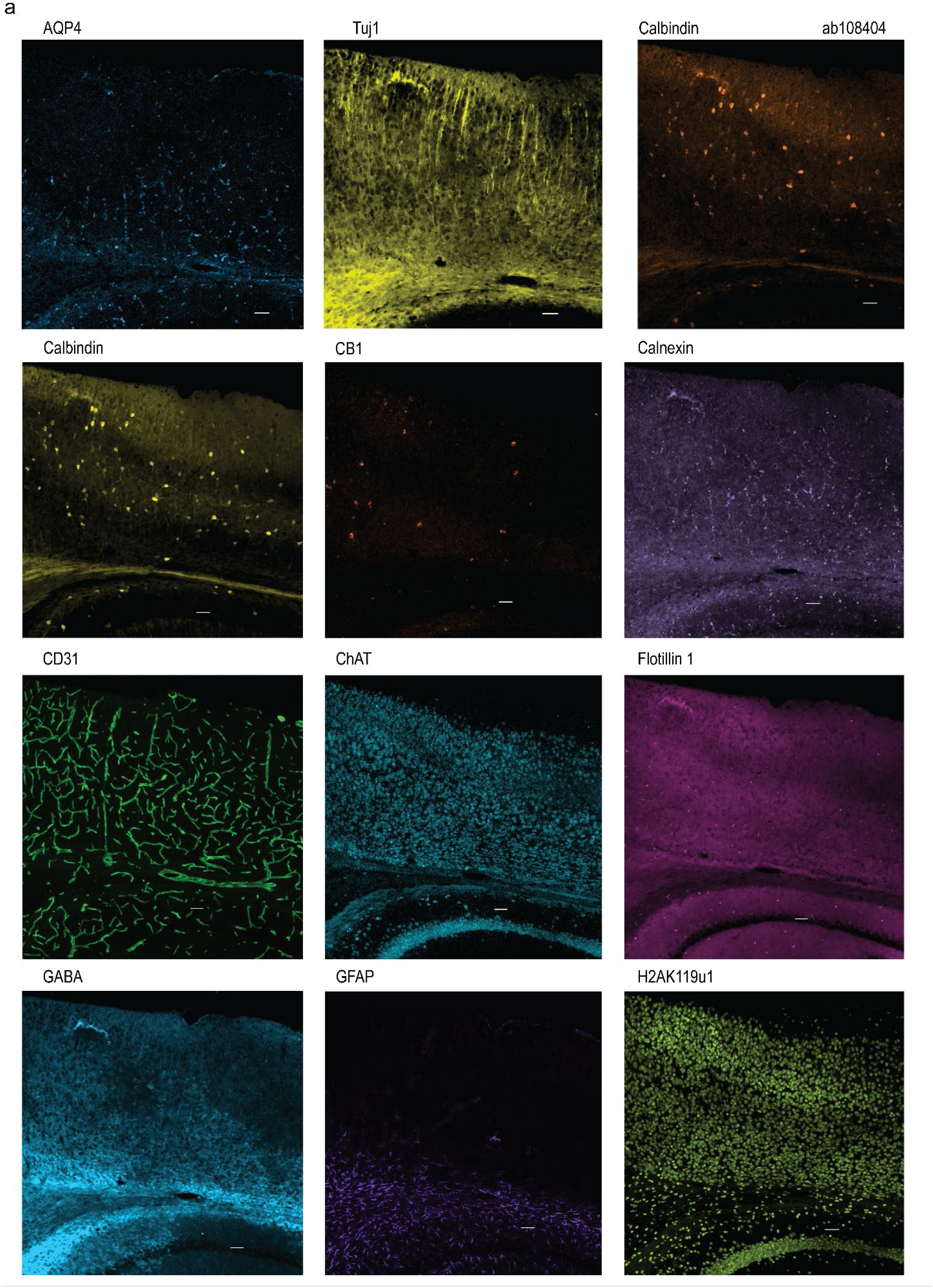
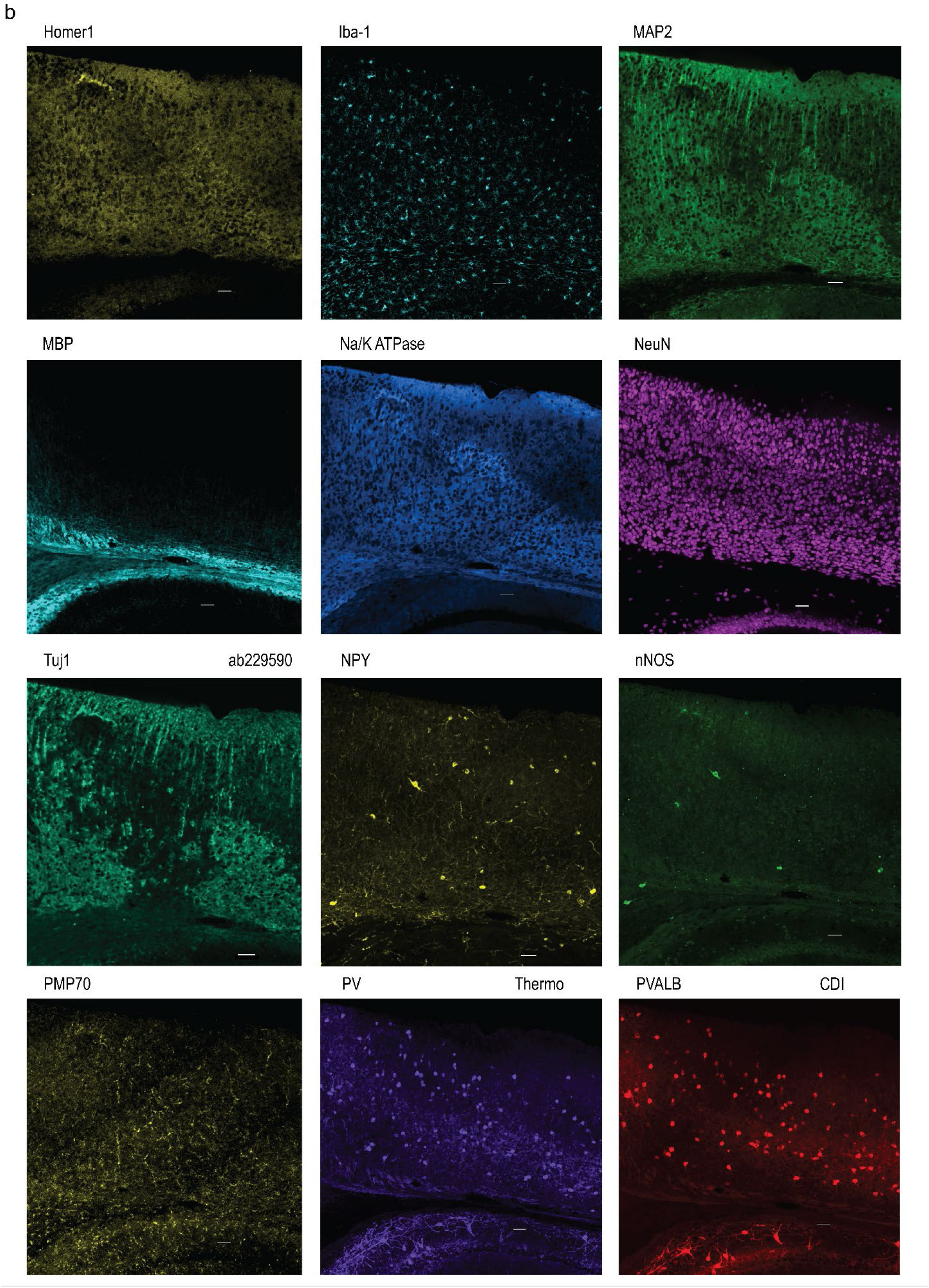
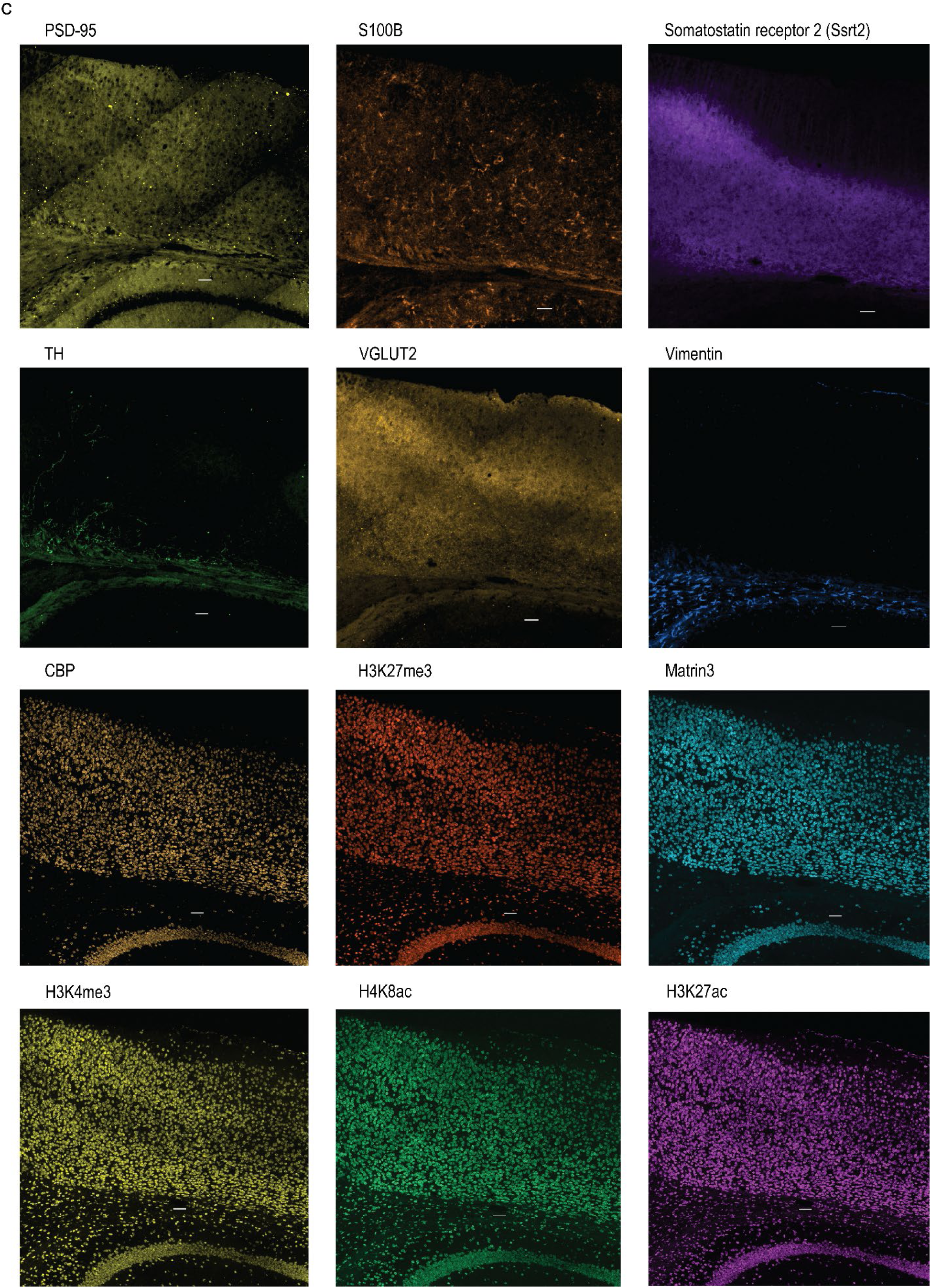
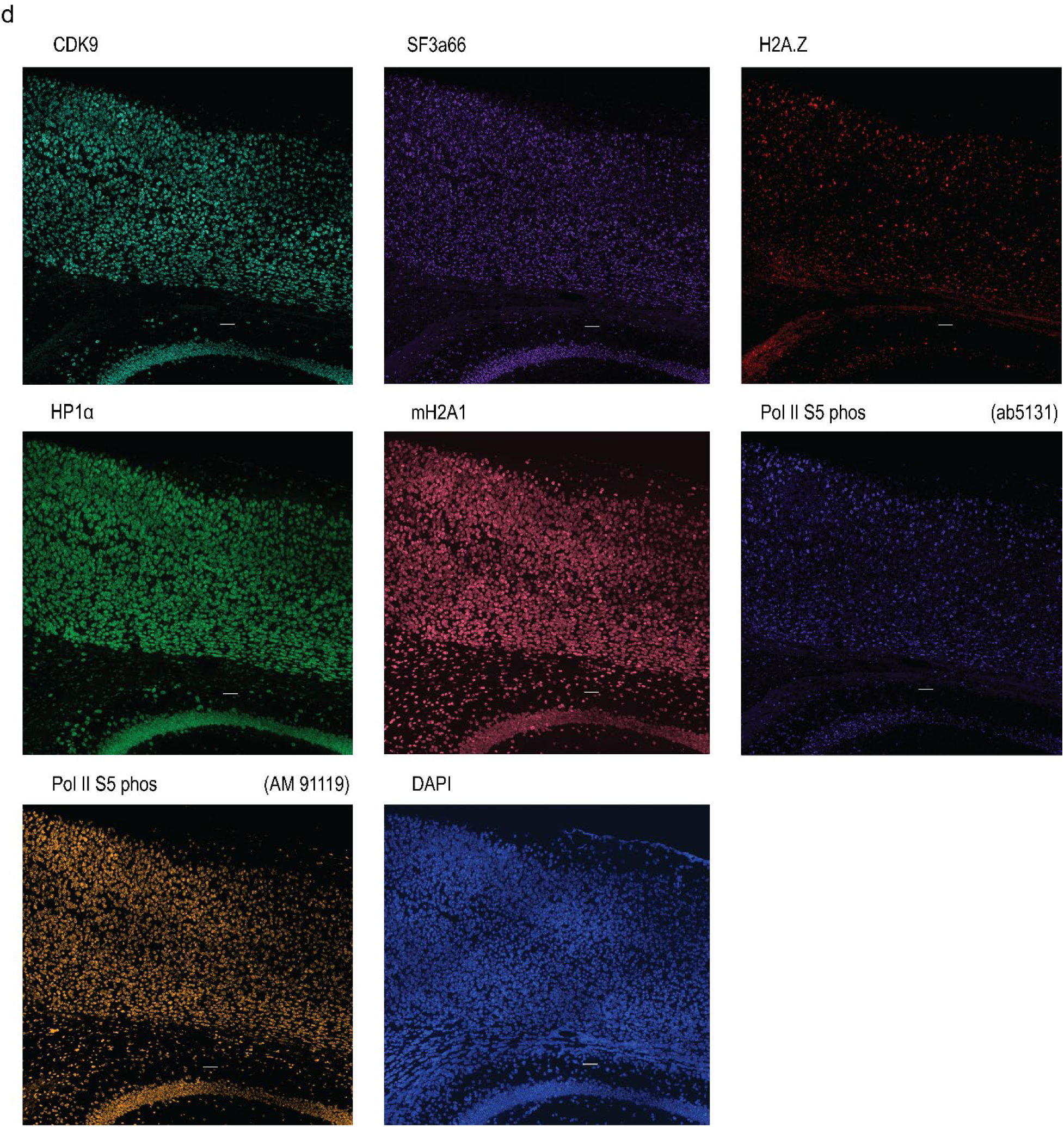
Enlarged views of a cortical region showing protein images acquired by multiplex protein and RNA cycleHCR. **a–d,** Representative images were chosen to display protein expression patterns in the cortical region. Images shown in this figure are zoomed-in views of the boxed region (ii) in Extended Data Fig. 5. Scale bars, 50 μm. 3D images were rendered in Imaris.

**Extended Data Fig. 8.**
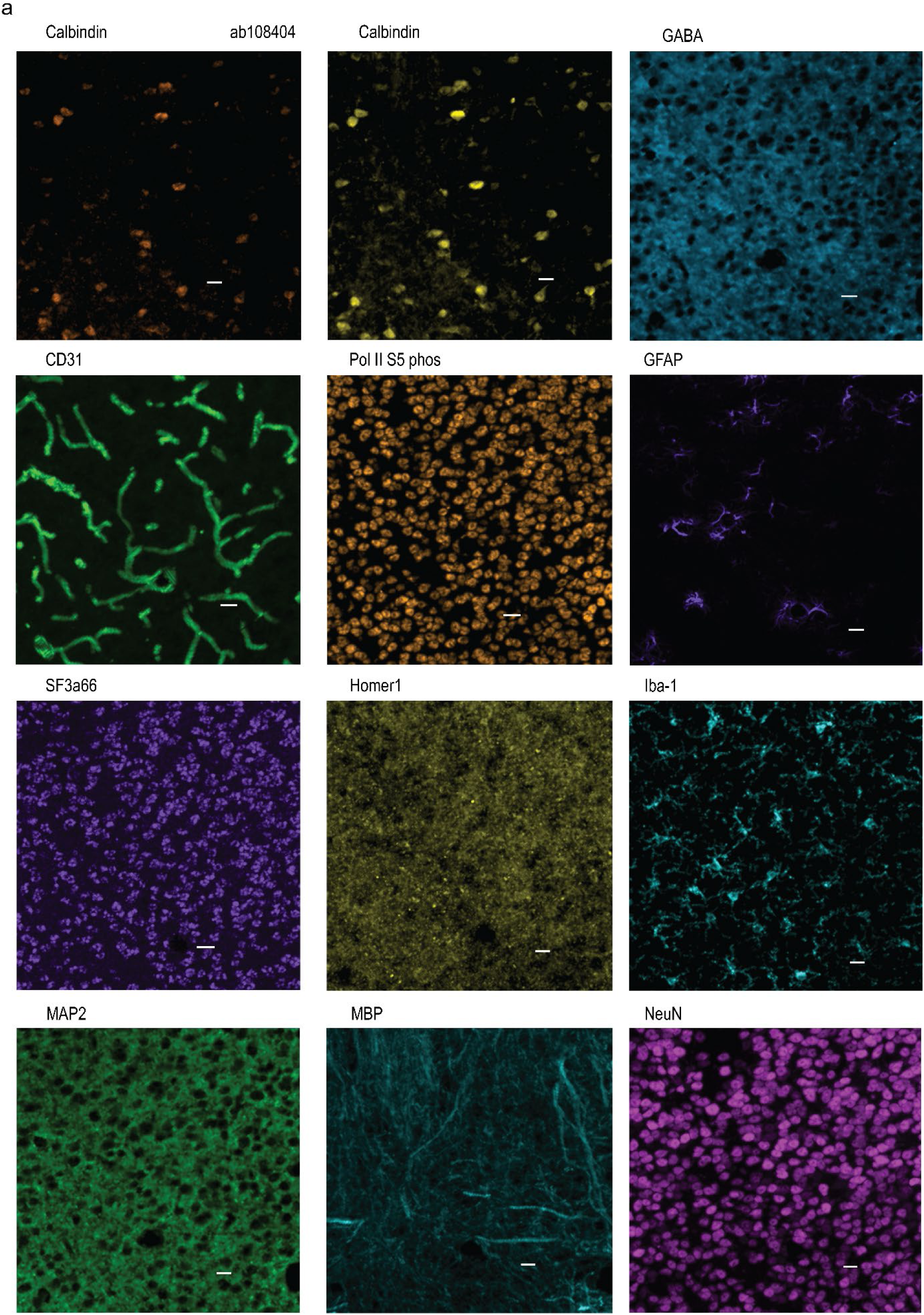
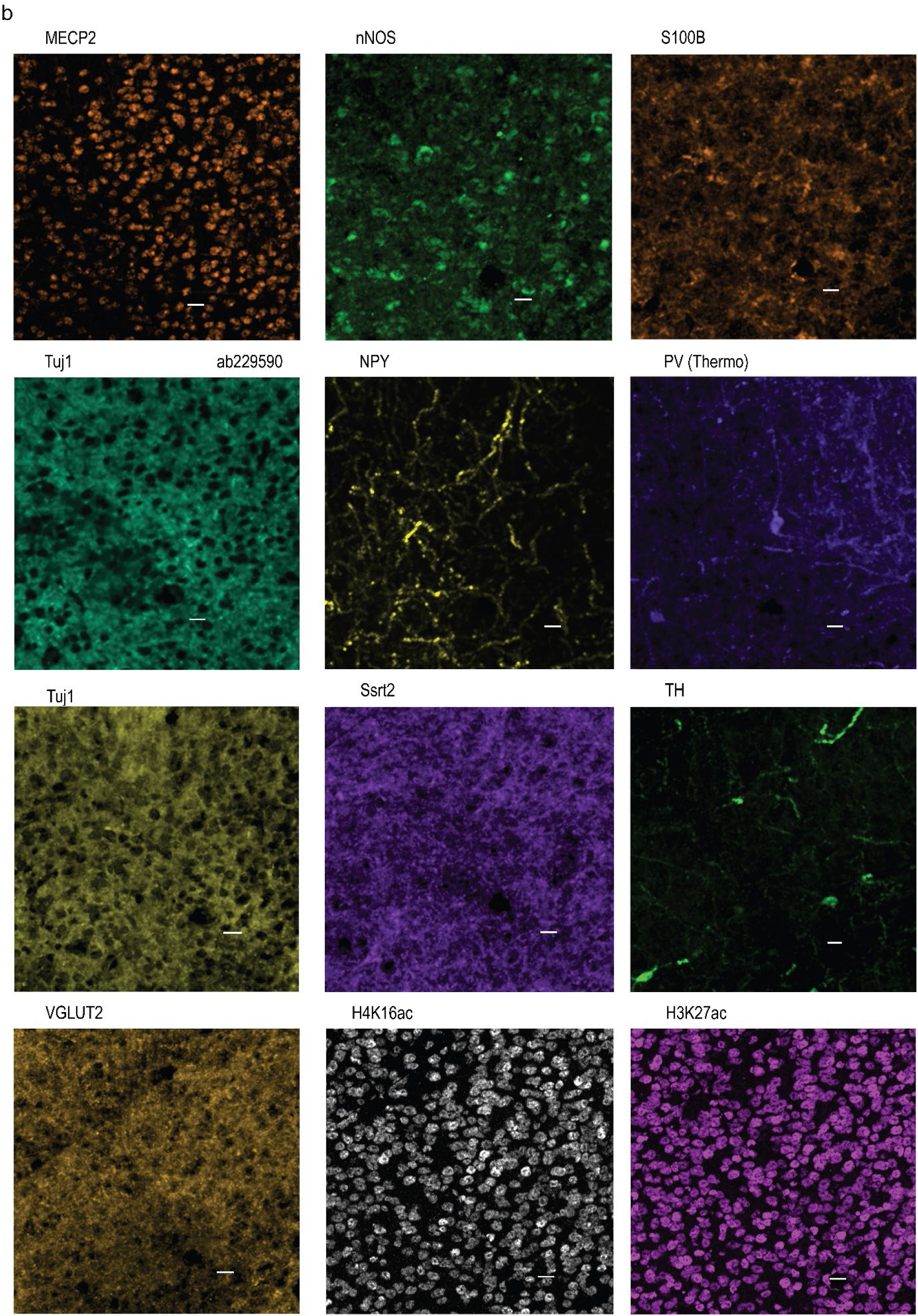
Enlarged view of a thalamic region showing protein images acquired by multiplex protein and RNA cycleHCR. **a–b,** Representative images were chosen to display protein expression patterns in a thalamic region. Images shown in this figure are zoomed-in views of the boxed region (iii) in Extended Data Fig. 6. Scale bars, 20 μm. 3D images were rendered in Imaris.

**Extended Data Fig. 9.**
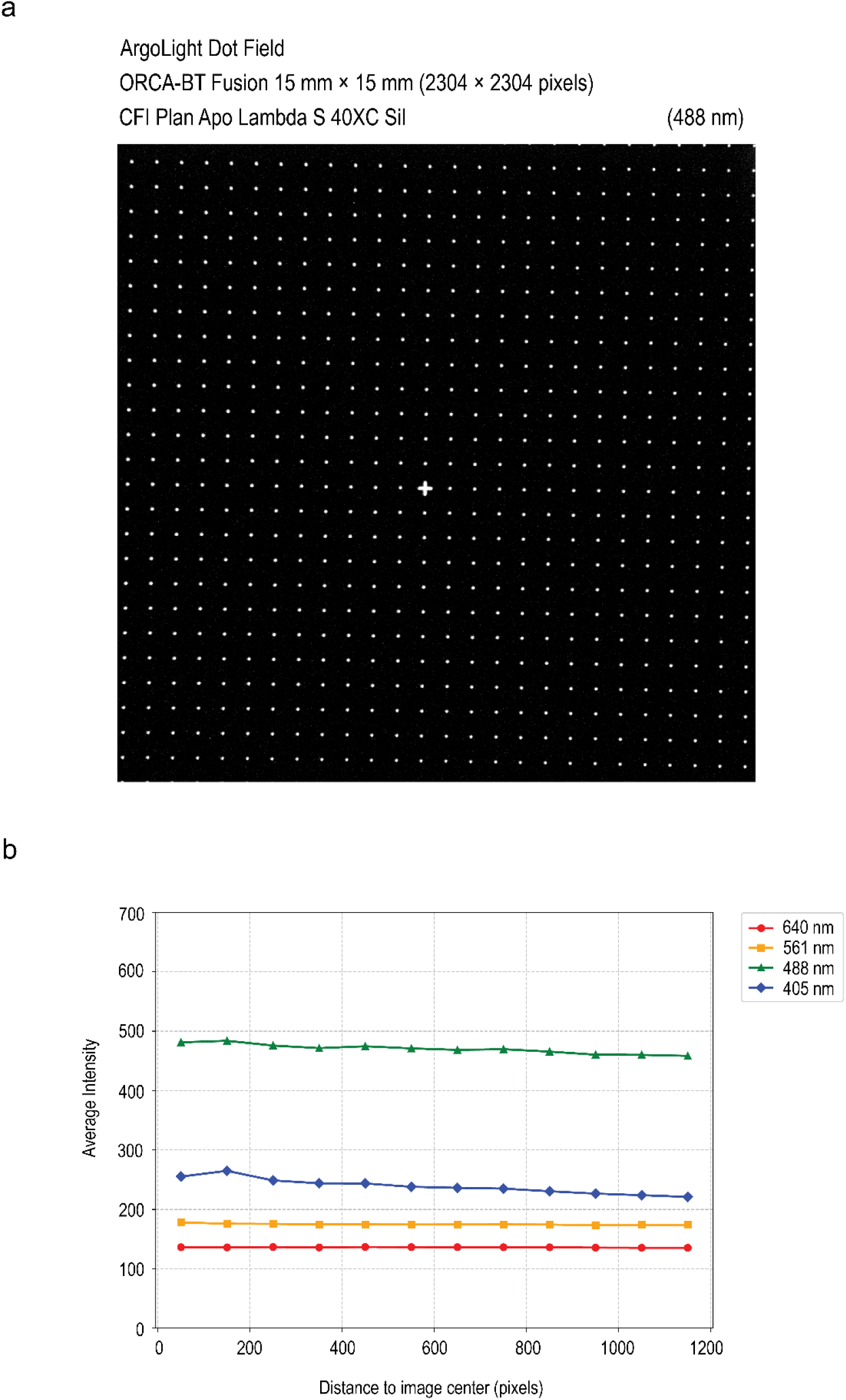
Evaluation of illumination and field uniformity. **a**, Representative full chip images of the ArgoLight dot field obtained using a 40x silicone oil immersion objective with 488 nm laser illumination. **b**, Illumination uniformity was evaluated by calculating the average dot intensity, plotted as a function of distance from the camera chip center, for different laser wavelengths: 405 nm, 488 nm, 561 nm, and 640 nm.

**Extended Data Fig. 10.**
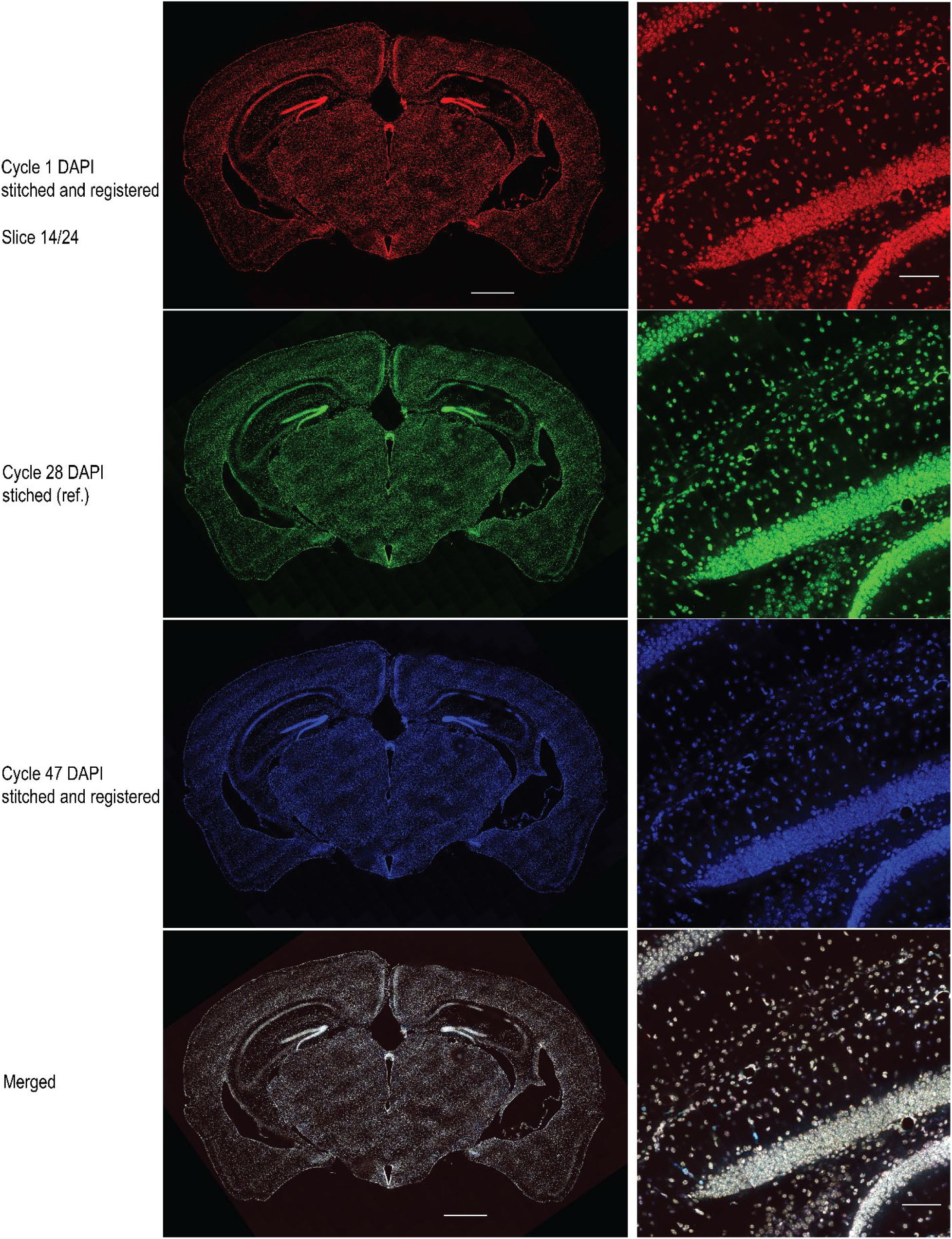
Cross-cycle registration validation for combined protein and RNA cycleHCR in mouse brain section. Stitched whole-brain 3D DAPI image acquired from each imaging cycle was used for cross-cycle registration. The alignment of the reference DAPI image (Cycle 28) with DAPI images from the first and the last cycle, Cycle 1 and Cycle 47, were overlaid together (last row). A zoomed-in view in the hippocampus shows a detailed examination of image registration. Scale bars, 1 mm (left), 100 μm (right).

**Extended Data Fig. 11.**
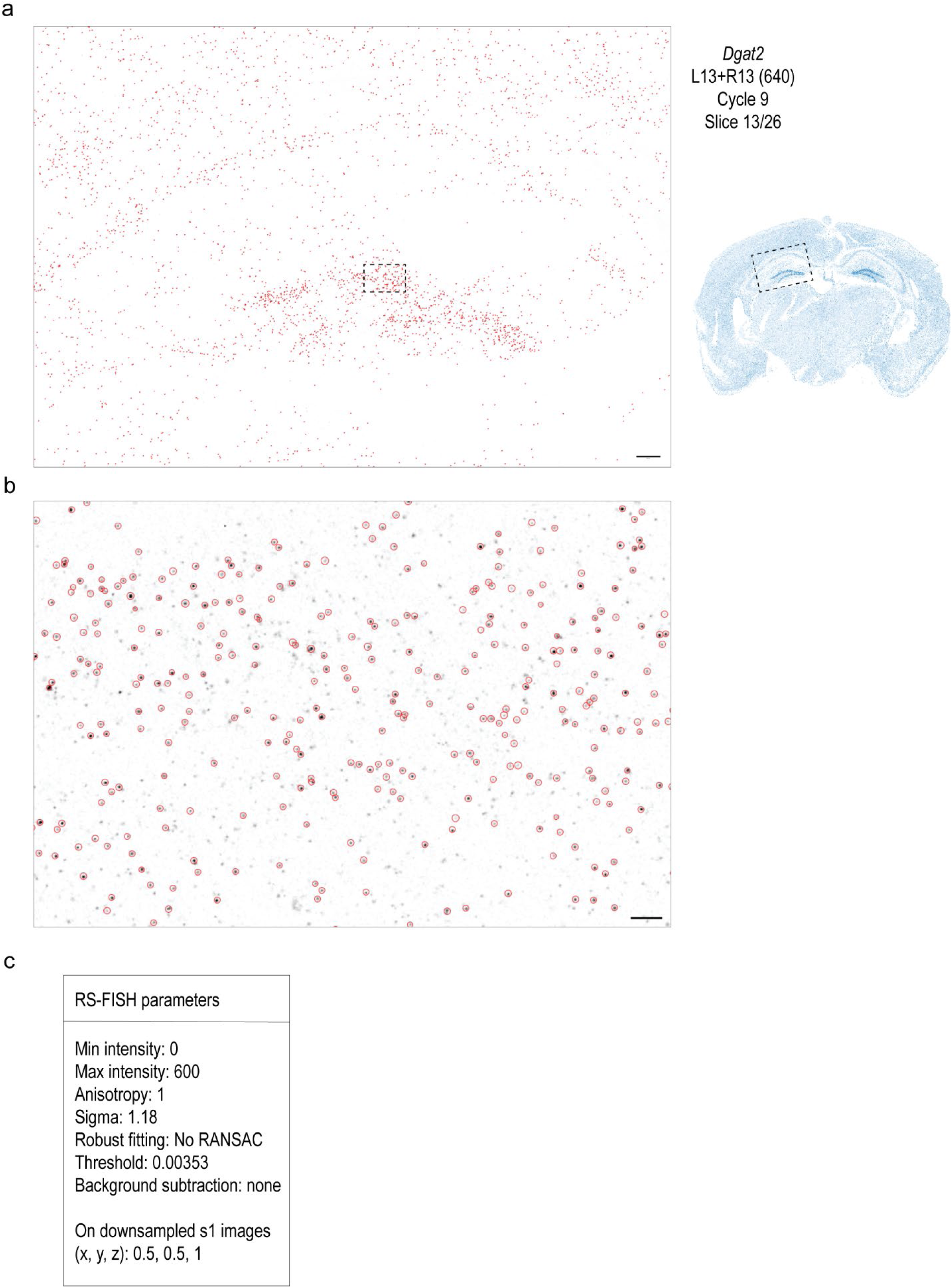
RNA spot detection using RS-FISH for joint protein and RNA cycleHCR analysis. **a**, Enlarged hippocampal region from a *Dgat2* gene image slice showing detected RNA spots. Red circles overlaid on an intensity-inverted, downsampled s1 image [sampling factors in (x, y, z): 0.5, 0.5, 1] indicate the RNA spots identified by RS-FISH. Scale bar, 100 μm. **b**, A zoomed-in view of the boxed region in **a** shows detailed examination of detected spots. Scale bar, 10 μm. **c**, RS-FISH parameters were carefully chosen to reduce false positive detections and to yield accurate spot localization. The same parameters were applied for RNA images in both replicates of mouse brain sections.

**Extended Data Fig. 12.**
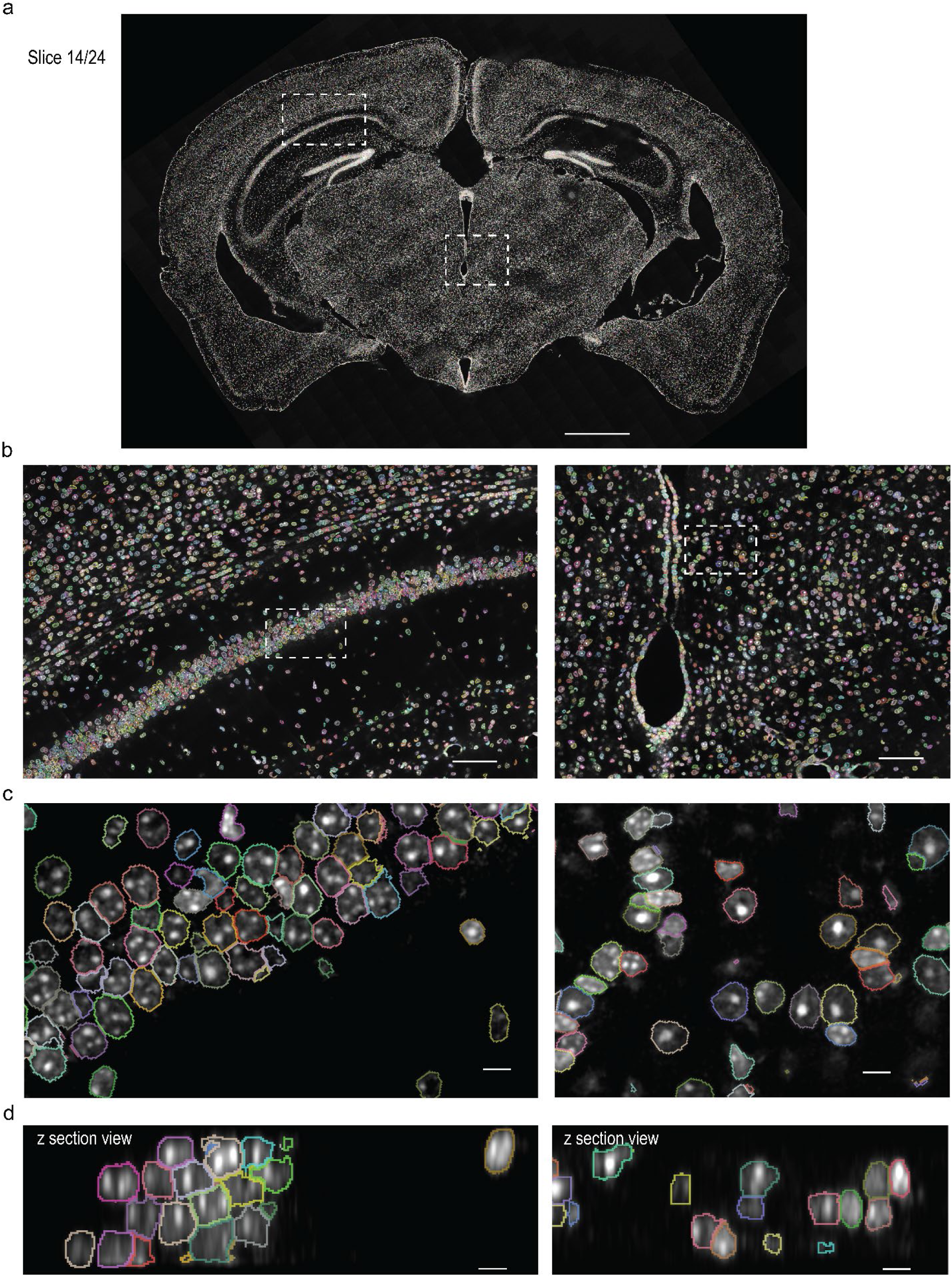
3D segmentation of nuclei in a mouse whole brain section using DAPI images. Individual nuclei in the stitched, registered whole brain DAPI image were segmented in 3D for downstream nuclear protein analysis and RNA spot-to-cell assignment. Cellpose with distributed cores was implemented to handle the large image datasets. The ‘cyto3’ model was used. Around 200,000 nuclei were segmented from each 40-μm whole-brain section. **a**, A slice view of segmented whole brain DAPI image is displayed. Scale bar, 1 mm. **b,c**, Hierarchical zoomed-in xy views in the box regions in (a, b) at different scales show closer examination of segmentation. Scale bars, 100 μm (b), 10 μm (c). **d**, Z-section views demonstrate that the segmentation captures nuclear structure in 3D. Scale bars, 10 μm. Images were rendered in Dragonfly.

**Extended Data Fig. 13.**
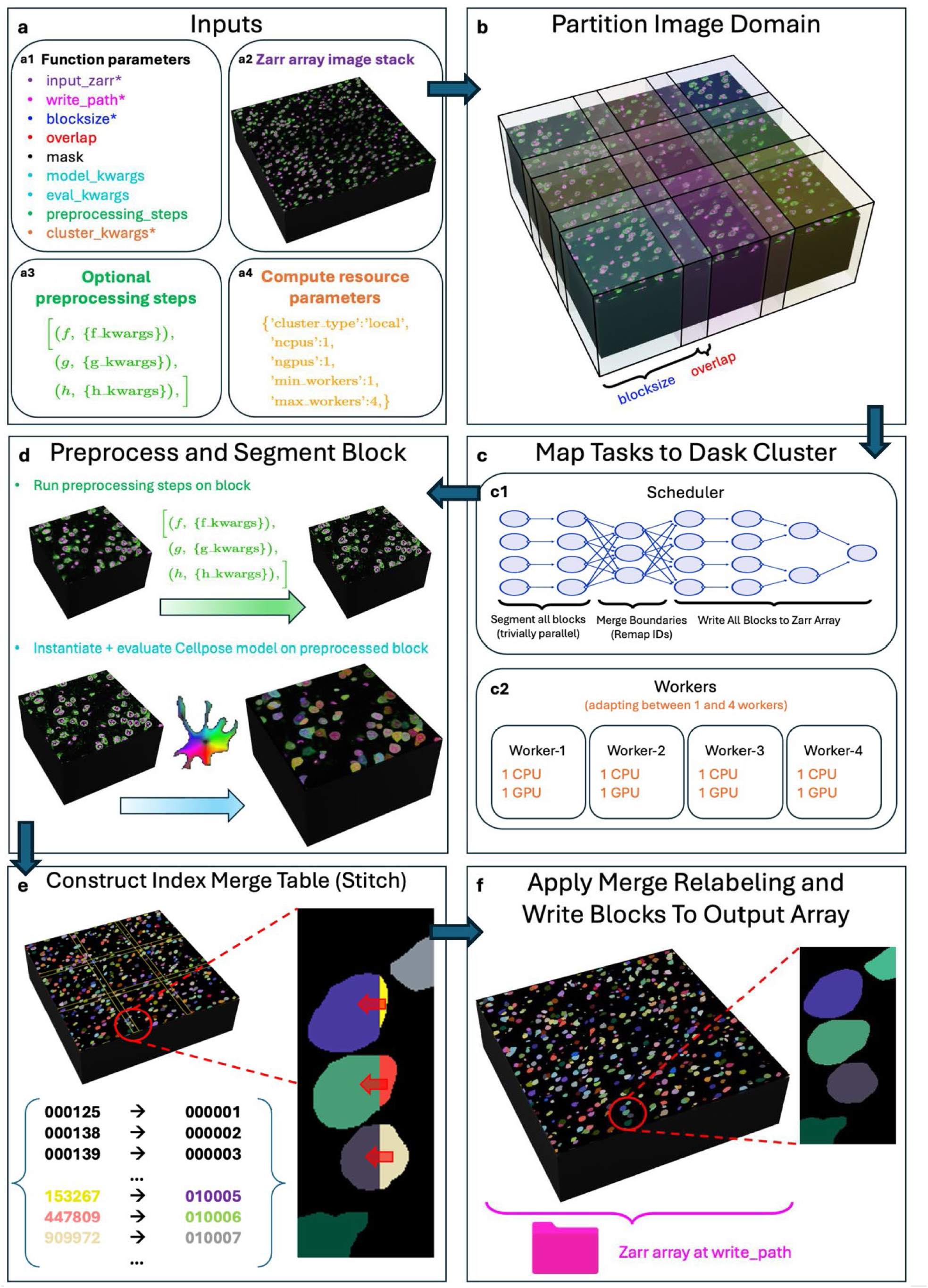
Distributed Cellpose for cell segmentation on large datasets. The distributed Cellpose module scales Cellpose^19^ to arbitrarily large 3D datasets by processing sub-regions (blocks) in parallel then stitching them to produce a seamless whole image segmentation. **a**, Distributed Cellpose is executed by a single function from which the user can control various aspects of the computation. **a1,** There are four required inputs (marked with stars) and several optional ones. **a2,** The image stack must be in a chunked big data format such as Zarr or N5. **a3,** If preprocessing must be done before Cellpose (e.g., smoothing or background subtraction), functions and their arguments can be provided; these will be executed on the blocks before Cellpose as part of the distributed computation. **a4,** The user must specify the compute resources available, e.g. whether working on a local workstation or cluster and how many CPUs and GPUs will be used for processing each block. **b,** Blocks overlap to ensure consistent segmentation across block seams. **c,** The computation is distributed using a Dask cluster^32^. **c1,** The computation proceeds in three general steps: preprocess and segment every block (see panel **d)**, merge cell segments across block boundaries (see panel **e**), and apply cell segment index remapping then write blocks to disk (see panel **f**). This is a classic map-reduce-map scheme where the first and third steps are trivially parallelizable, but the middle step requires sharing block boundary information across all workers. **c2,** The Dask cluster maps computations to a pool of workers, the number of which and their size, are controlled by the user. **d,** For each block, preprocessing steps then Cellpose are executed. The user can fully parameterize the Cellpose model (e.g., provide a path to a custom pre-trained model) and specify evaluation parameters. This is the most compute heavy step, but it is trivially parallelizable over all blocks. **e,** All seams between blocks are inspected and cell indices that should merge are recorded in a table. This is somewhat parallelizable, however ultimately a single process must construct one table that is globally consistent across the entire image domain. **f,** The index remapping table is shared with all workers. Block-local cell indices are remapped to the globally consistent indices then the block is written to disk. This step is also trivially parallelizable.

**Extended Data Fig. 14.**
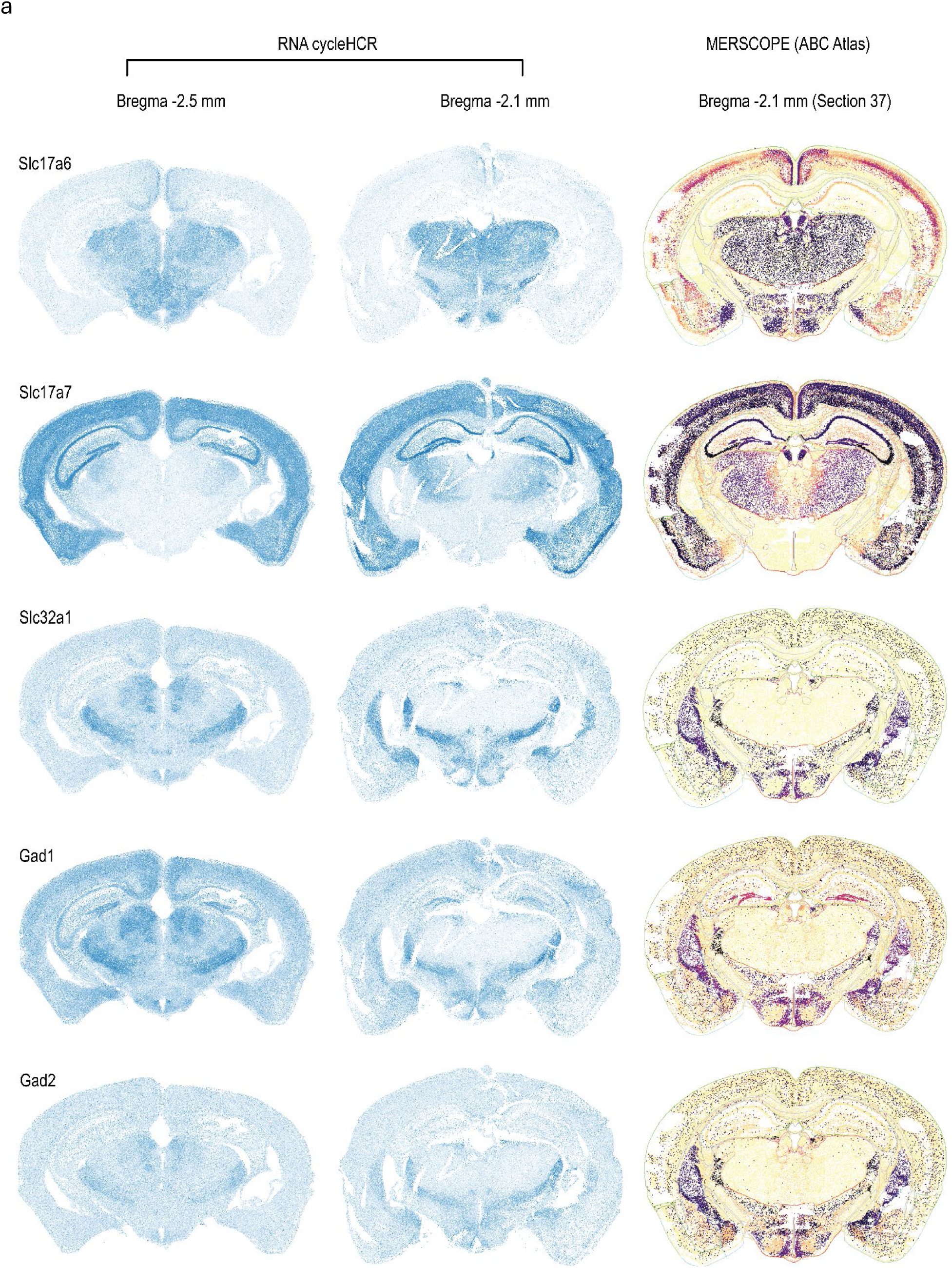
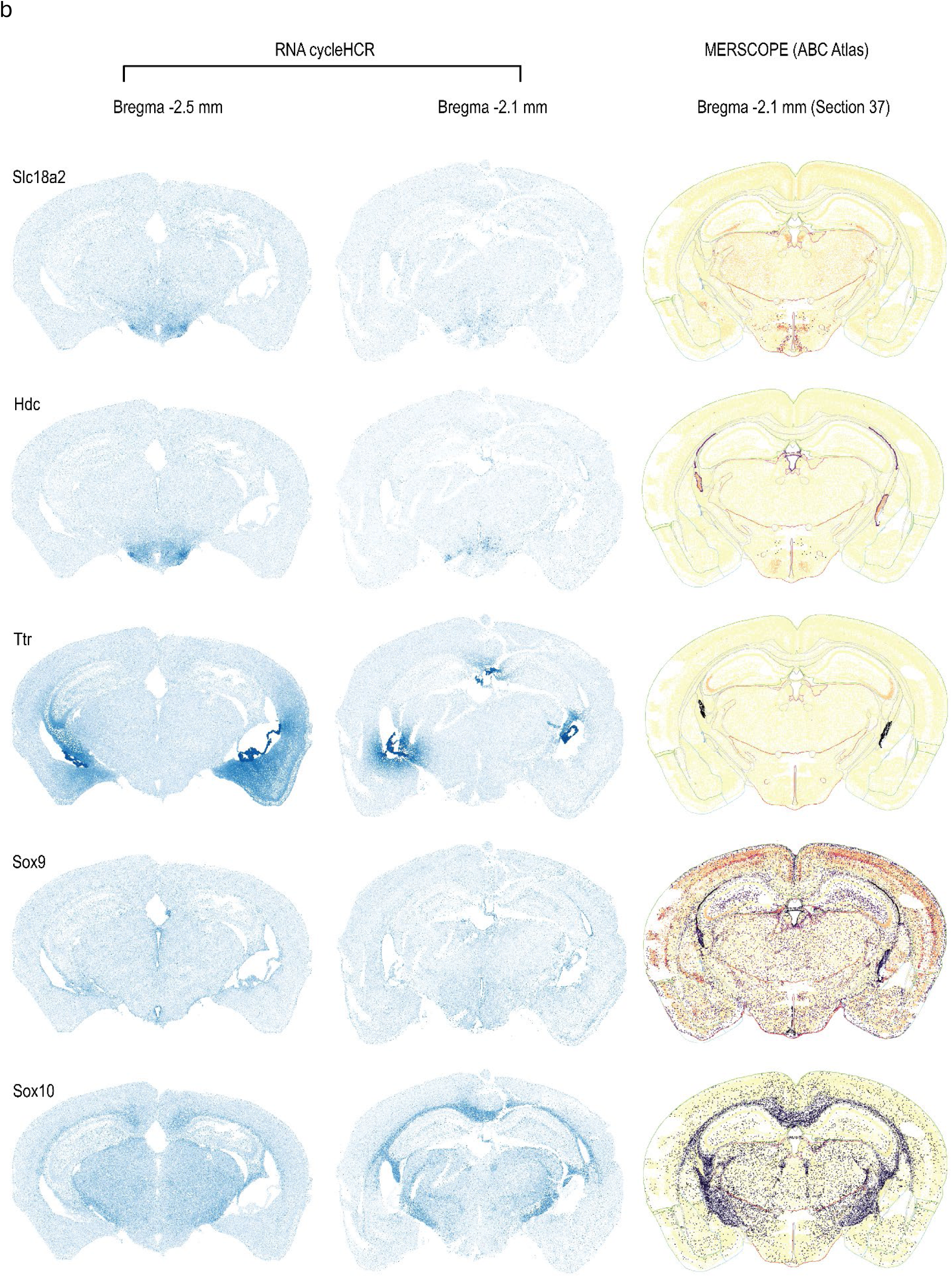
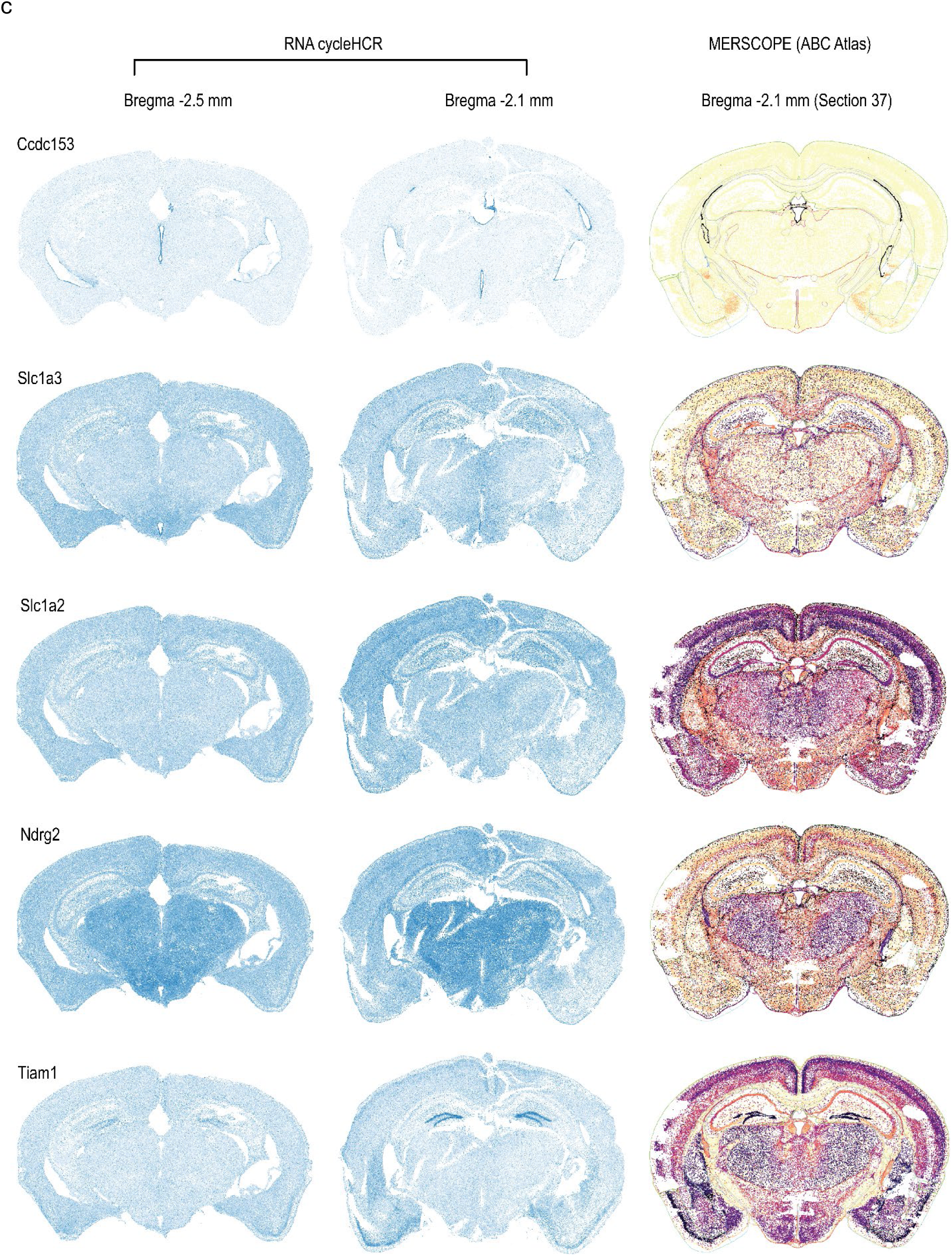
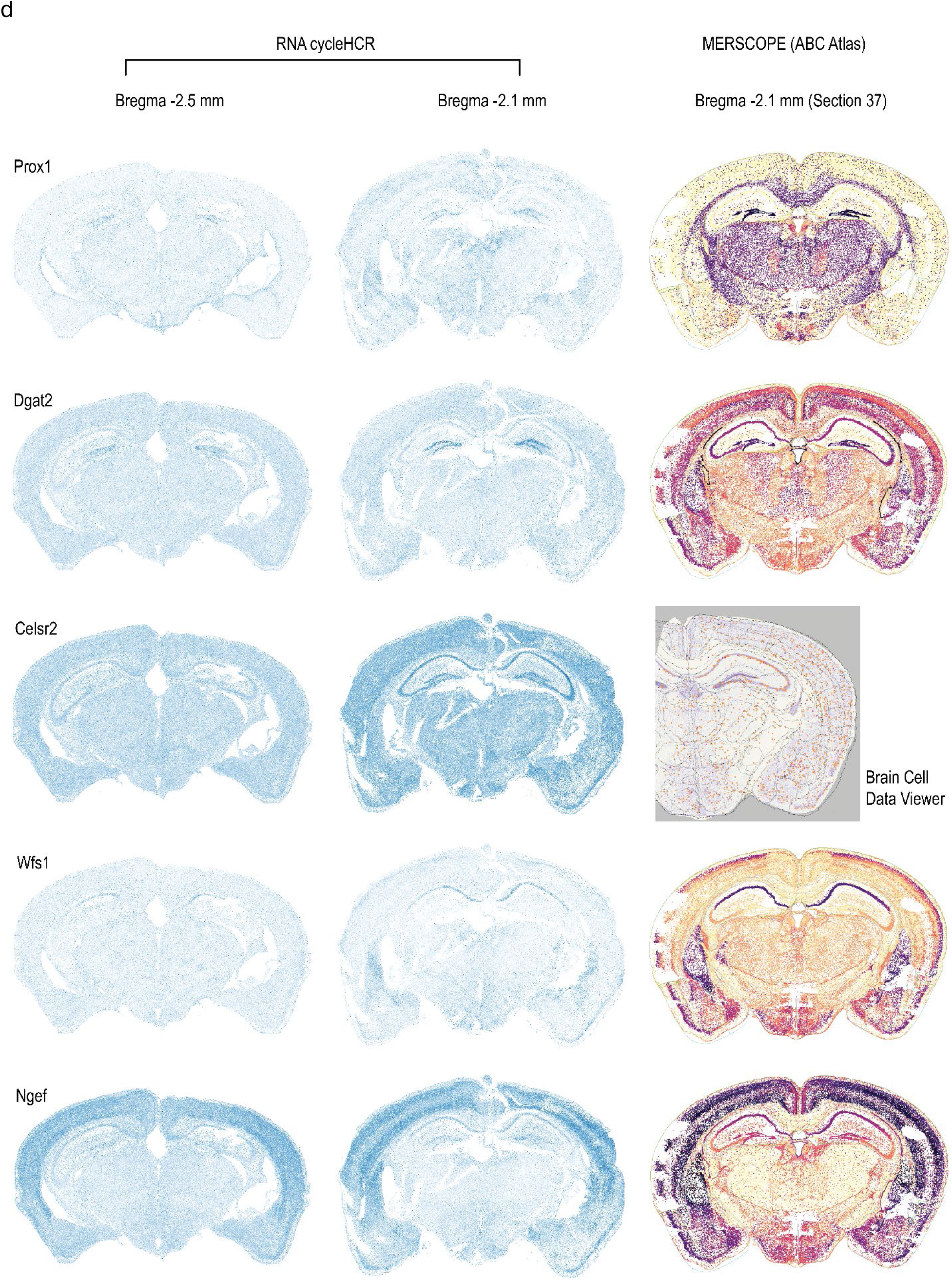
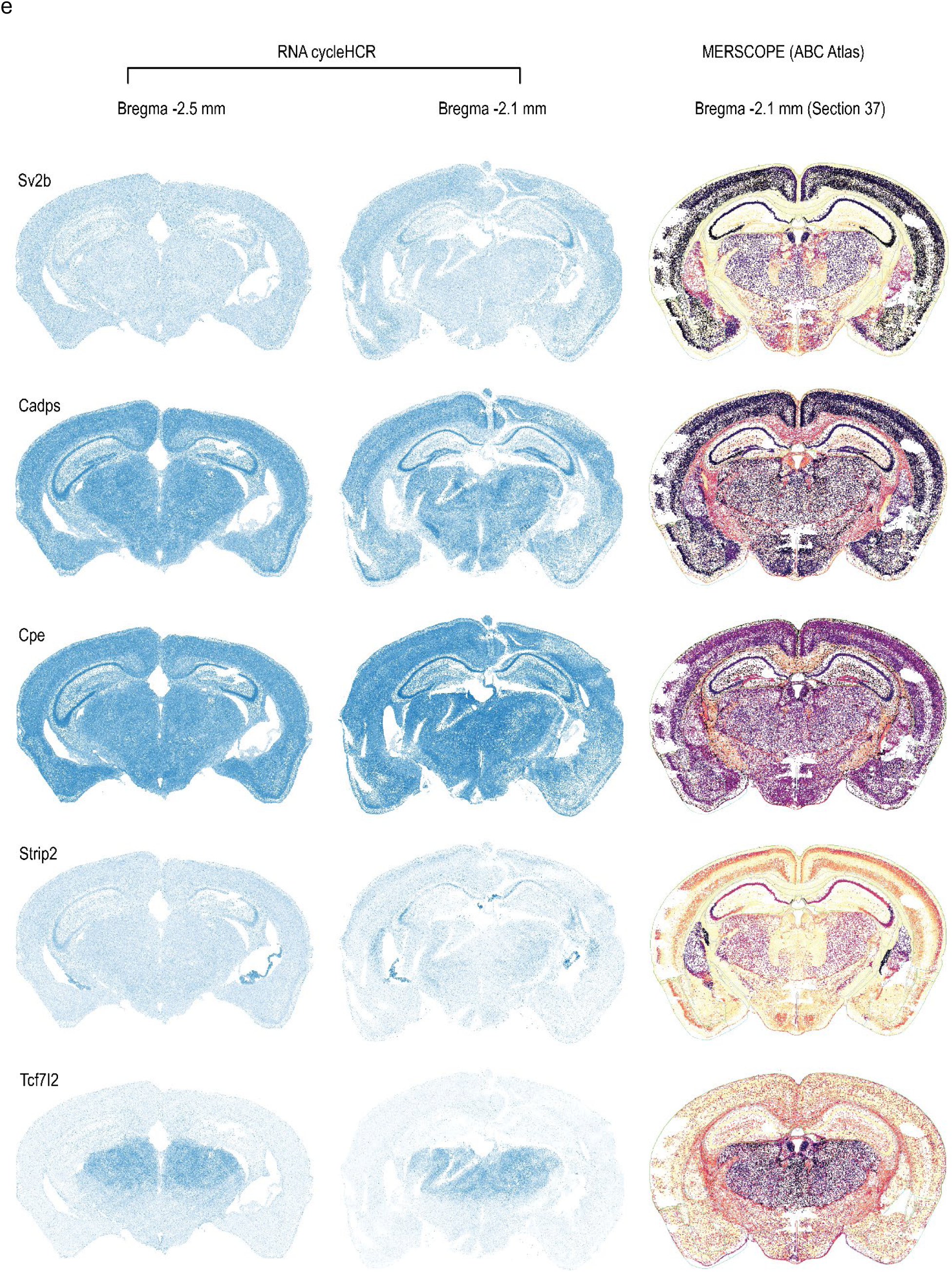
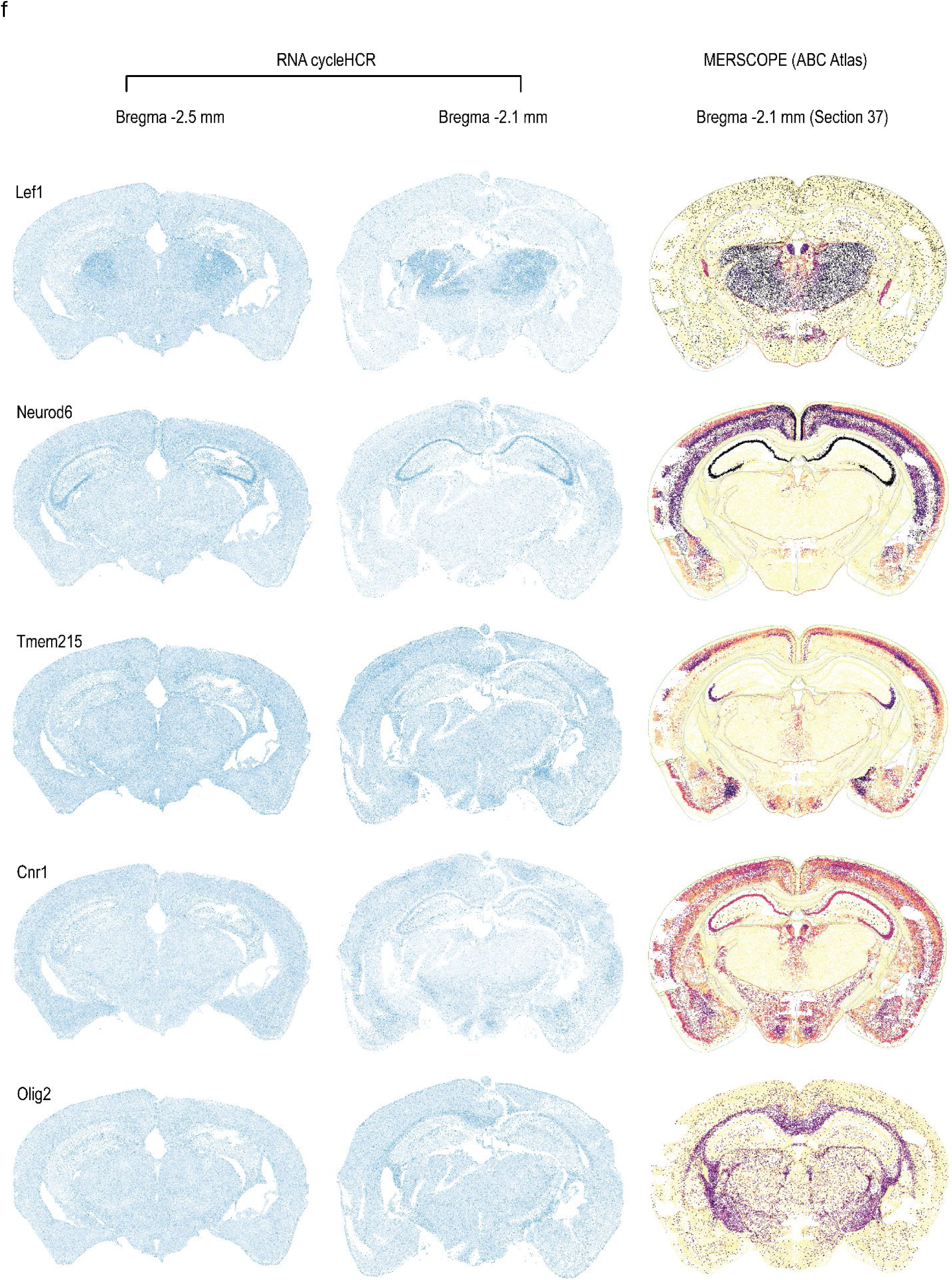
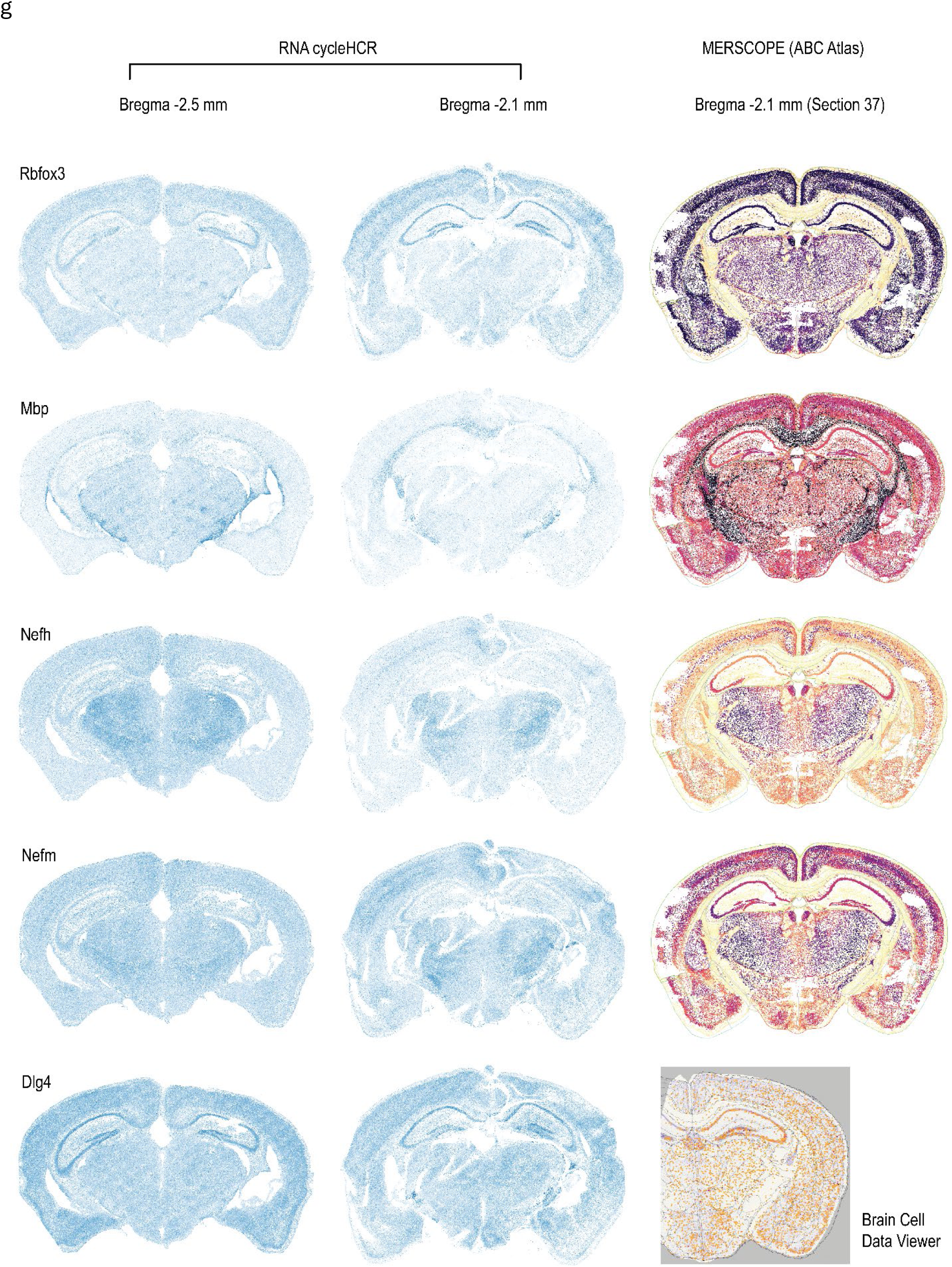
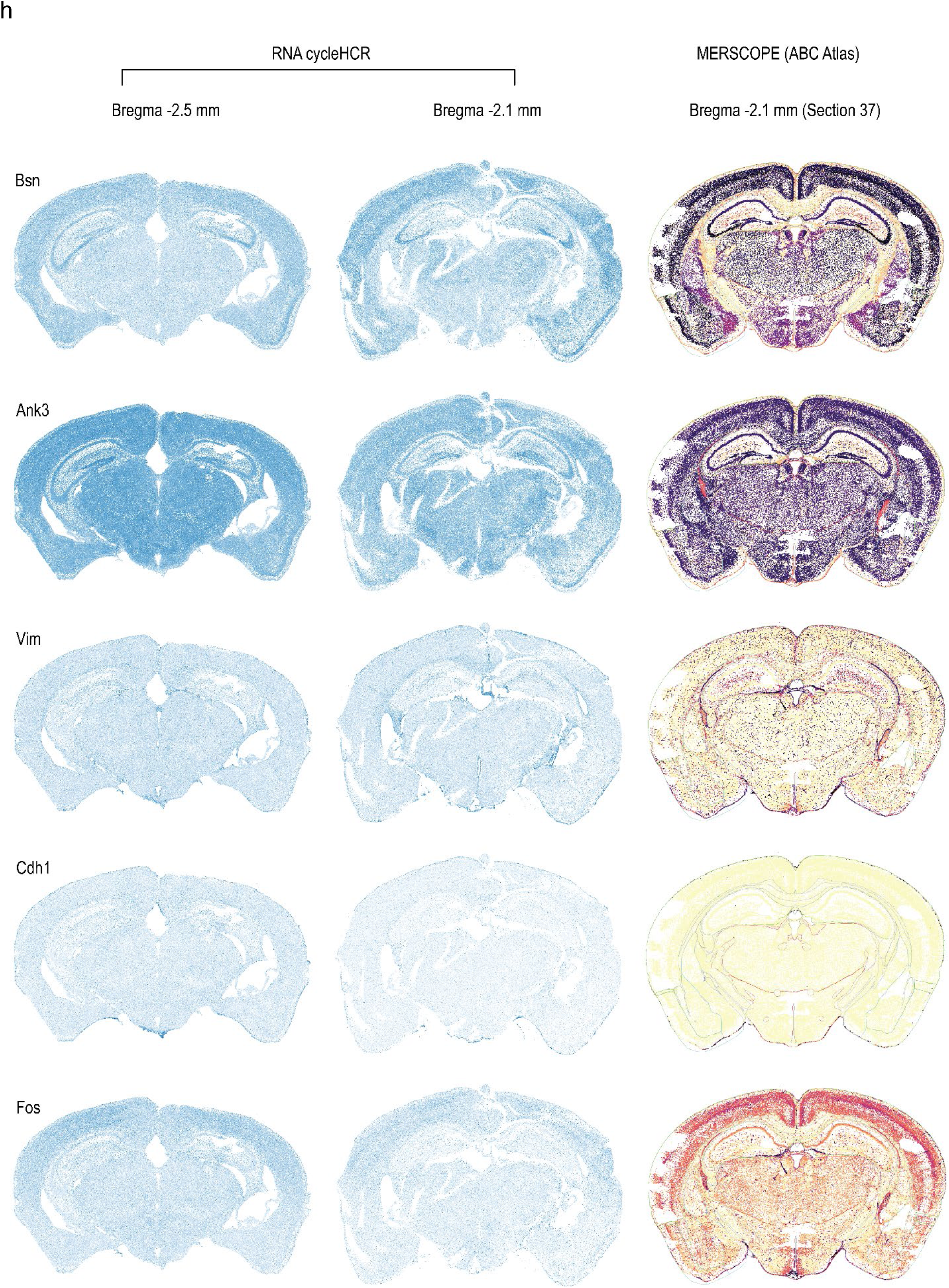
RNA cycleHCR patterns in the two mouse brain sections and cross-method comparison with reference data from the Allen Brain Cell Atlas (ABC Atlas)^20^. **a–h,** The first two columns show the projected patterns of normalized RNA counts/cell of individual genes in the 40-μm mouse whole brain sections. The left column displays Brain Section 1 (Bregma ∼ −2.5 mm); the middle column displays Brain Section 2 (Bregma ∼ −2.1 mm). The right column displays RNA expression patterns acquired by MERSCOPE v.1 in the ABC Atlas. Mouse brain section No. 37 with a similar Bregma coordinate of Brain Section 2 was selected from the ABC Atlas. For genes not included in the ABC Atlas, spatial expression patterns were obtained from Section No. 047 in the Brain Cell Data Viewer^33^ (https://www.braincelldata.org/genex).

**Extended Data Fig. 15.**
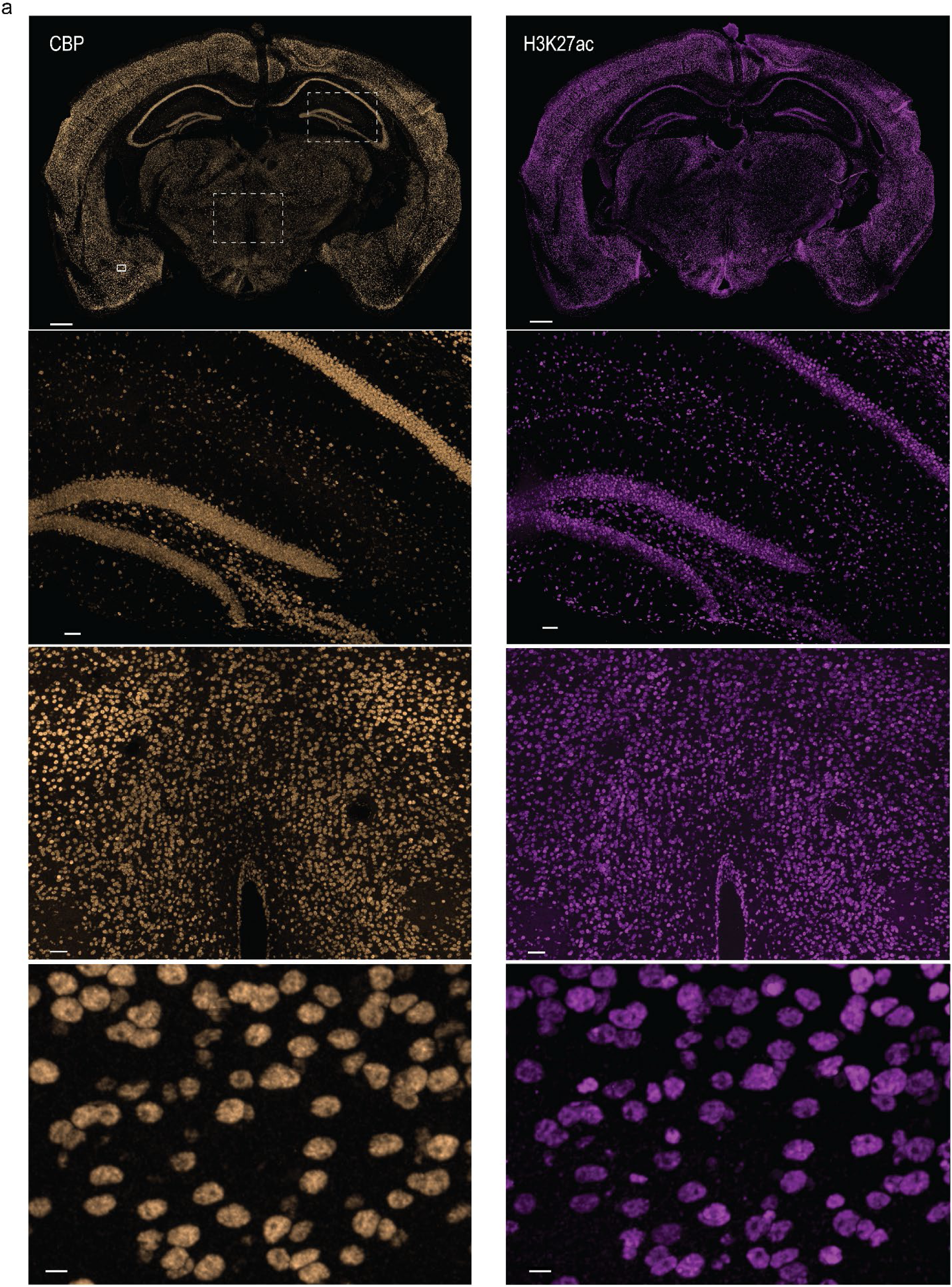
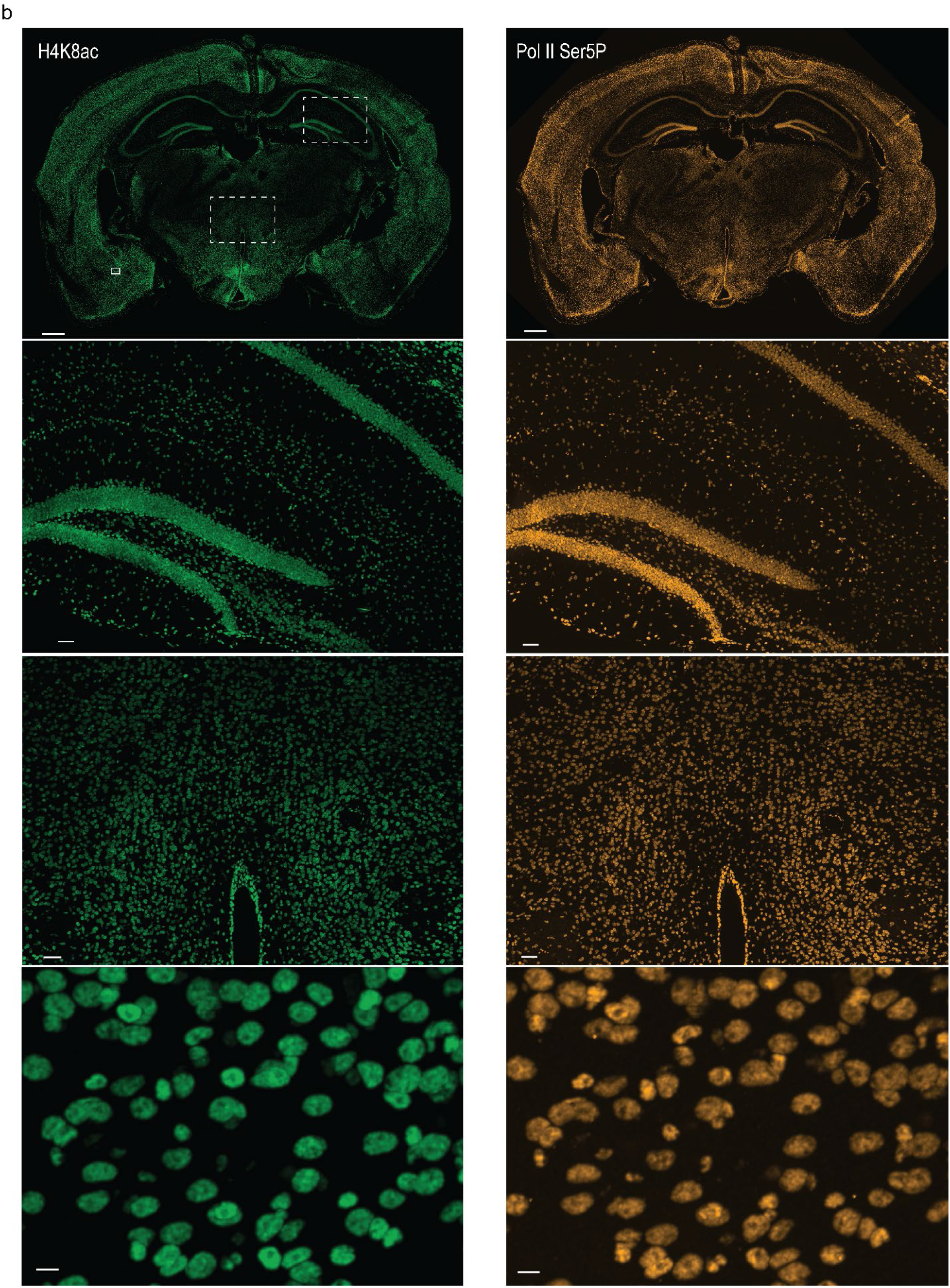
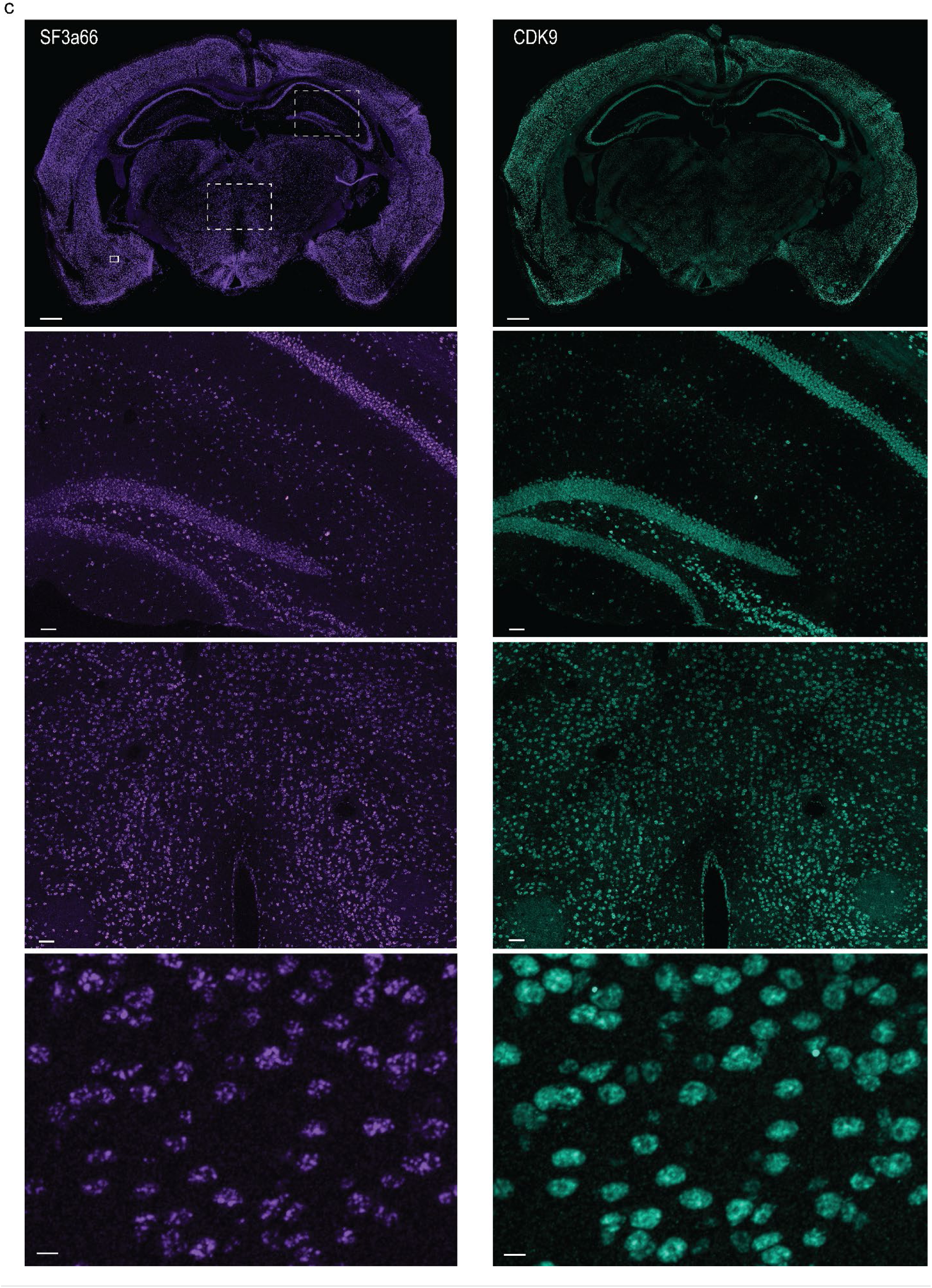
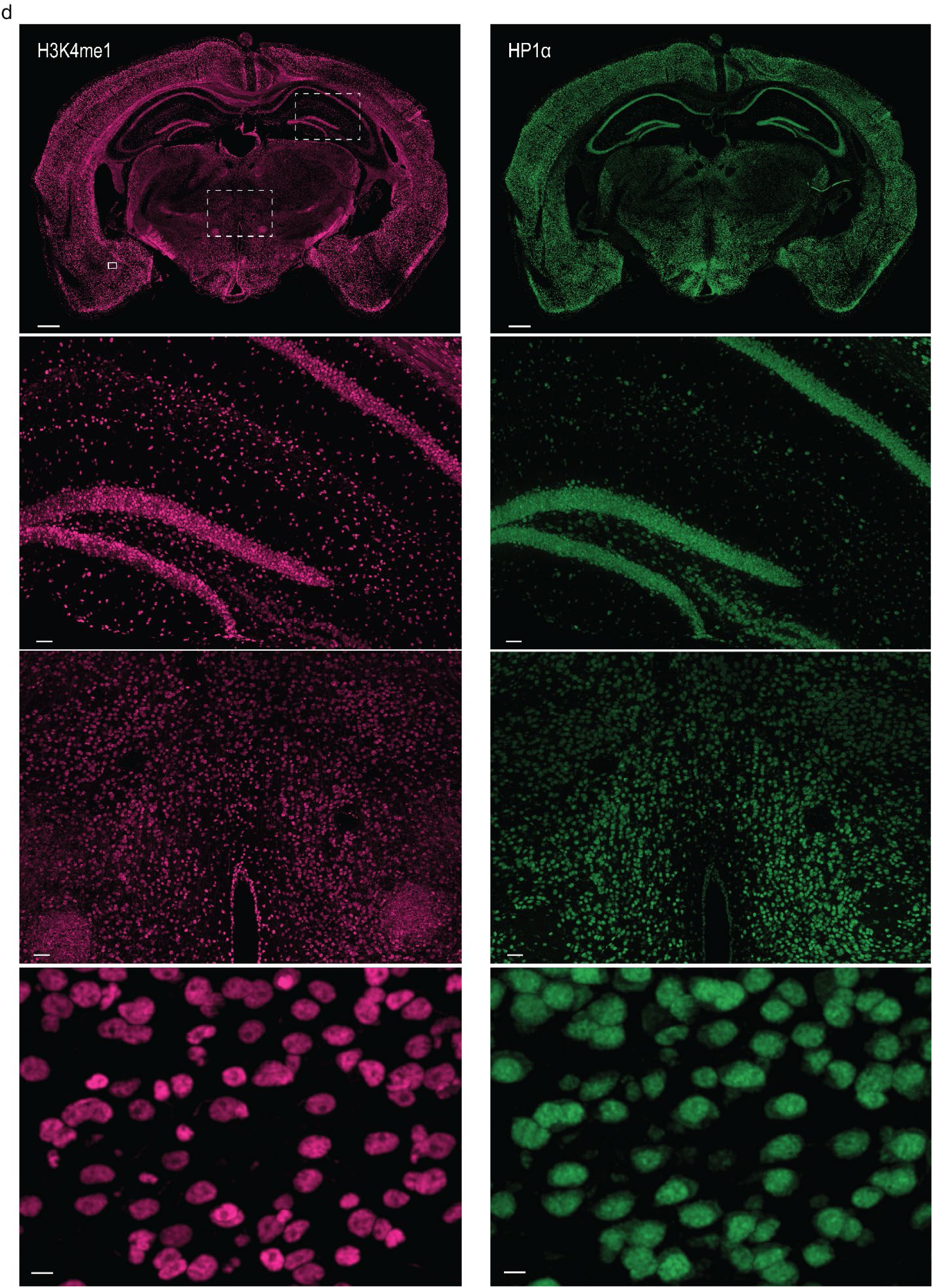
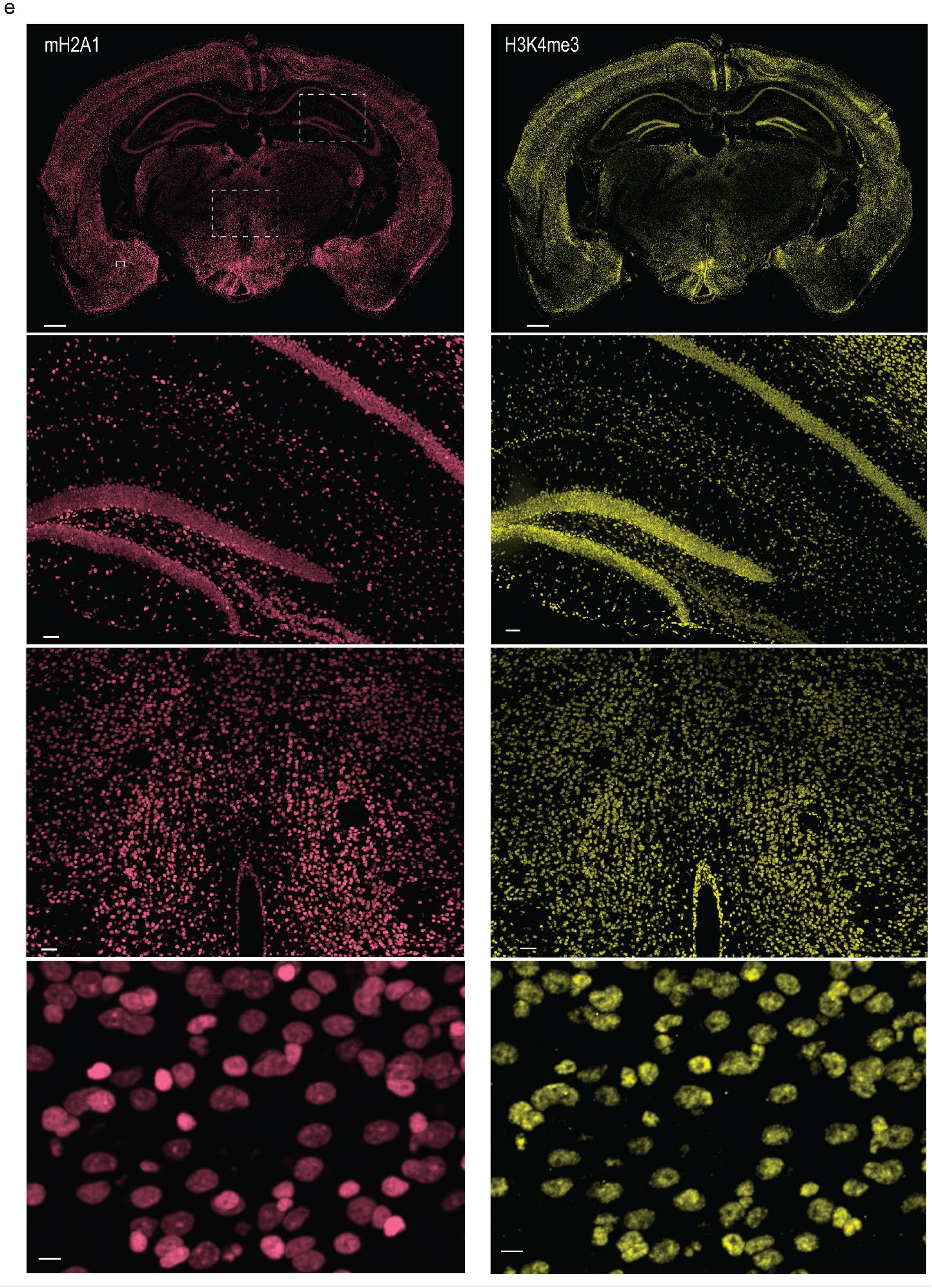
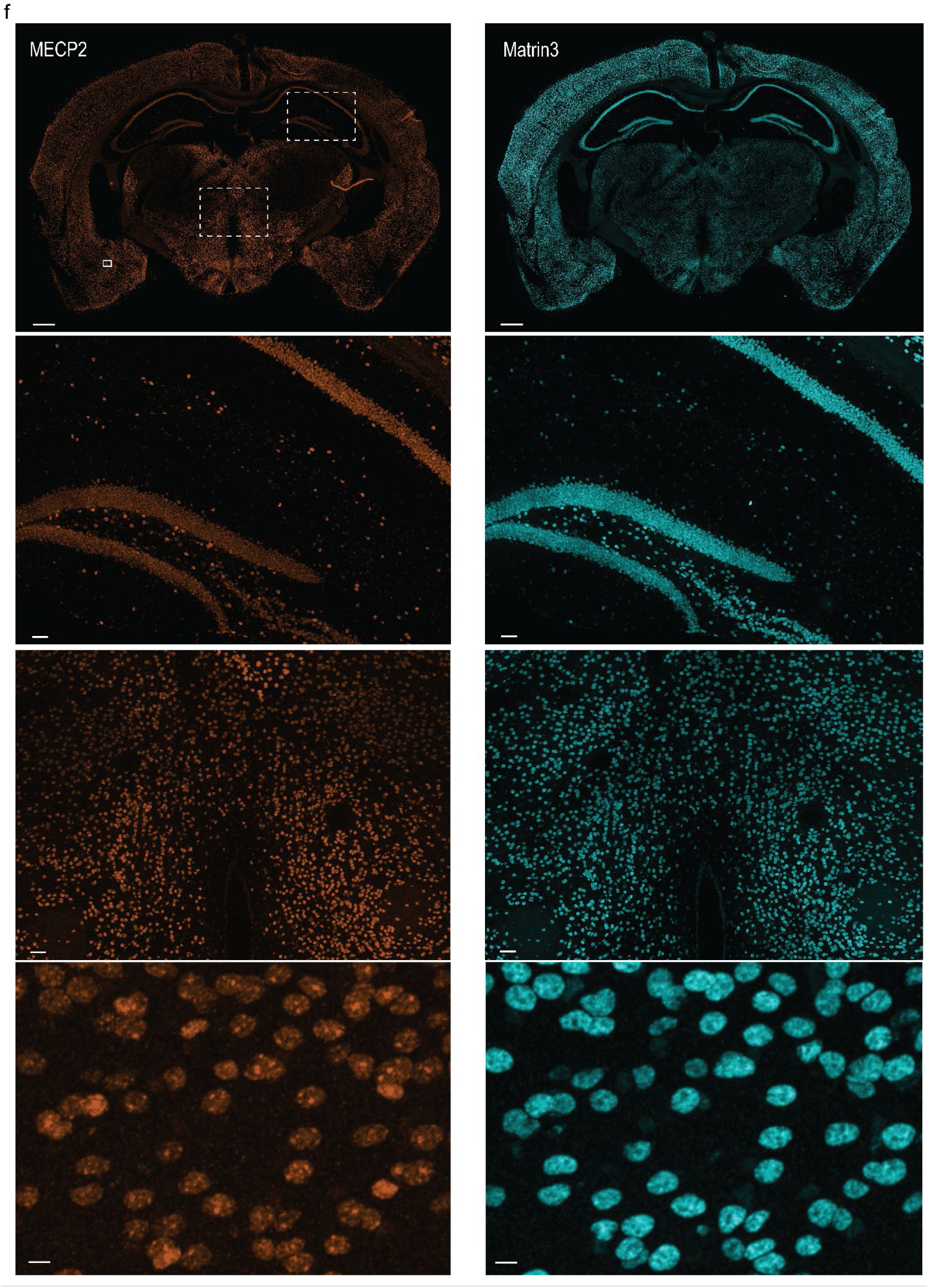
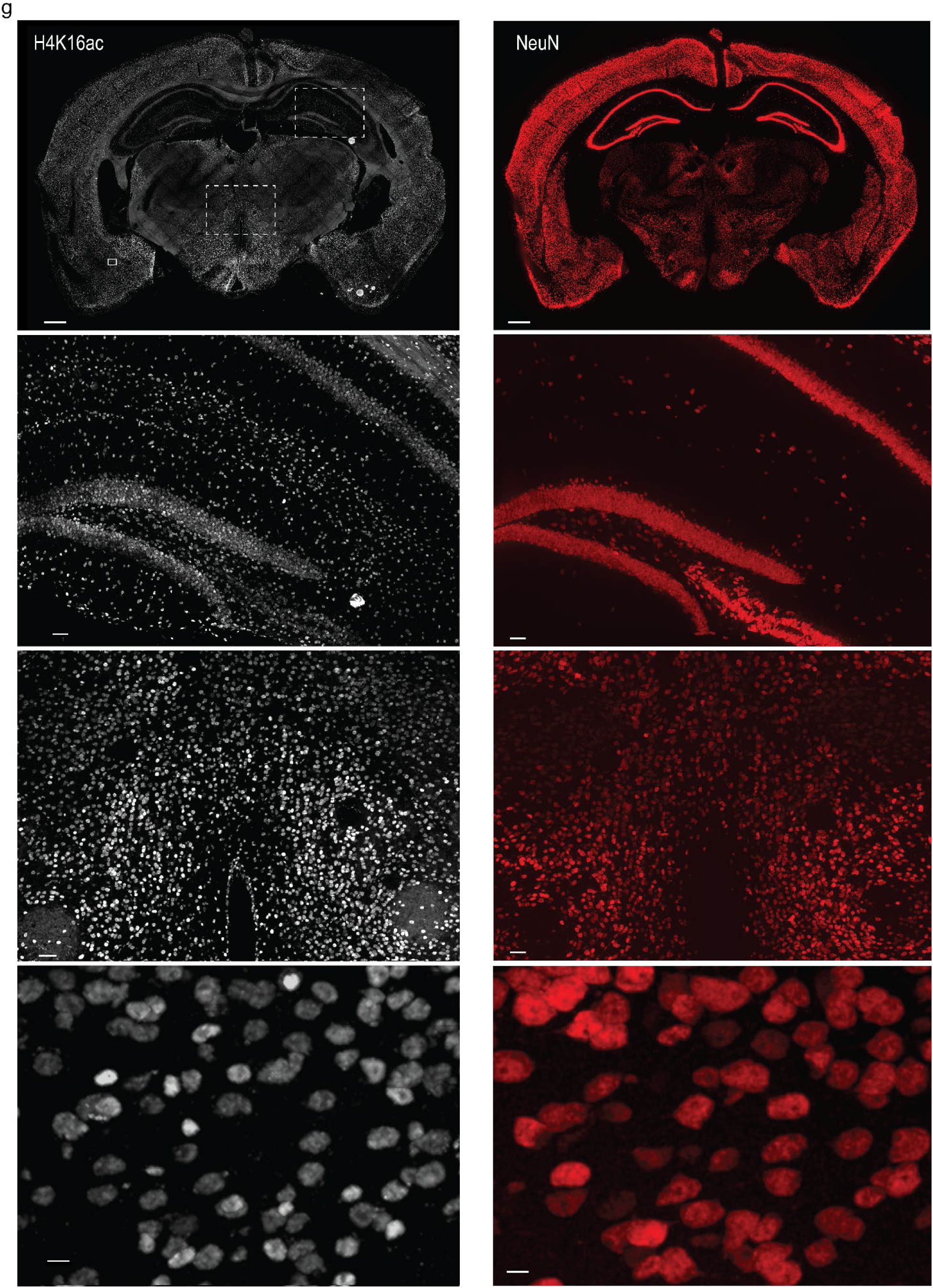
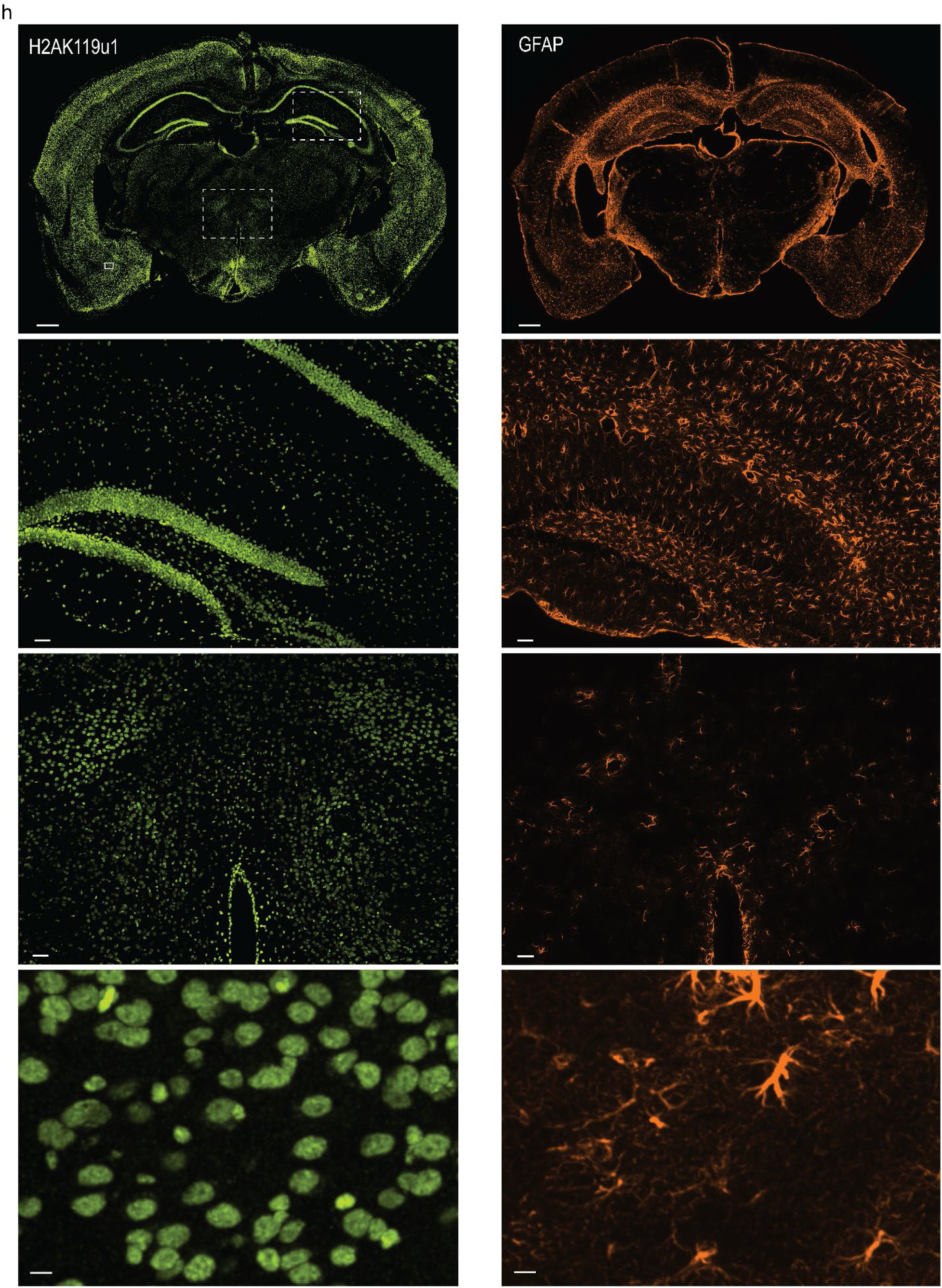
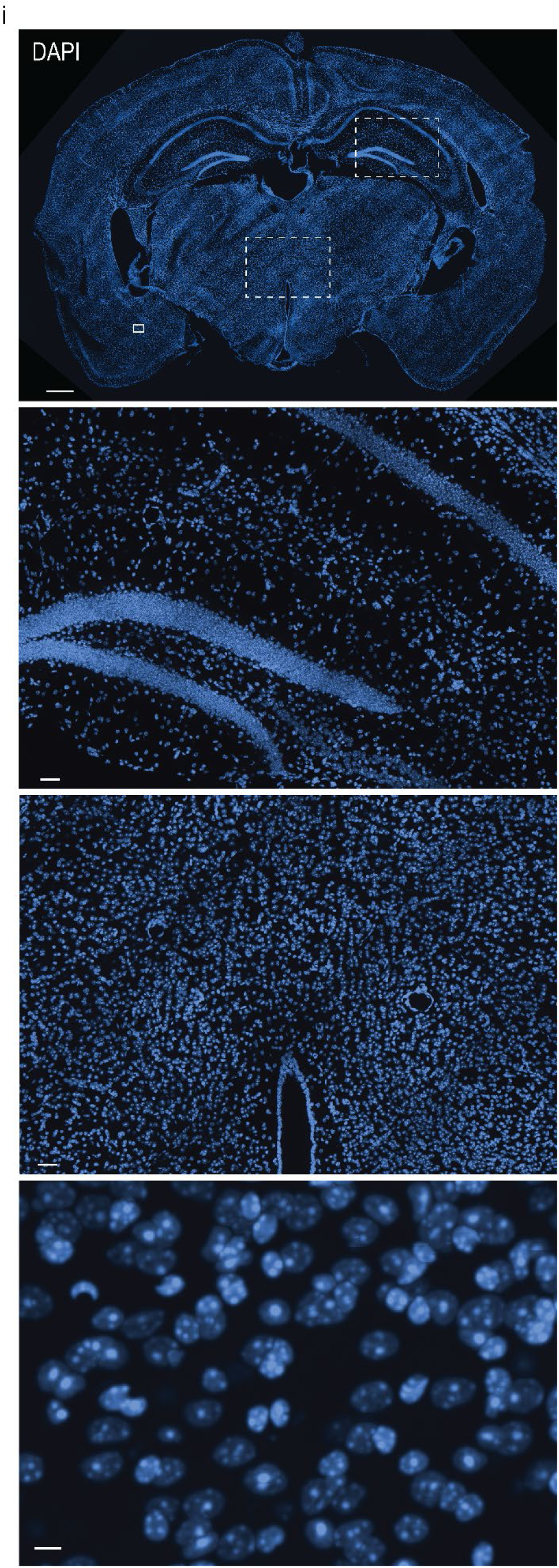
Protein images acquired by multiplex protein and RNA cycleHCR in Brain Section 2. **a–i**, 3D views of stitched and registered images of the 40-μm mouse brain section visualized in Imaris. The top rows display the whole brain section. The 3 rows below are enlarged views of the boxed regions in the top image. Variation of nuclear protein fluorescence intensities are observed across different anatomical regions of the mouse brain. **a,** CBP (CREB Binding Lysine Acetyltransferase. Acetylates histones), H3K27ac, **b,** H4K8ac (active gene transcription), PolII Ser5phos, **c,** SF3a66, CDK9, **d,** H3K4me1, HP1α, **e,** mH2A1 (macroH2A, transcriptional repression), H3K4me3, **f,** MECP2 (Methyl-CpG-binding protein 2), Matrin3, **g,** H4K16ac (active transcription), NeuN (neuron cell type marker), **h,** H2AK119u1 [stabilizing PRC1 (the Polycomb Repressive Complex 1) and PRC2 activities], GFAP (astrocyte cell type marker), **i,** DAPI (dsDNA) Scale bars, 500 μm (top row), 100 μm (second row), 100 μm (third row), 10 μm (bottom row).

**Extended Data Fig. 16.**
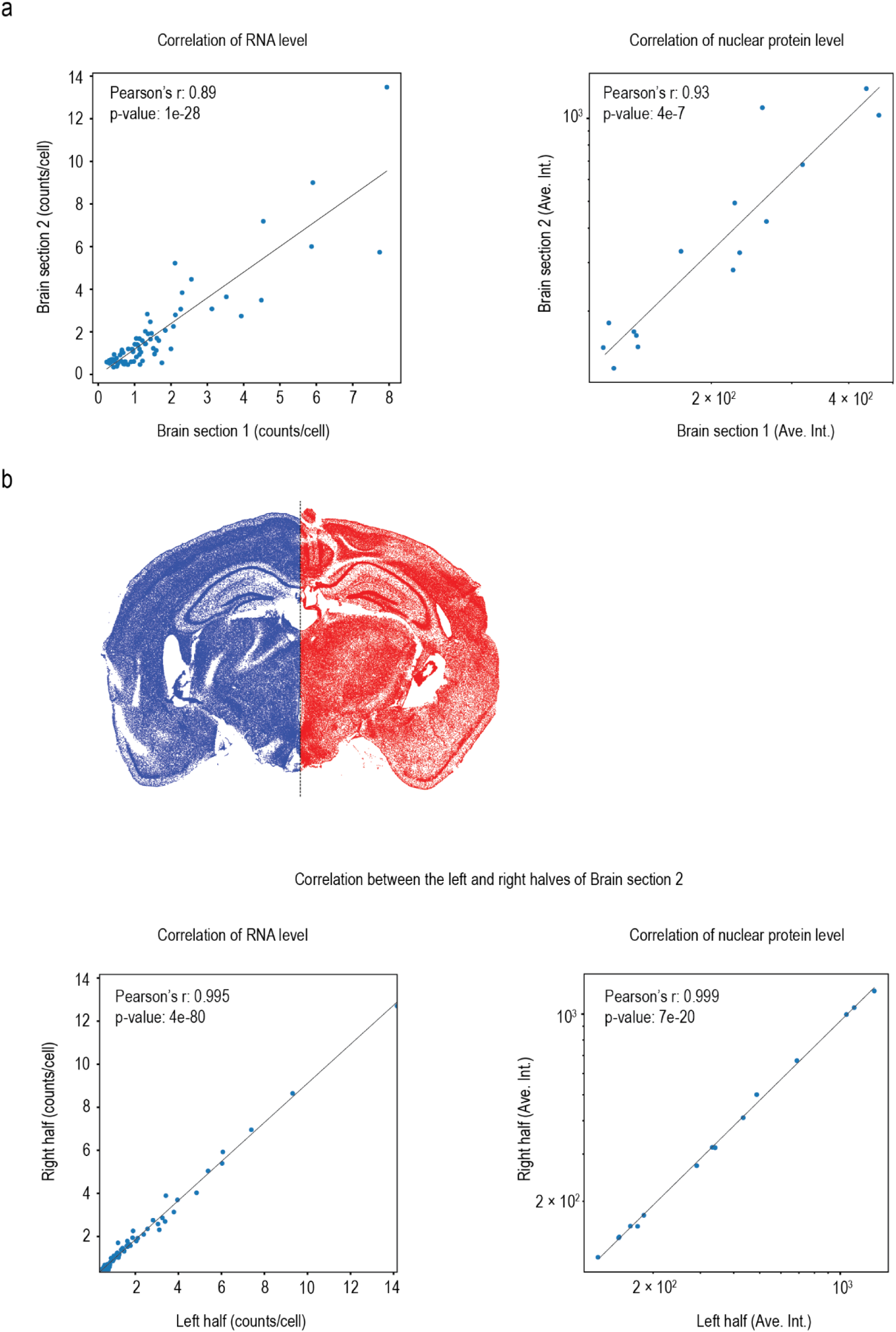
Reproducibility of RNA and protein cycleHCR evaluated by correlations across biological replicates and across spatial coordinates. **a,** Cross-replicate correlations for RNA levels and nuclear protein levels between two mouse brain sections. RNA levels were quantified as average raw spot counts/cell for each gene. Nuclear protein levels were quantified as average fluorescence intensity per nucleus for each protein. **b**, Spatial correlations of RNA levels and nuclear protein levels between left and right halves of the same mouse brain section.

**Extended Data Fig. 17.**
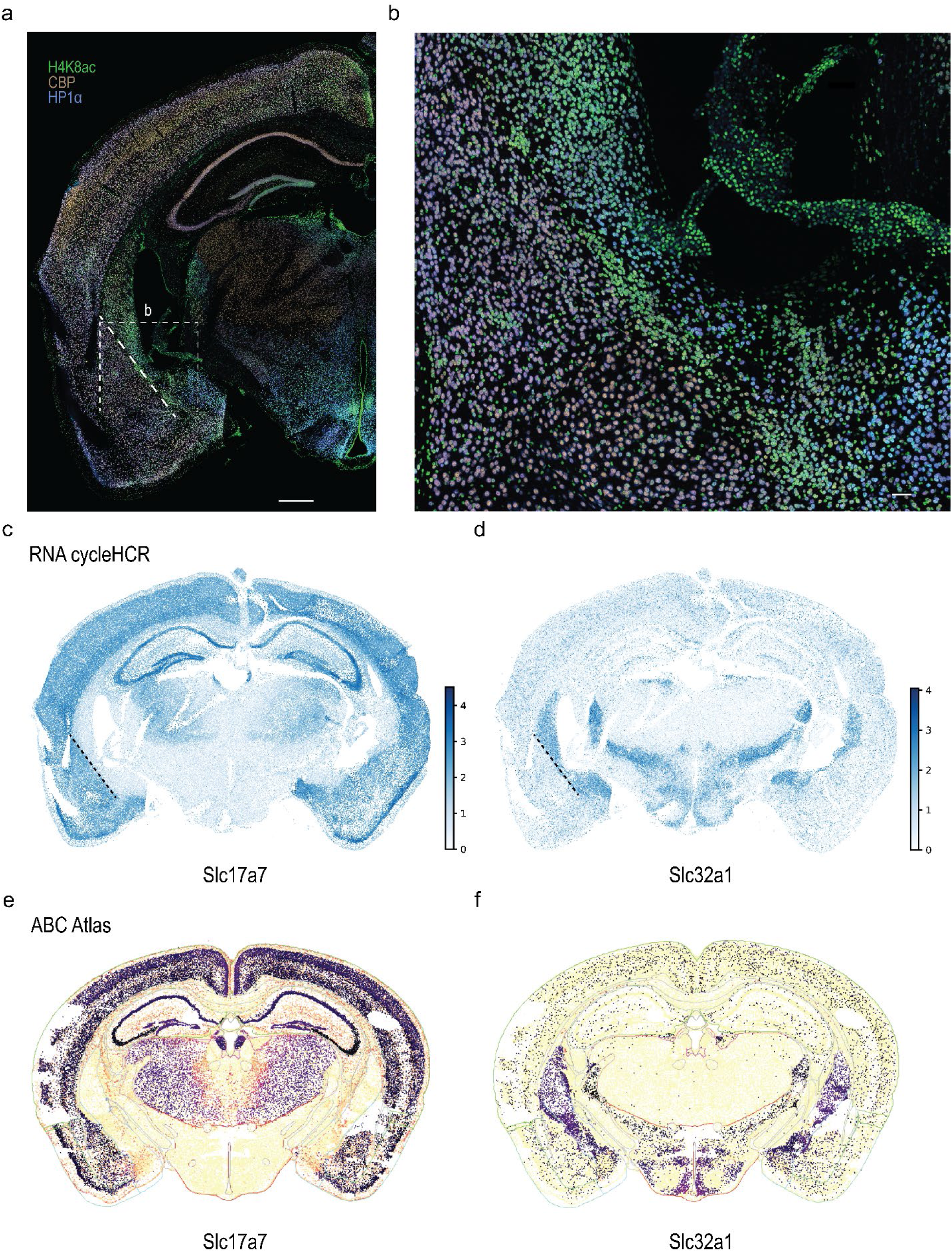
Nuclear protein images exhibit distinct fluorescence intensities in different anatomical regions of the mouse brain. **a,** A composite image of three nuclear proteins (CBP, H4K8ac, and HP1α) exhibits varying fluorescence intensity in different anatomical regions in the mouse brain. The white dashed line highlights the boundary of higher CBP fluorescence intensity on the left (cerebral cortex) and higher H4K8ac fluorescence intensity on the right (cerebral nuclei). Scale bar, 500 μm. **b,** Closer examination of the nuclear protein expression patterns in the boxed region in **a**. Scale bar, 50 μm. **c–d,** The same region with distinct nuclear protein expression patterns highlighted in **a,b** also display distinct RNA expression patterns. RNA cycleHCR patterns of a glutamatergic neuron marker, *Slc17a7* (*VGLUT1*) (**c**), and a GABAergic neuron marker, *Slc32a1* (*VGAT*) (**d**) are shown. The black dashed line highlights the same regional boundary in **a**. **e–f**, RNA expression patterns of *Slc17a7* (**e**) and *Slc32a1* (**f**) from the ABC Atlas show consistent spatial patterns.

**Extended Data Fig. 18.**
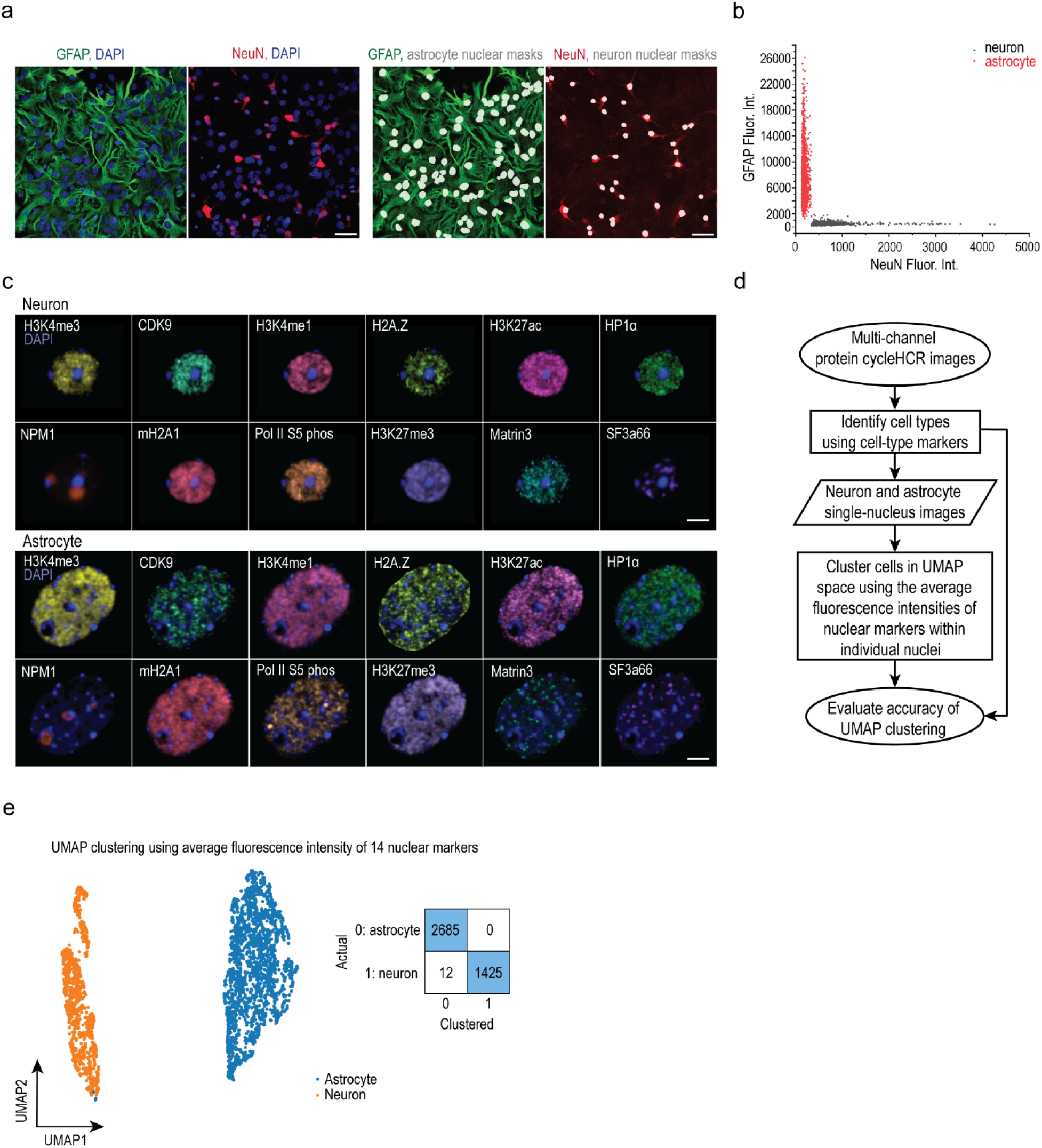
Nuclear protein fluorescence intensities enable accurate classification of neurons and astrocytes in primary cell culture. **a,** Identification of cell types in mouse hippocampal primary cell culture using GFAP (astrocyte marker) and NeuN (neuron marker). The two images on the left show the maximum intensity projection of GFAP and DAPI (first image) and NeuN and DAPI (second image) from the same field of view. The two images on the right display nuclear masks of identified astrocytes overlaid with GFAP image (third image) and nuclear masks of identified neurons overlaid with NeuN image (fourth image). Scale bars, 50 μm. **b,** Average fluorescence intensity of GFAP in a perinuclear region (y-axis) versus average NeuN fluorescence intensity (x-axis) for identified neurons and astrocytes. Perinuclear GFAP intensity was measured in a 5-pixel-wide ring surrounding the nuclear mask at the middle z-plane. NeuN intensity was calculated as the mean value within each 3D nuclear mask. **c**, Twelve nuclear protein markers imaged by cycleHCR, including H3K4me3 (promoter of active genes), CDK9 (gene transcription regulation), H3K4me1 (enhancers), H2A.Z (transcription start sites), H3K27ac (active chromatin), HP1α (heterochromatin), NPM1 (nucleolus granular component), mH2A1 (gene repression and activation), Pol II Ser5P, (transcription initiation), H3K27me3 (heterochromatin), Matrin3 (RNA-binding nuclear matrix), and SF3a66 (spliceosome). These protein images are overlaid with DAPI, shown in 3D rendering for a neuron and an astrocyte using Imaris. Scale bars, 5 um. **d,** Analysis workflow for classifying neurons and astrocytes using nuclear marker images acquired by protein cycleHCR. **e,** Single-cell average fluorescence intensities of 14 nuclear markers (fibrillarin, DAPI in addition to 12 proteins shown in (c)) were used for UMAP clustering to distinguish neurons and astrocytes, achieving 99.7% separation accuracy [(2685 + 1425) / (2685 + 1437)] when compared to NeuN/GFAP-based cell type identification.

**Extended Data Fig. 19.**
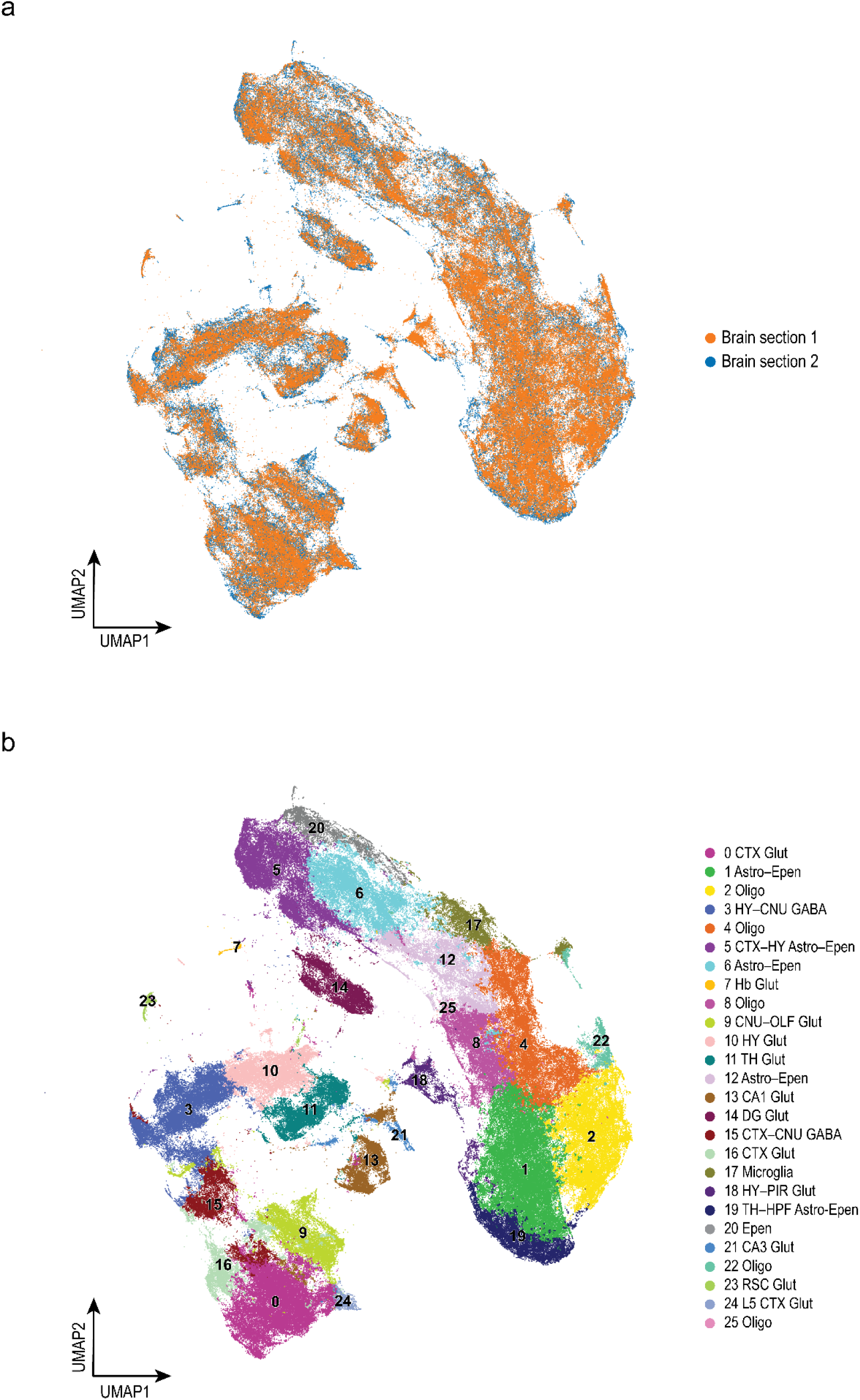
Integrated UMAP using nuclear marker average fluorescence intensities of two mouse brain replicates. **a**, Single-nucleus average fluorescence intensities of 15 nuclear markers were used to generate UMAP. 373,817 cells from the two brain sections were integrated in the same UMAP using Harmony^34^ (178,927 cells from Brain Section 1; 194,890 cells from Brain Section 2). **b**, Clusters of cells from the two brain sections based on the UMAP in **a**.

**Extended Data Fig. 20.**
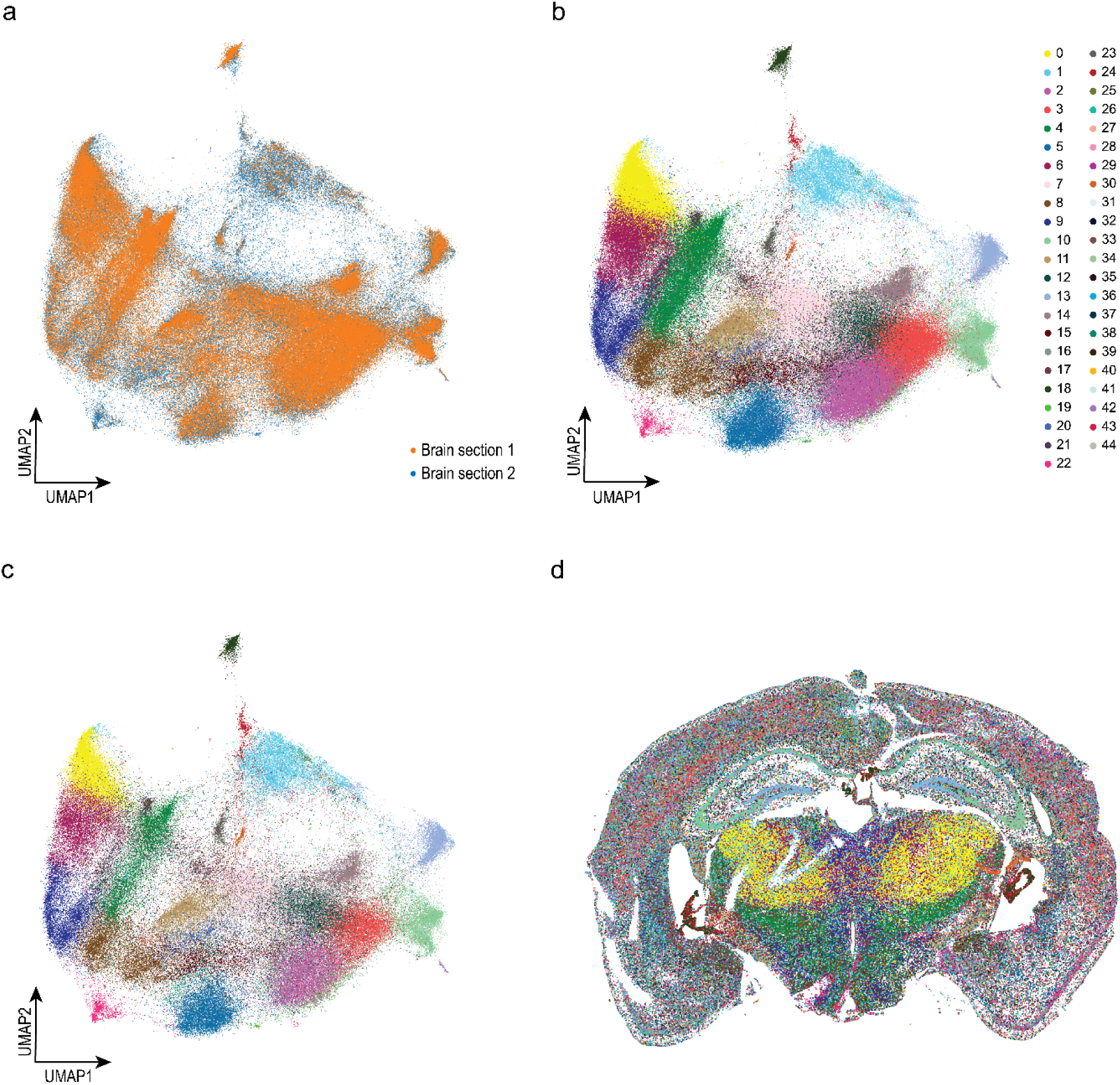
UMAP clustering using single-cell gene expression levels from two mouse brain section replicates imaged by joint protein and RNA cycleHCR. **a**, Single-cell RNA expression levels of 79 genes were used to construct UMAP. Cells from the two brain sections were integrated in the same UMAP space using Harmony. **b**, Clusters of cells from two brain sections based on the UMAP in **a**. **c**, Clusters of cells from Brain Section 2. **d**, Spatial locations of cells from Brain Section 2, colored by RNA UMAP cluster assignment. The same color scheme from **b** is used in **c,d**.

**Extended Data Fig. 21.**
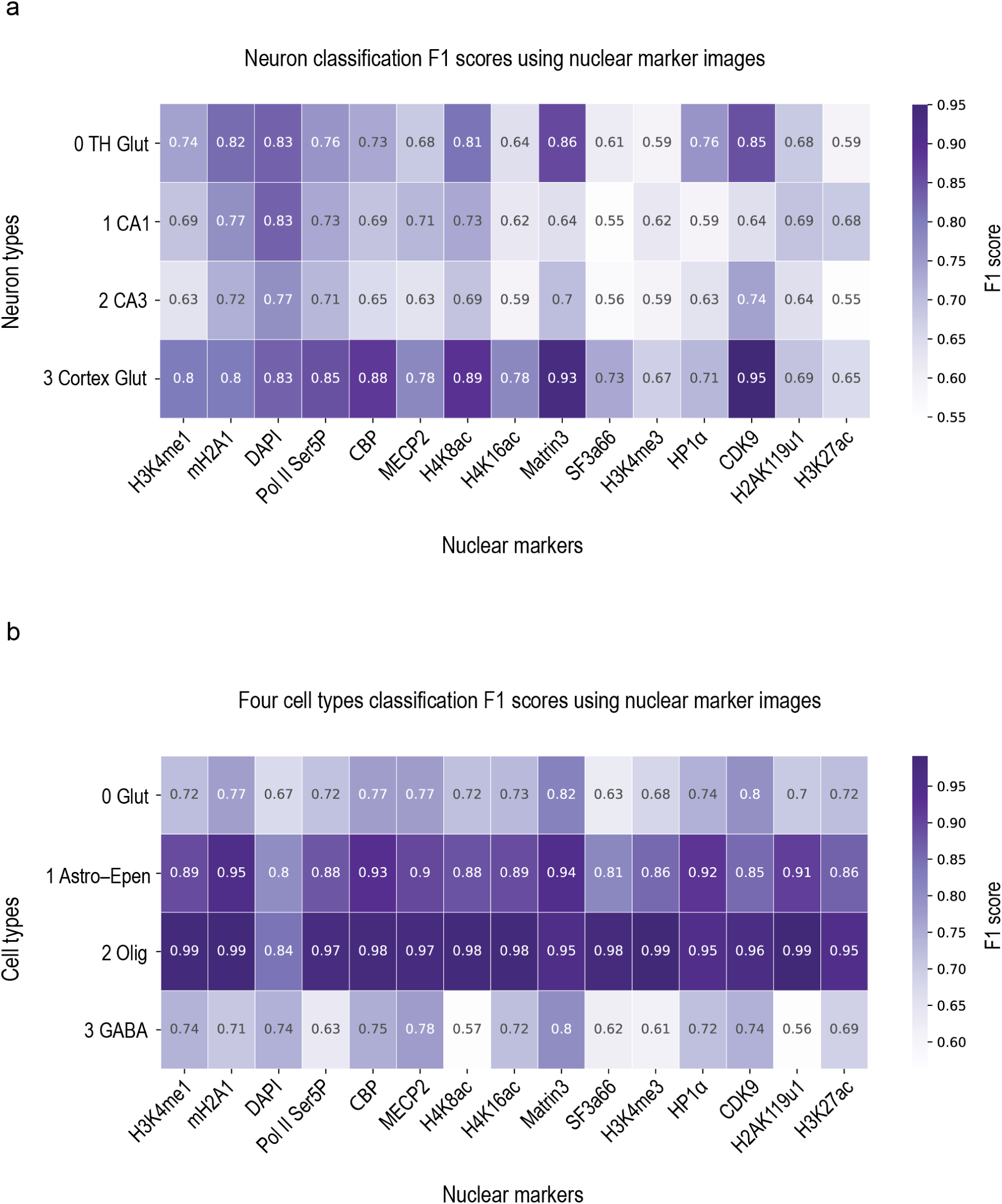
Classification F1 scores using one-channel nuclear marker images. Image classifiers were trained using randomly selected training datasets of ∼1300 single-nucleus images per class with a modified multi-channel microscopy classification workflow from the Medical Open Network for AI (MONAI). Classifiers were validated using validation datasets of ∼600 single-nucleus images per class. **a**, F1 scores for classification of 4 classes of excitatory neurons in different anatomical regions as described in Fig. 6. **b**, F1 scores for classification of 4 classes of different cell types as described in Extended Data Fig. 23.

**Extended Data Fig. 22.**
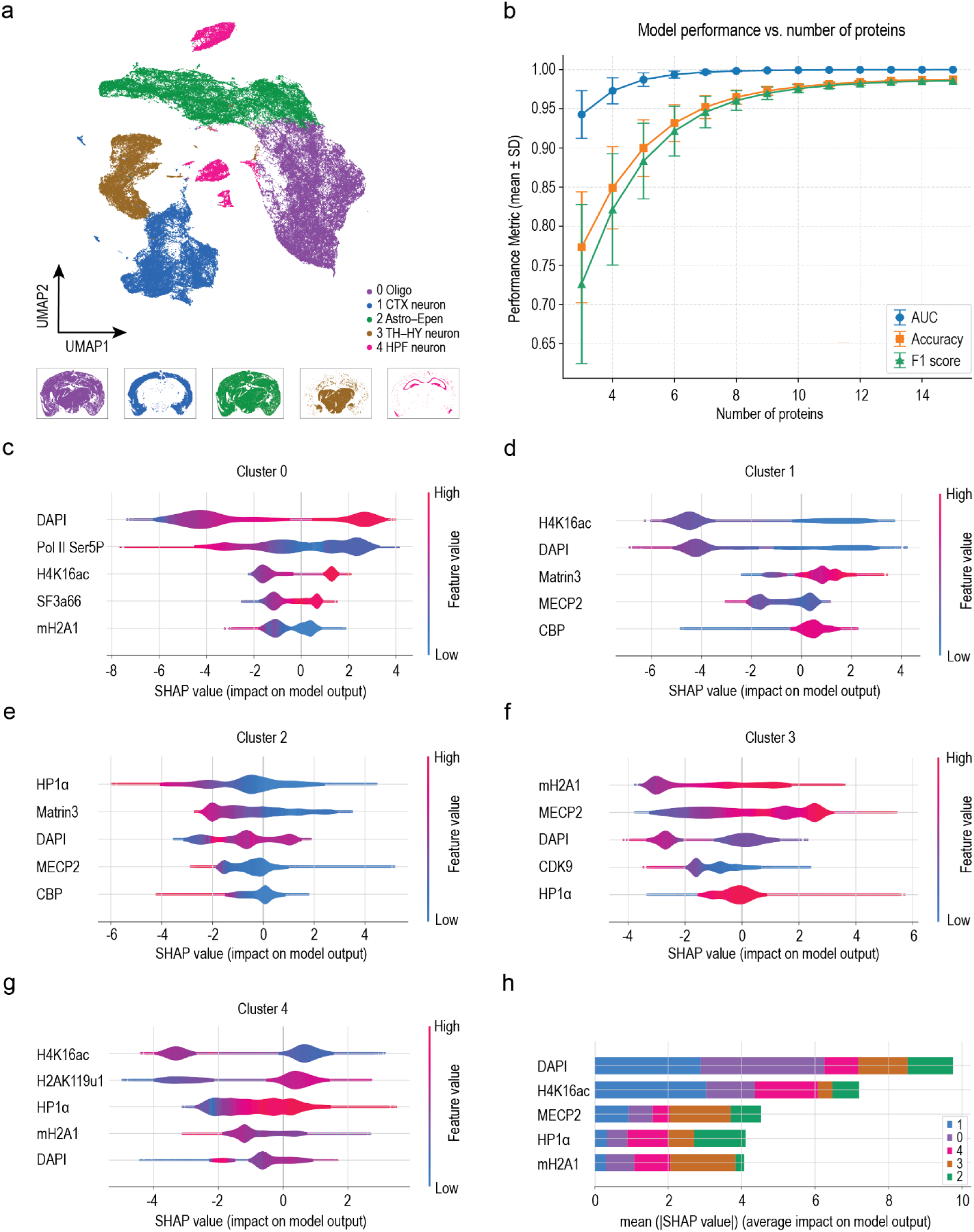
Classification performance using varying numbers of nuclear protein intensities and importance of different proteins to classification. **a**, Five major cell type clusters comprising all 194,724 cells in Brain Section 2, constructed using nuclear protein intensities. Cell types were determined based on highly expressed marker genes in each cluster (Cluster 0: Sox10; Cluster 1: Ngef, Slc17a7; Cluster 2: Slc1a3; Cluster 3: Slc17a6; Cluster 4: Slc17a7). **b**, Average XGBoost^35^ classification performance metrics (accuracy, F1 score, and AUC) using nuclear protein intensities as a function of the number of proteins used. Each curve represents the average (± SD) across all cell type classifications using different combinations of proteins. **c–g**, The XGBoost classification model trained with 15 nuclear protein intensities was used for SHAP (SHapley Additive exPlanations)^24^ analysis. SHAP value distribution plots showing the top 5 nuclear marker features with the highest impact on classification decisions for each of the five cell clusters. Color indicates high (red) or low (blue) nuclear protein intensity values. Positive SHAP values indicate positive predictions for the cluster, whereas negative values indicate negative predictions. **h**, Overall importance of each nuclear protein’s intensity on classification predictions across all cell clusters, calculated as mean absolute SHAP values for each cluster. Stacked bars show cluster-specific contributions. Only the top 5 nuclear markers are shown.

**Extended Data Fig. 23.**
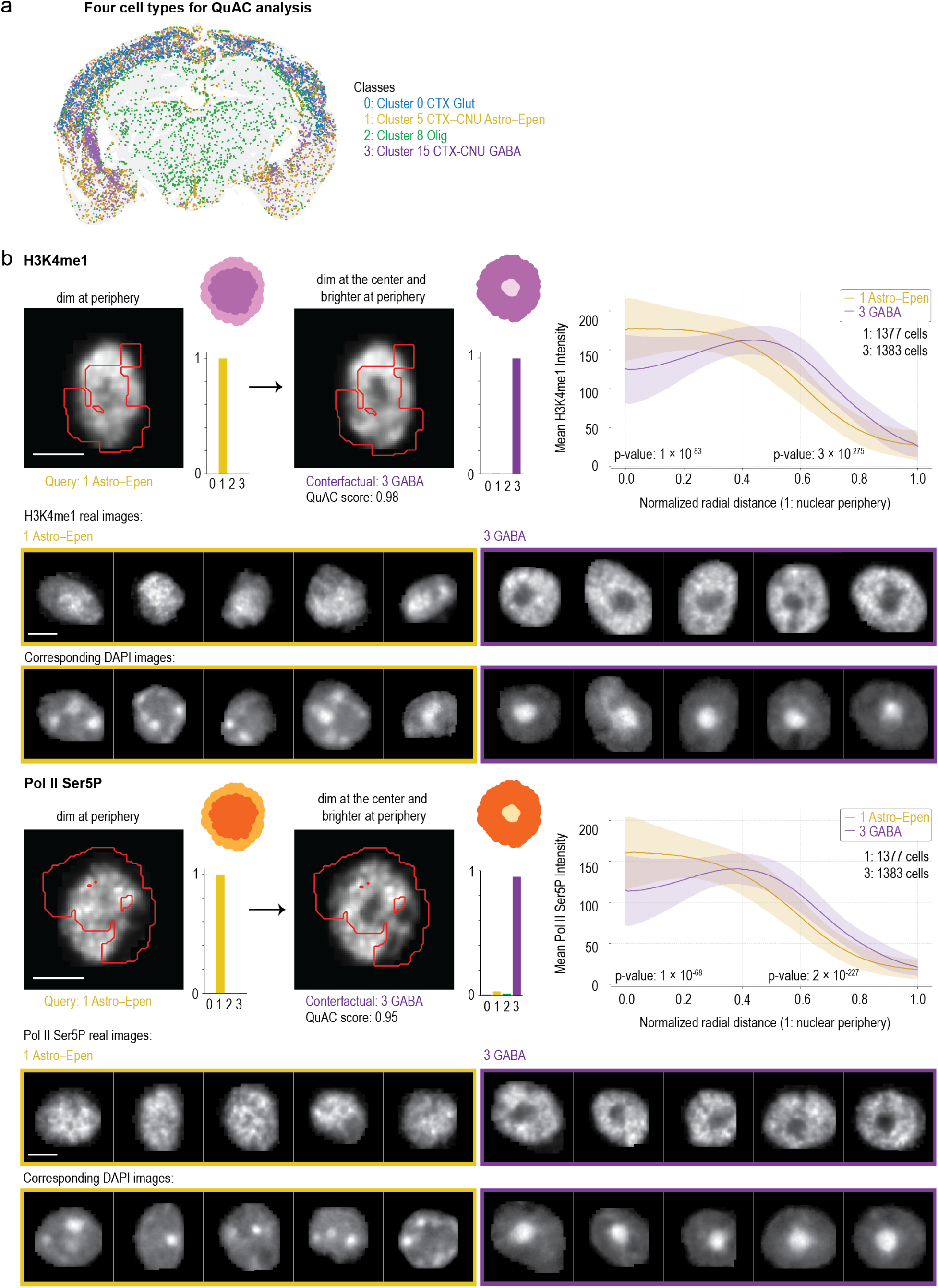
Cell-type-specific nuclear architectures revealed by explainable ML. **a,** Four cell clusters of different cell types with overlapping spatial distributions in the cerebral cortex, including glutamatergic neurons, astrocyte–ependymal cells, oligodendrocytes, and GABAergic neurons, were selected from UMAP clustering (∼2,000 randomly sampled cells per cluster) for QuAC analysis as described in Fig. 6a. **b**, Distinct euchromatin distribution features in two cell types discovered by QuAC. Representative query and counterfactual images of euchromatin markers H3K4me1 and Pol II Ser5P demonstrate QuAC-identified cell-type-specific nuclear features. Class 1 Astro–Epen cells exhibit predominant localization of active euchromatin near the nuclear center and heterochromatin near the nuclear periphery. Class 3 GABAergic neurons display a common pattern of heterochromatin at the nuclear center with euchromatin distributed throughout other nuclear regions. Population-level measurements of average intensity versus radial distance from the nuclear center for these two nuclear markers reveal significant differences in their radial distributions. P-values for average nuclear protein intensity were calculated at the nuclear center (radial distance = 0) and near the nuclear periphery (radial distance = 0.7). These distribution patterns are also displayed in representative nuclear images alongside their DAPI (heterochromatin) images. P values are reported using pairwise Mann-Whitney U tests. Scale bars, 5 μm.

## Supplementary Table and Table Legends 1 - 3

**Supplementary Table 1.** List of antibodies used for protein cycleHCR. Forty-six antibodies imaged through joint RNA and protein cycleHCR together with 79 genes in the same mouse brain section. Three additional antibodies were only for imaging 7 proteins in a hippocampal section using the same left barcode and highly similar right barcodes. ^a^ Different antibodies were conjugated with the same oYo-Link oligo sequence and hybridized with different L + R barcode pairs in multiplex protein cycleHCR. Among the 46 antibody complexes used simultaneously, 11 antibodies were conjugated with oYo-Link oligo sequence No. 2, and 4 antibodies were conjugated with oYo-Link oligo sequence No. 1. This approach enables multiple antibodies to share the same oYo-Link oligo reagent, significantly reducing the cost of multiplex protein cycleHCR. ^b^ Sample images of protein expression patterns can be found for most targets on antibody vendor websites. Selected publications are listed here as additional references for immunostaining patterns.

|  | Antigen | Host species | Clonality | Vendor | Cat. # | Shared oYo No. <sup>a</sup> | Ref<sup>b</sup> |
| --- | --- | --- | --- | --- | --- | --- | --- |
| **46 antibodies used for joint protein and RNA cycleHCR** |  |  |  |  |  |  |  |
| 1 | H3K27me3 | Rabbit | M | Cell Signaling Technology | 9733 |  |  |
| 2 | H3K27ac | Rabbit | M | Abcam | ab177178 |  |  |
| 3 | H4K8ac | Mouse IgG1 | M | Active Motif | 61525 | 1 |  |
| 4 | Pol II Ser5 phos | Mouse | M | Active Motif | 91119 |  |  |
| 5 | SF3A66 | Mouse IgG1 | M | Abcam | ab77800 | 1 |  |
| 6 | CDK9 | Rabbit | M | Abcam | ab239364 |  |  |
| 7 | CBP | Rabbit | M | Cell Signaling Technology | 7389 | 2 |  |
| 8 | HP1 alpha | Rabbit | M | Abcam | ab109028 | 2 |  |
| 9 | mH2A1 | Rabbit | M | Abcam | ab232602 |  |  |
| 10 | H3K4me3 | Rabbit | P | Abcam | ab8580 |  |  |
| 11 | MECP2 | Rabbit | M | Cell Signaling Technology | 3456 | 2 |  |
| 12 | Matrin 3 | Rabbit | M | Abcam | ab151714 |  |  |
| 13 | H3K4me1 | Rabbit | P | Abcam | ab8895 |  |  |
| 14 | Pol II Ser5 phos | Rabbit | P | Abcam | ab5131 | 2 |  |
| 15 | H4K16ac | Rabbit | P | Sigma-Aldrich | 07-329 | 2 |  |
| 16 | AQP4 | Mouse IgG1 | M | CDI LABS | BCN456.2.1B5 | 1 | 36 |
| 17 | Neuron beta III Tubulin (Tuj1) | Mouse | M | R&D Systems | MAB1195 | 2 |  |
| 18 | H2AK119u1 | Rabbit | M | Cell Signaling Technology | 8240 | 2 |  |
| 19 | NeuN | Rabbit | M | Cell Signaling Technology | 36662 |  | 37-39 |
| 20 | H2A.Z | Rabbit | M | Abcam | ab208691 |  |  |
| 21 | S100B | Rabbit | P | Thermo Fisher | PA5-78161 |  |  |
| 22 | nNOS | Rabbit | M | Abcam | ab76067 |  | 40 |
| 23 | Somatostatin receptor 2 (Sstr2) | Rabbit | M | Abcam | ab134152 | 2 | 41 |
| 24 | Calbindin | Rabbit | M | Abcam | ab108404 | 2 | 42, 43 |
| 25 | Cannabinoid Receptor 1 (CB1) | Rabbit | M | Cell Signaling Technology | 93815 |  | 44 |
| 26 | Parvalbumin (PV) | Mouse | M | CDI LABS | BCN14.1.1 C9 | 2 | 38, 43 |
| 27 | CD31 | Goat | P | R&D Systems | AF3628 |  | 41-43 |
| 28 | ChAT | Goat | P | Millipore | AB144P |  | 45 |
| 29 | Neuropeptide Y (NPY) | Rabbit | M | Cell Signaling Technology | 77100 |  | 46 |
| 30 | Flotillin 1 | Rabbit | P | Abcam | ab133497 |  | 42 |
| 31 | GABA | Rabbit | P | Sigma-Aldrich | A2052 |  | 47 |
| 32 | Vimentin | Rabbit | M | Cell Signaling Technology | 46173 |  | 48 |
| 33 | Calbindin | Rabbit | M | Cell Signaling Technology | 46343 |  |  |
| 34 | Iba-1 | Rabbit | P | FUJIFILM Wako | 019-19741 |  | 42, 43 |
| 35 | PMP70 | Rabbit | P | Invitrogen | PA1-650 |  | 38, 39 |
| 36 | Homer1 | Rabbit | P | SYSY | 160 003 |  | 49 |
| 37 | MAP2 | Rabbit | M | Cell Signaling Technology | 65911 |  | 50 |
| 38 | VGLUT2 | Mouse | M | Abcam | ab79157 | 1 |  |
| 39 | Neuron specific<br>beta III Tubulin | Rabbit | P | Abcam | ab229590 | 2 |  |
| 40 | MBP | Mouse | M | BioLegend | 808401 |  |  |
| 41 | Calnexin | Rabbit | P | Abcam | ab22595 |  | 51 |
| 42 | Tyrosine<br>Hydroxylase (TH) | Mouse | M | BioLegend | 818001 |  | 52 |
| 43 | Parvalbumin (PV) | Rabbit | P | Thermo Fisher | PA1-933 |  | 53 |
| 44 | Na/K ATPase | Rabbit | M | Abcam | ab76020 |  | 40, 41,<br>43 |
| 45 | GFAP | Rabbit | P | Abcam | ab278054 |  | 54 |
| 46 | PSD-95 | Mouse | M | Antibodies Inc | 75-028 |  | 38, 39 |
| **Only in similar barcode labeling of 7 antibodies** |  |  |  |  |  |  |  |
| 47 | Fibrillarin | Rabbit | P | Abcam | ab5821 |  |  |
| 48 | Nup98 | Rabbit | M | Cell Signaling Technology | 2598 |  |  |
| 49 | Ankyrin G | Mouse | M | Antibodies Inc | 75-146 |  |  |

**Supplementary Table 2.** List of 79 genes imaged by joint protein and RNA cycleHCR in mouse brain sections and the associated RNA probe library. Provided in a separate file SpplementaryTableS2.xlsx. Two tables are in this file. The first sub-table details each gene’s ID, fluorescence channel, and L + R barcodes. The second sub-table contains the sequences comprising the RNA probe library for the 79 genes.

**Supplementary Table 3.**
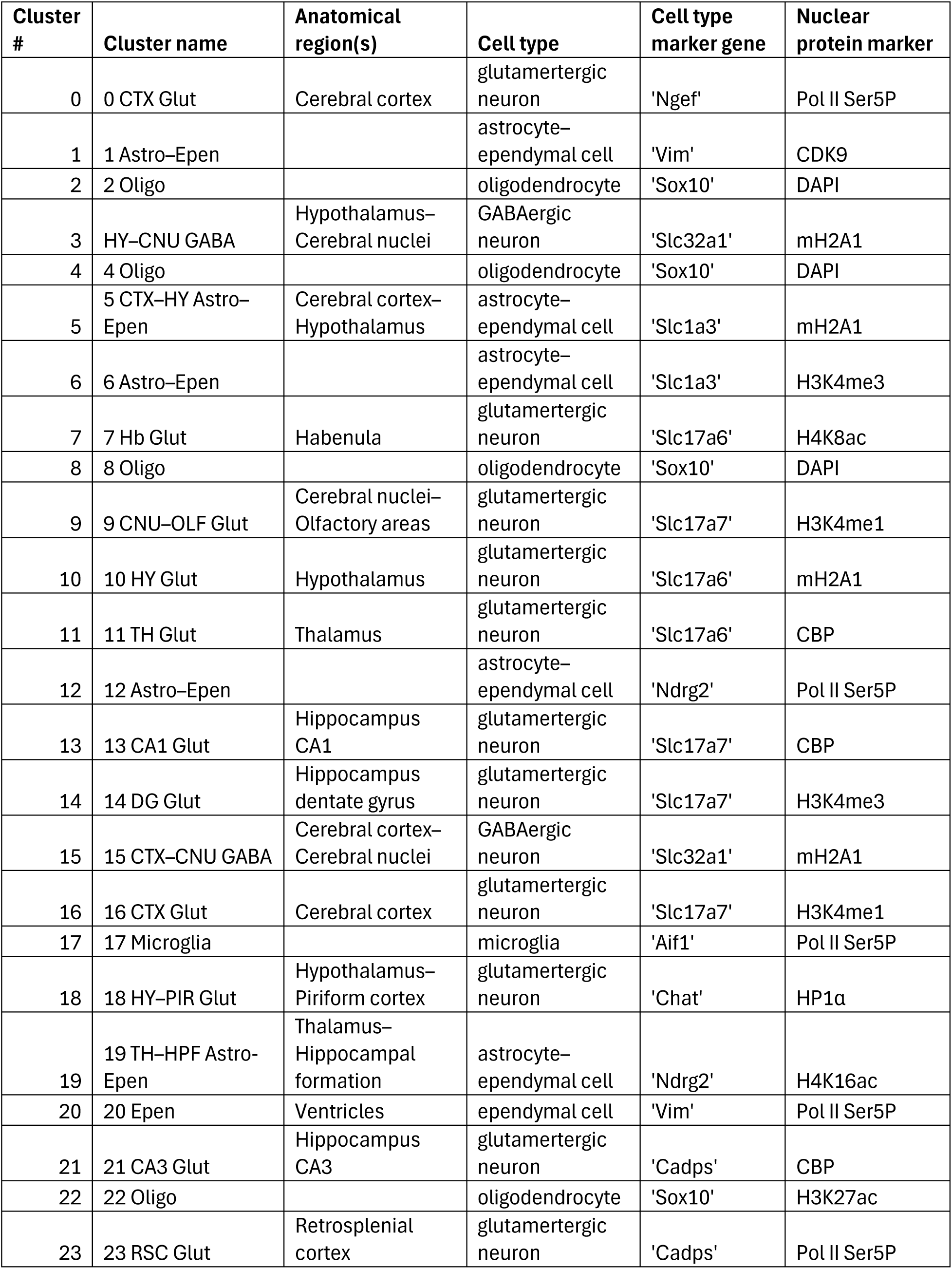
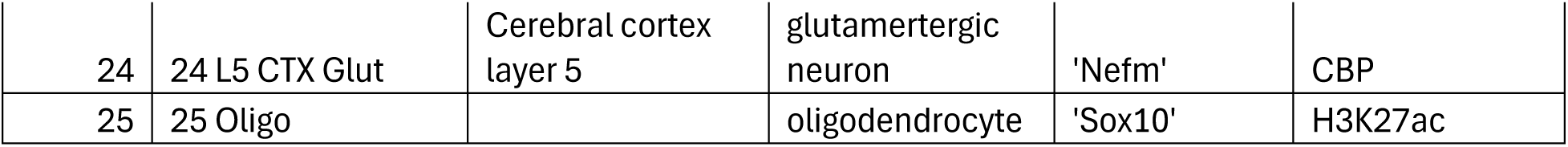
Cell clusters defined by nuclear protein intensities. Spatial regions indicate where each cluster is predominantly localized. Nuclear protein markers and cell type marker genes were identified as those with the highest log fold change compared to all other clusters. Cell types were assigned based on marker gene expression.

**Supplementary Table 4.**
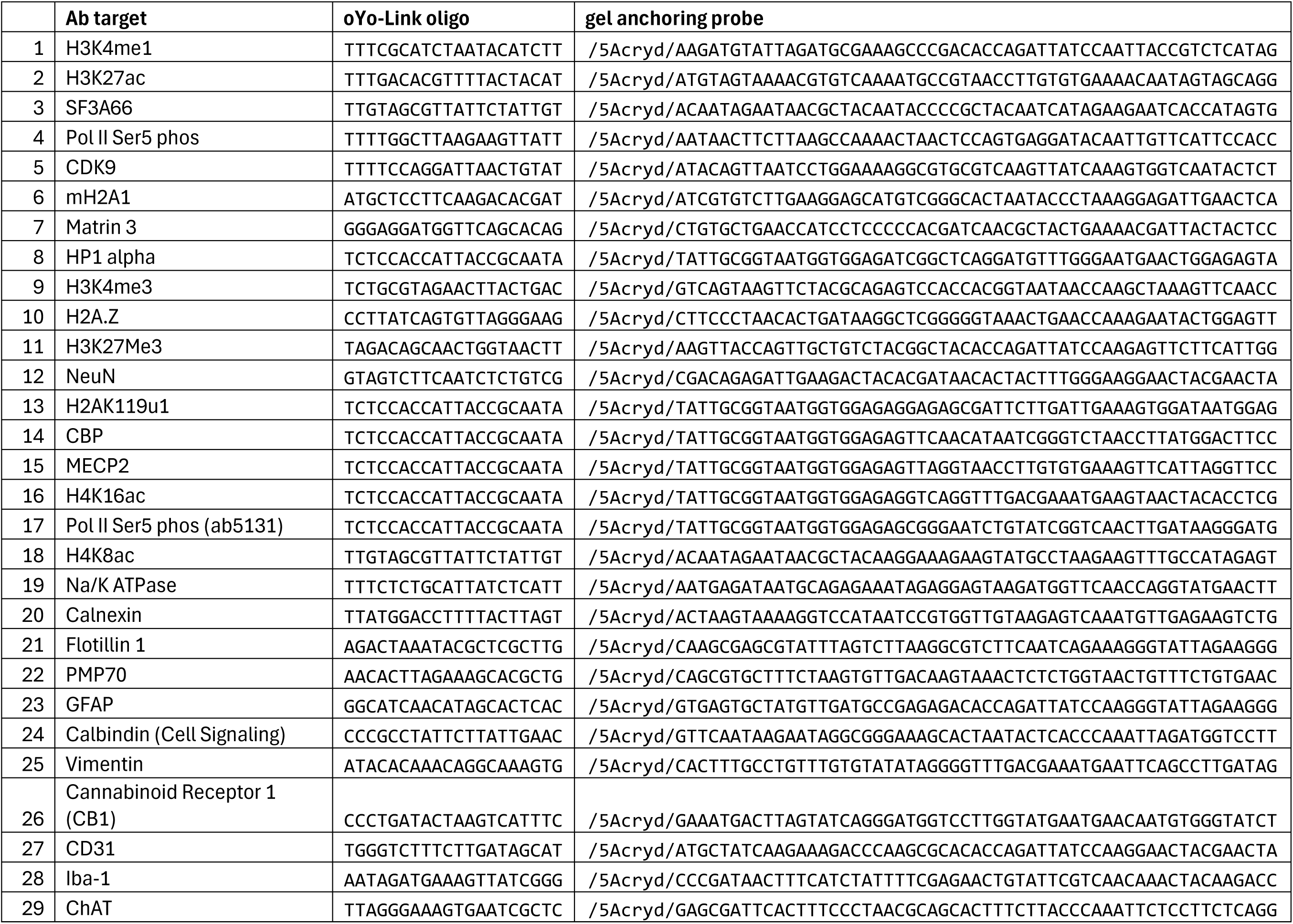
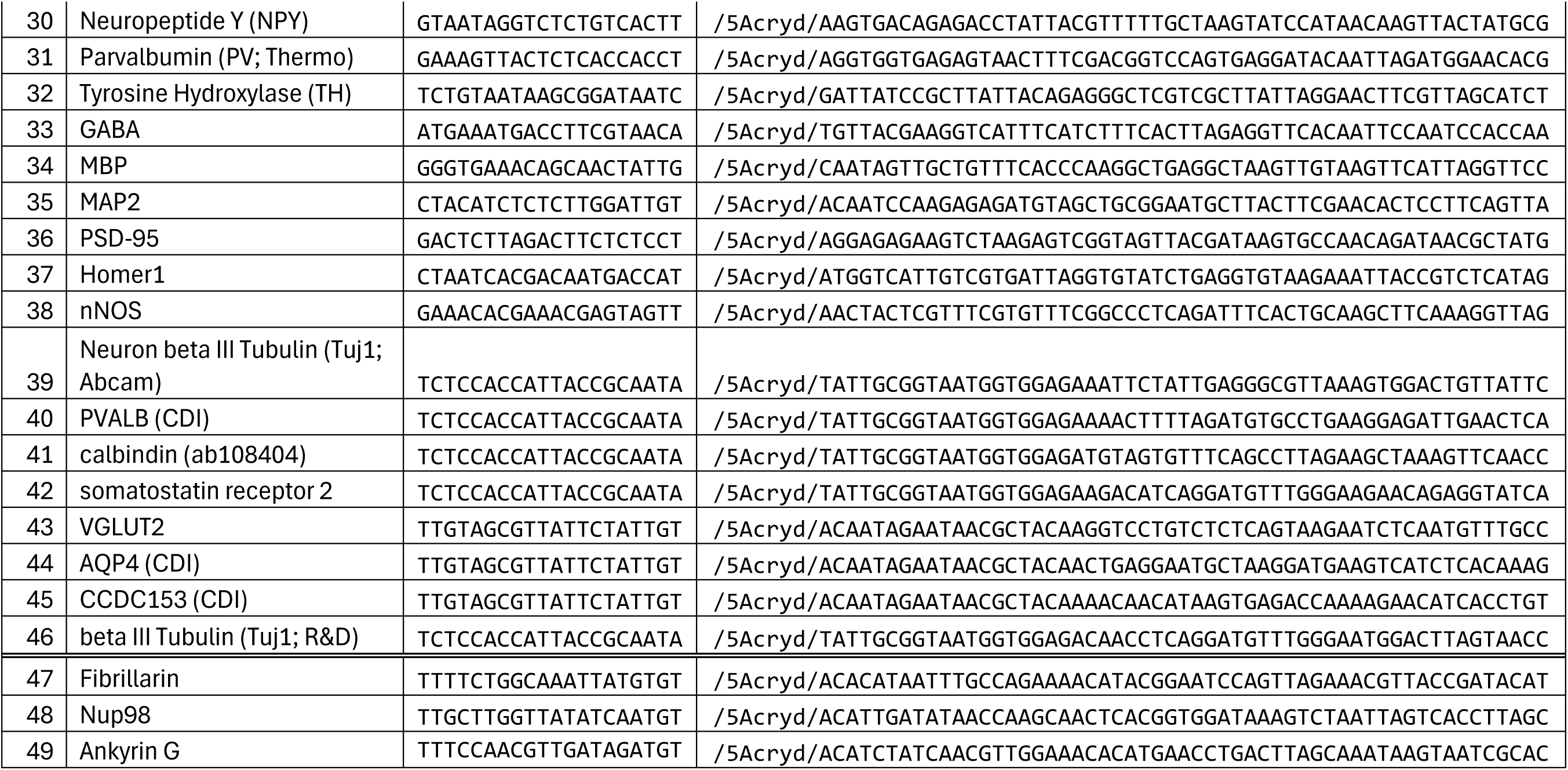
Sequences of oYo-Link DNA docking strand and gel-anchoring probe assigned to each antibody for protein cycleHCR.

